# Neuronal circuitry underlying female aggression in *Drosophila*

**DOI:** 10.1101/2020.05.27.118810

**Authors:** Catherine E. Schretter, Yoshinori Aso, Marisa Dreher, Alice A. Robie, Michael-John Dolan, Nan Chen, Masayoshi Ito, Tansy Yang, Ruchi Parekh, Gerald M. Rubin

## Abstract

Aggressive social interactions are used to compete for limited resources and are regulated by complex sensory cues and the organism’s internal state. While both sexes exhibit aggression, its neuronal underpinnings are understudied in females. Here, we describe a set of connected neurons in the adult *Drosophila melanogaster* central brain that drive female aggression. We identified a population of sexually dimorphic aIPg neurons whose optogenetic activation increased, and genetic inactivation reduced, female aggression. Analysis of GAL4 lines identified in an unbiased screen for increased female chasing behavior revealed the involvement of another sexually dimorphic neuron, pC1d, and implicated pC1d and aIPg neurons as core nodes regulating female aggression. pC1d activation increased female aggression and electron microscopy (EM) connectomic analysis demonstrated that aIPg neurons and pC1d have strong reciprocal connections. Our work reveals important regulatory components of the neuronal circuitry that underlies female aggressive social interactions and provides tools for their manipulation.

## Introduction

Aggressive behaviors are important for gaining access to resources, including food and territory, and are exhibited by both sexes in multiple species (Anderson, 2016; Kravitz and Huber, 2003; Zwarts et al., 2012). As aggressive actions carry the risk of injury, strict regulation of aggression is needed to facilitate survival. Sensory information about the presence of other individuals and the nature of the surrounding environment strongly modulate aggressive social interactions (Chen and Hong, 2018; Hoopfer, 2016). However, understanding the neuronal mechanisms by which such stimuli influence aggression has been hindered by a lack of knowledge about the structure of the underlying neuronal circuits, particularly in females.

Centers mediating, or conveying the information necessary for, aggression have been identified in the medial hypothalamus through classic experiments using electrical stimulation in cats and rodents (Albert et al., 1979; Bandler et al., 1972; Berntson, 1973; Chi and Flynn, 1971; Gregg, 2003; Kruk et al., 1983; Lammers et al., 1988; Siegel et al., 1999; Takahashi and Miczek, 2015; Woodworth, 1971). Such key regions are thought to perform a different role than other brain areas that facilitate aggressive interactions by altering the overall level of social behavior (Siegel et al., 1999). Recent work using opto-and chemo-genetic techniques have narrowed down these key regions to small populations of cells in mice, including those expressing estrogen receptor alpha (Esr1) and progesterone receptor (PR) in the ventrolateral part of the ventromedial hypothalamus (VMHvl) (Hashikawa et al., 2017; Lee et al., 2014; Yang et al., 2013). While Esr1^+^ neurons in the VMHvl regulate aggression in both male and female mice, there are sex differences in the populations involved (Hashikawa et al., 2017). Additionally, the VMHvl has also been implicated in other female sexual behaviors (Hashikawa et al., 2017; Lee et al., 2014; Pfaff and Sakuma, 1979a, 1979b; Yang et al., 2013), further complicating the identification of the specific cells that mediate aggressive interactions.

Since the first observation of aggressive behaviors in *Drosophila* by Sturtevant in 1915, social behaviors associated with attack and threat displays in flies have been well described ethologically (Shelly, 1999; Sturtevant, 1915; Ueda and Kidokoro, 2002; Zwarts et al., 2012). While male aggression is heightened in the presence of mate-related cues, female flies display increased aggressive encounters in the presence of limited nutrients, such as yeast, and when near egg laying sites (Bath et al., 2017, 2018; Lim et al., 2014; Shelly, 1999; Ueda and Kidokoro, 2002). Additionally, social isolation can increase aggression in both male and female flies (Hoffmann, 1990; Ueda and Kidokoro, 2002). As in mammals (Hashikawa et al., 2017), aggressive behaviors in flies include sex-specific components, such as head butting in female flies, as well as those that are shared between the sexes (Nilsen et al., 2004). Due to the complexity of the behavior and the sensory stimuli that influence its presentation, a circuit diagram would greatly facilitate understanding the underlying neuronal mechanisms. To gain a mechanistic understanding of how these behaviors are regulated and executed, we will need to identify the specific cells that contribute in each sex and place them in the context of larger neuronal circuits.

The fruit fly, *Drosophila melanogaster*, provides a good model for dissecting the neuronal circuitry of aggression due to the genetic tools available for targeting and manipulating individual cell types, the availability of extensive connectomic information, and the relative simplicity of its nervous system and behavior (Bellen et al., 2010; Dionne et al., 2018; Kravitz and Huber, 2003; Scheffer et al., 2020; Simpson and Looger, 2018; Tirian and Dickson, 2017). In male flies, studies investigating the neuronal correlates of aggression have implicated a group of 18 – 34 cells in the central brain, the P1/pC1 cluster, as well as various neuropeptides and biogenic amines, including neuropeptide F, tachykinin, and octopamine (Alekseyenko et al., 2019, 2014; Asahina, 2018, 2017; Asahina et al., 2014; Dierick and Greenspan, 2007; Hoopfer et al., 2015; Hoyer et al., 2008; Ishii et al., 2020; Wohl et al., 2020; Wu et al., 2020; Zhou et al., 2008). However, research on female aggressive social interactions has been less extensive as females exhibit less aggression under the behavioral conditions used for males. There are also sex differences in the behavioral components and underlying neurons important for aggression (Hoopfer et al., 2015; Nilsen et al., 2004). Genes involved in sexual differentiation, including *doublesex* (*dsx*) and *fruitless* (*fru*), contribute to sexual behaviors (Dickson, 2008; Pavlou and Goodwin, 2013; Siwicki and Kravitz, 2009; Yamamoto, 2007; Yamamoto and Koganezawa, 2013; Zhou et al., 2014). Recent work has revealed the involvement of the *doublesex*-expressing (*dsx^+^*) pC1 cluster, a group of 5 cell types, in promoting aggressive phenotypes in female flies (Fathy, 2016; Palavicino-Maggio et al., 2019). As in the VMHvl of mice, cells within this cluster can be divided into multiple subtypes and particular subtypes have been shown to also be involved in other female behaviors, including mating and egg laying (Wang et al., 2020). Understanding the flow of the information within the neuronal circuit will require knowledge of which cells within the pC1 cluster contribute preferentially or exclusively to aggressive behaviors.

Recent advances in connectomics, combined with previously established genetic tools for manipulating neuronal cell types, facilitate systematic and comprehensive studies of the circuitry contributing to behavior. We have used these methods to characterize a population of morphologically similar, sexually dimorphic, cholinergic, and *fruitless-*positive neurons, a subset of the aIP-g neurons described by Cachero et al. (2010). We found that optogenetic activation of these cells, even in the absence of aggression-promoting environmental conditions, increased female aggression. This behavior extended beyond the stimulation period, similar to the persistent phenotype generated by P1 neuronal activation in males (Hoopfer et al., 2015). Importantly, blocking aIPg synaptic transmission resulted in diminished female aggression, suggesting they modulate wild-type interactions. We then identified a second cell type, a particular pC1-subtype (pC1d), that also induces aggression upon activation. Analysis of a connectome of a large part of the fly central brain (Scheffer et al., 2020) revealed that aIPg and pC1d neurons are reciprocally and strongly connected, consistent with a coordinated role in female aggression. Taken together, our work yields insights into female aggressive behavior and identifies the two cell types that appear to form the key nodes of the circuit that implements these behaviors.

## Results

### Identification of neurons modulating female aggressive behaviors

In a behavioral screen using split-GAL4 lines to examine another phenotype, we noted a dramatic increase in head butting and fencing behaviors, known components of female aggression (Bath et al., 2017; Nilsen et al., 2004; Palavicino-Maggio et al., 2019; Ueda and Kidokoro, 2002), upon stimulation of one neuronal subset of approximately eleven cells (see Video 1 compared with Video 2). We generated multiple, independent split-GAL4 lines (Dionne et al., 2018; Luan et al., 2006; Pfeiffer et al., 2010) labelling this same neuronal population. Group-housed virgin females from these lines expressing the red-shifted opsin CsChrimson were then screened for behavioral changes upon light activation (Kim et al., 2015; Klapoetke et al., 2014). We tracked freely moving flies within a standardized 127 mm arena using video-assisted tracking software to perform automated behavioral analyses (Branson et al., 2009; Robie et al., 2017; Simon and Dickinson, 2010). These lines all exhibited similar increases in aggressive behaviors, such as fencing and head butting (Figure 1, Figure 1-figure supplements 1, 2 and 3; compare Videos 1 and 2). Consistent with the behaviors being optogenetically induced, these interactions were virtually absent when all *trans*-retinal was omitted from the food (Figure 1-figure supplement 4).

**Figure 1.**
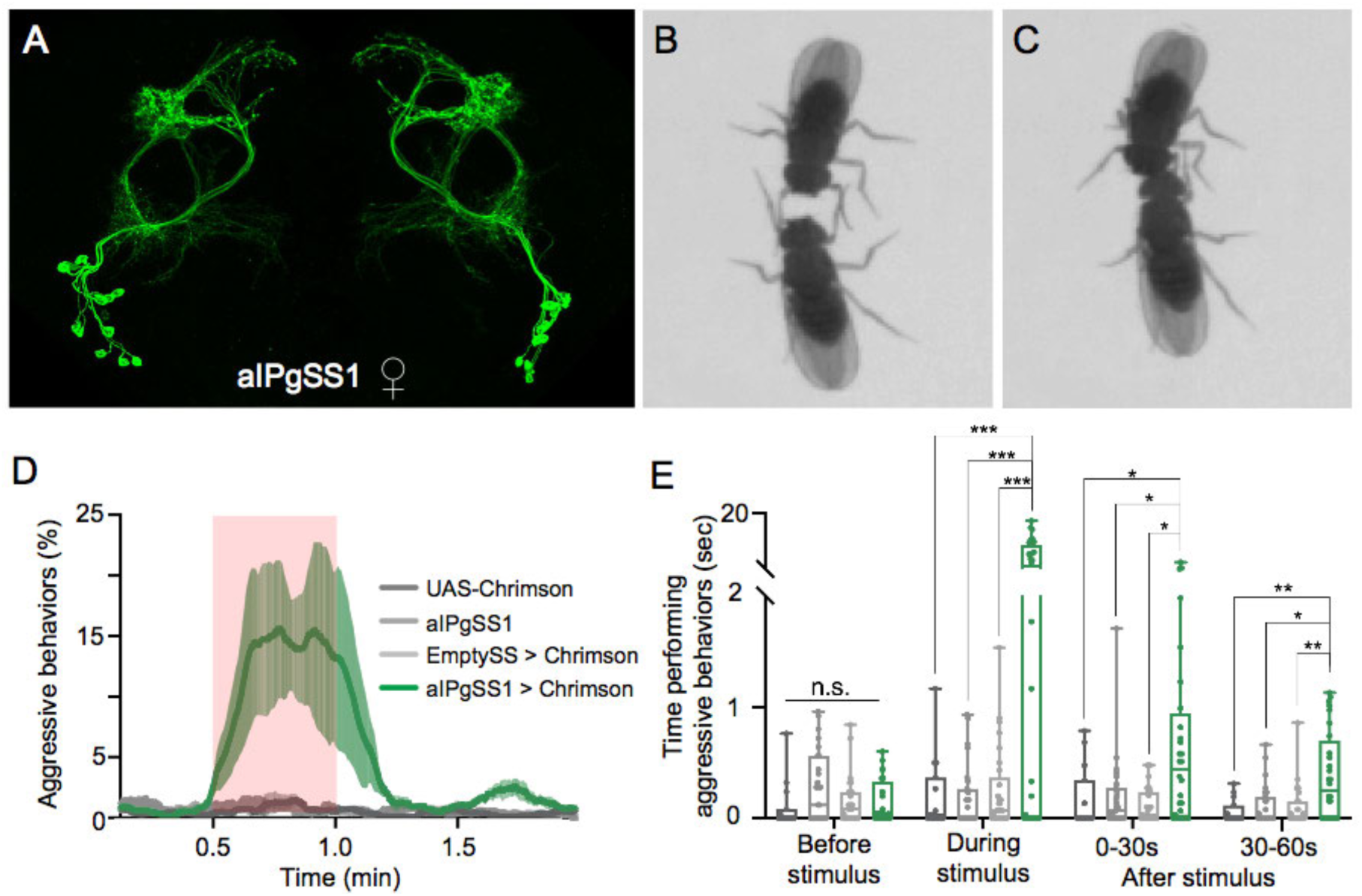
Optogenetic activation of 11 aIPg neurons results in an increase in female aggressive behaviors. (A) Maximum intensity projection (63x) image of the central brain of a female from the aIPgSS1 line crossed with 20xUAS-CsChrimson::mVenus and stained with anti-GFP antibody. Images of the complete brain and ventral nerve cord of a female and male of the same genotype are shown in Figure 1-figure supplement 1A – C. Images of individual aIPg neurons, generated by stochastic labeling are shown in Figure 1-figure supplement 2. (B, C) Images of female flies displaying fencing (B) and head butting (C) behaviors. (D) Percentage of flies engaging in aggressive behaviors, defined as displaying fencing or head butting, as measured by a JAABA classifier over the course of a 2-minute trial during which a 30-second 0.4 mW/mm^2^ continuous light stimulus was delivered. The red shading indicates the stimulus period. The mean is represented as a solid line and shaded error bars represent standard variation between experiments. Each experiment included approximately 15 flies. See Figure 1 – figure supplement 7G for additional quantification. For a description of the JAABA classifier, see Materials and Methods. 20xUAS-CsChrimson, n = 3 experiments; aIPgSS1, n = 3 experiments; EmptySS > 20xUAS-CsChrimson, n = 3 experiments; aIPgSS1 > 20xUAS-CsChrimson, n = 3 experiments. (E) Total time an individual spent performing aggressive behaviors during each of four 30-second periods: prior to, during, immediately following, and 30-60 sec after the stimulus. Points represent individual flies: 20xUAS-CsChrimson, n = 15 flies; aIPgSS1, n = 27 flies; EmptySS > 20xUAS-CsChrimson, n = 29 flies; aIPgSS1 > 20xUAS-CsChrimson, n = 28 flies. Box-and-whisker plots show median and interquartile range (IQR); whiskers show range. A two-way ANOVA with a Tukey’s multiple comparisons post-hoc test was used for statistical analysis. Asterisk indicates significance from 0: *p<0.05; **p<0.01; ***p<0.001; n.s, not significant.

To characterize these cells, we used whole mount immunohistochemistry and confocal imaging with an mVenus-tagged CsChrimson reporter. These lines labelled a set of neurons with cell bodies in the inferior protocerebrum and major projections in the anterior optic tubercle (AOTU), anterior ventrolateral protocerebrum (AVLP), superior medial protocerebrum (SMP), and superior intermediate protocerebrum (SIP) (Figure 1A, Figure 1-figure supplements 1 and 2). The cells observed in our split-GAL4 lines morphologically resemble, and appear to be a subset of, the 32 aIP-g neurons described in Cachero et al. (2010), which were previously classified as *fru*^+^ auditory interneurons with sexually dimorphic projections. We refer to this subset as aIPg. In two of our aIPg lines (aIPgSS1 and aIPgSS4), no expression was seen in males (Figure 1-figure supplement 1C). We tested for changes in male behavior in the aIPgSS1 line and found no differences (Figure 1-figure supplement 5), which was unsurprising given the absence of detectable expression of this line in males (Figure 1 – figure supplement 1). Transcript analysis of aIPg split-GAL4 lines confirmed the expression of fruitless (*fru*) as well as genes associated with use of acetylcholine as a neurotransmitter, which is consistent with previous descriptions (Cachero et al. 2010), as well as and short neuropeptide F (sNPF) (Figure 1-figure supplement 6).

### Quantification of the behavioral phenotypes upon activation of aIPg neurons

Aggressive behaviors have multiple components. We began by quantifying three such behaviors, chasing, touching, and walking, using a set of previously created and validated automatic behavior classifiers (Robie et al., 2017). We found that flies increased touching compared to the empty split-GAL4 control during a 30-second stimulation (Figure 1-figure supplement 7A – B). A low level of chasing was also detected upon activation (Figure 1-figure supplement 7C – D). Although the average walking velocity of the flies over the course of the trial did not differ compared to controls, a sharp decrease in the percent of flies walking followed stimulus onset (Figure 1-figure supplement 7E – F). Examination of behavior metrics for individual flies, or per-frame features, revealed a significant increase in the number of flies within two body lengths during stimulation; moreover, apparent orientation of flies towards one another can be easily observed, suggestive of visually directed movement (Figure 1-figure supplement 7H; compare Videos 1 and 2). Additionally, behavioral changes were both diminished and delayed if stimulation experiments were conducted in the absence of visible light (Figure 1-figure supplement 8), indicating that vision is important in the initial phase of these actions.

Touching and chasing are components of social behaviors, including courtship and aggression, in male flies (McKellar et al., 2019; Nilsen et al., 2004). While certain features of aggression are shared between males and females, there are sex-specific aspects, including head butting and the way in which behavioral patterns progress during an encounter (Nilsen et al., 2004). To examine female-specific attributes, we generated and validated a new JAABA classifier for female aggression (Supplementary Table 1). An aggressive event was defined as either an instance of fencing (Figure 1B) and/or head butting (Figure 1C), as these behaviors were not always distinguishable at the image resolution used for quantification. Examples of fencing and head butting are shown at a high resolution and frame rate in Video 3. Female aggression encompasses a range of behaviors involved in attack and threat displays; however, in this paper we used the term “aggression” in a limited way to refer to fencing and head butting behaviors. Employing this classifier, we found that a 30-second stimulation of lines labelling aIPg neurons increased the number of flies engaged in aggressive behaviors as well as the amount of time individuals spent in such activities (Figure 1D – E, Figure 1-figure supplement 7G, Videos 1 and 2). These behaviors extended beyond the activation period (Figure 1D – E), suggesting a persistent internal state.

Both the expression level of the effector and the light intensity used for optogenetic stimulation can influence behavior and, in extreme cases, be cytotoxic (unpublished observations; Kim et al., 2015). Higher levels of stimulation also increase the possibility that cells expressing the effector at levels too low for detection with confocal microscopy may contribute to the observed behavior. For these reasons, we examined the expression patterns of our split-GAL4 lines with the highest level of the effector used in our behavioral experiments and did not detect expression in other cell types or obvious toxicity in the aIPg cells (Figure 1, Figure 1-figure supplement 1). We also conducted experiments using 5xUAS-, 10xUAS-, and 20xUAS-CsChrimson constructs that are expected to produce a four-fold range of effector expression (Pfeiffer et al 2010; Figure 1-figure supplement 9A, C). Finally, we varied light intensity over a 10-fold range, 0.04, 0.1 and 0.4 mW/mm^2^ (Figure 1-figure supplement 9B). Similar effects on aggressive behavior were found under all conditions, strongly supporting the conclusion that the cell types we observe in the split-GAL4 lines are mediating the observed phenotypes.

We examined the effect of the stimulus frequency, as this can affect changes in the behavioral state (Hoopfer et al., 2015). We modified the stimulus delivery from the constant protocol used previously to a pulse stimuli. Delivery of a 5 Hz 0.1 mW/mm^2^ stimulus (5% duty cycle, or fraction of time during which the stimulus is on) over 30 seconds induced significant behavioral changes (Figure 1-figure supplement 10A – C). Changing the frequency and duty cycle (10 Hz, 0.1 mW/mm^2^ stimulus, 10% duty cycle) increased aggressive behaviors, but frequencies beyond 10 Hz did not significantly change behavior from the levels observed at 10 Hz. However, higher frequencies did increase aggression during the off period, suggesting that increasing the frequency and/or total intensity of the stimulation can increase persistence (Figure 1-figure supplement 10A – D). These experiments demonstrate that aIPg neurons promote female aggressive interactions under a range of stimulus conditions, and, at least under certain stimulus protocols, can cause a change in brain state that continues beyond the stimulus period.

### aIPg neurons mediate wild type female aggressive social interactions

The infrequent occurrence of female aggressive events (Bath et al., 2017; Shelly, 1999; Ueda and Kidokoro, 2002), at least under laboratory conditions, has made it difficult to study its neuronal correlates. To facilitate such experiments, we optimized environmental conditions to increase the level of female aggression in wild-type files. Alterations to diet and life history are known to increase female aggression in wild-type flies (Bath et al., 2017; Ueda and Kidokoro, 2002). We varied these parameters and arena size to observe interactions between pairs of flies. A 1 mm spot of yeast was also included within the arena and the diet of the flies was adjusted to restrict protein for 20 to 24 hours prior to testing (Ueda and Kidokoro, 2002). Under these conditions, we observed sufficient levels of aggression to examine the effects of inactivation of aIPg neurons with the synaptic inhibitor tetanus toxin (Sweeney et al., 1995). A significant reduction in the time spent performing aggressive behaviors, as measured using both manual and automated behavioral analyses, was observed with three different split-GAL4 lines (Figure 2, Figure 2-figure supplement 1A – B, and Figure 2-figure supplement 2A – D). Such changes did not appear to be due to decreased movement as flies exhibited similar or higher velocity compared to controls over the 30-minute trial (Figure 2-figure supplement 1C – D). These results indicate that aIPg neurons are important for modulating aggressive behaviors in wild-type females.

**Figure 2.**
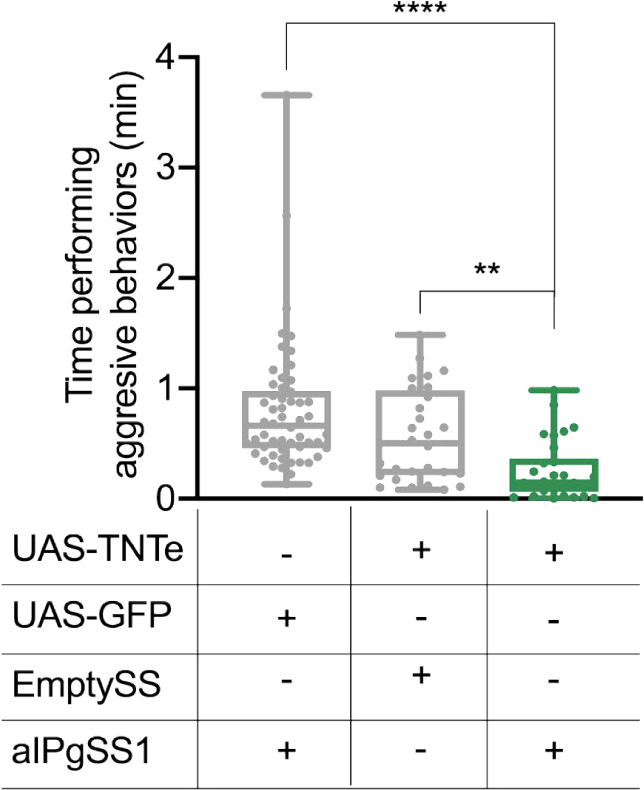
aIPg inactivation results in diminished aggressive social interactions. Total time an individual spent performing aggressive behaviors over a 30-minute trial. Individuals were pooled over 2 independent testing days during the same week. Points indicate individual flies. aIPgSS1 > UAS-GFP, n = 54 flies; EmptySS > UAS-TNTe, n = 28 flies; aIPgSS1 > UAS-TNTe, n = 30 flies. Box-and-whisker plots show median and interquartile range (IQR); whiskers show either 1.5 × IQR of the lower and upper quartiles. Kruskal–Wallis and Dunn’s post hoc tests were used for statistical analysis. Asterisk indicates significance from 0: **p<0.01; ****p<0.0001.

### Activation of aIPg overrides the requirement for specific environmental conditions for female aggressive behaviors

In addition to food availability, the genotype and sex of the target fly influence aggression (Bath et al., 2020, 2018; Lim et al., 2014; Ueda and Kidokoro, 2002). Our previous neuronal activation experiments demonstrated aggression even in the absence of competition for food (Figure 1D – E). We investigated the effects of activation status and the sex of the opponent in experiments using pairs of flies. Irrespective of whether aIPg neurons were stimulated in the both females, photostimulation in the absence of food increased the total time spent displaying aggression and decreased attack latency, or the time from stimulus on until the first aggressive event (Figure 3A – B, Figure 3-figure supplement 1 A – B, and Video 3). Females in which aIPg neurons were activated also displayed aggression when paired with wild-type males (Figure 3C, Figure 3-figure supplement 1 C, and Video 4). Mating behaviors share neuronal circuitry with aggression in males (Asahina et al., 2014; Hoopfer et al., 2015; Watanabe et al., 2017). Nevertheless, the copulation latency following stimulation of aIPg neurons did not significantly differ from that of controls (Figure 3-figure supplement 2). Taken together, our results indicate that activation of aIPg neurons can induce aggression irrespective of many environmental conditions normally associated with increased aggression in wild-type females.

**Figure 3.**
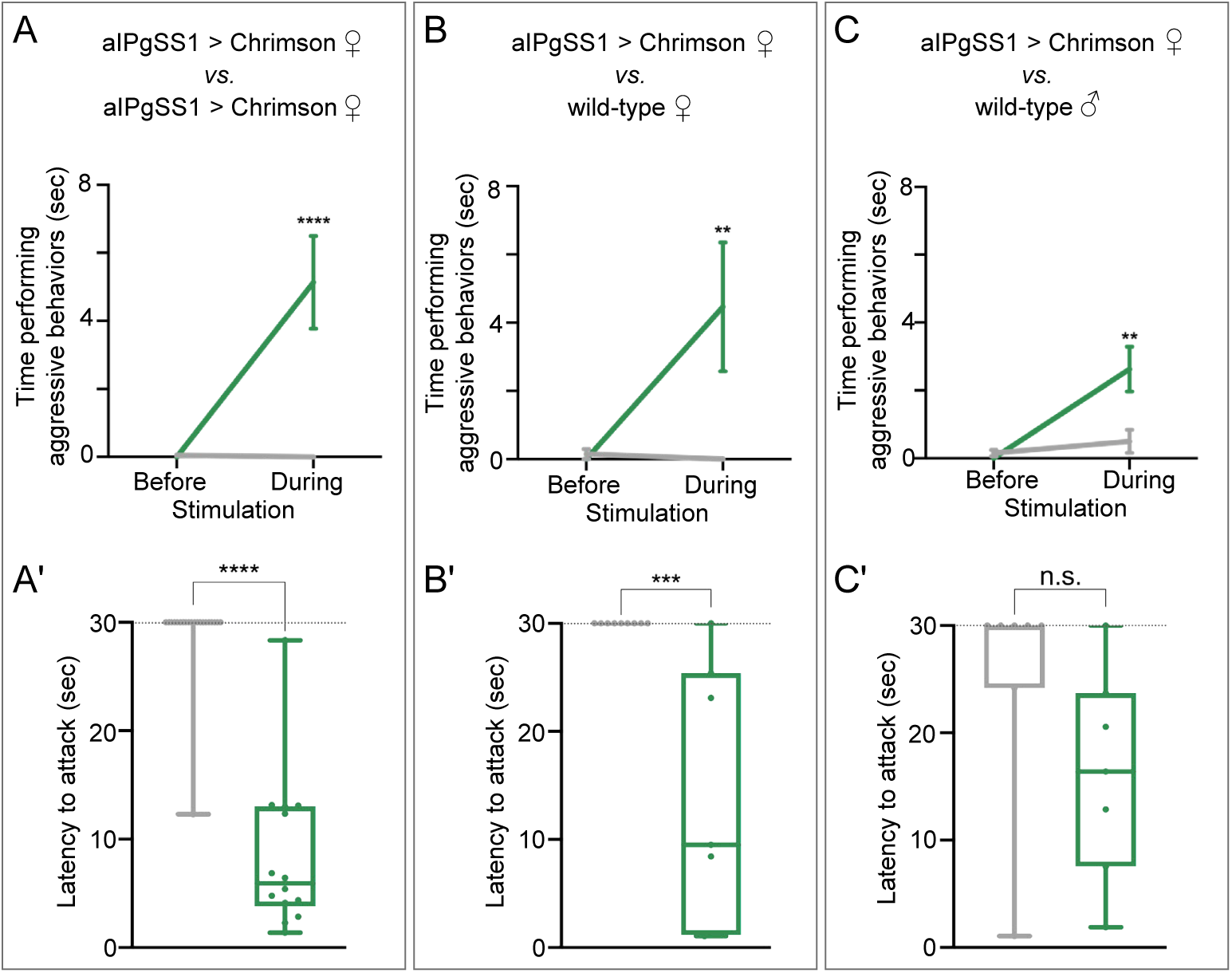
aIPg activation increases aggression against wild-type females and males. (A– C) Total time an individual spent performing aggressive behaviors in a 16 mm arena over the 30 second period prior to or during stimulation. The plots refer only to the behavior of aIPgSS1 > Chrimson females and each arena contained only two flies: (A) Two aIPgSS1 > Chrimson females; (B) an aIPgSS1 > Chrimson female and a wild-type (Canton-S) female; and (C) a aIPgSS1 > Chrimson female and a wild-type (Canton-S) male. The green line shows the stated genotype; the gray line shows the results when EmptySS > Chrimson was used instead of aIPgSS1 > Chrimson. (A’ – C’) Amount of time during a 30 second 0.1 mW/mm^2^ continuous stimulation period until first aggressive encounter. Points indicate individual flies. Dotted lines indicate the end of the trial and error bars are mean +/- S.E.M. Box-and-whisker plots show median and interquartile range (IQR); whiskers show either 1.5 × IQR of the lower and upper quartiles. A’: EmptySS > 20xUAS-Chrimson, n = 22 flies; aIPgSS1 > 20xUAS-Chrimson, n = 14 flies; B’: EmptySS > 20xUAS-Chrimson, n = 8 flies; aIPgSS1 > 20xUAS-Chrimson, n = 7 flies; C’: EmptySS > 20xUAS-Chrimson, n = 7 flies; aIPgSS1 > 20xUAS-Chrimson, n = 7 flies. A Mann-Whitney *U* (A’ – C’) post hoc test or a two-way ANOVA with a Sidak’s multiple comparisons post-hoc test (A – C) was used for statistical analysis. Asterisk indicates significance from 0: *p<0.05; **p<0.01; ****p<0.0001; n.s., not significant.

### Identifying additional cell types involved in mediating female aggression

Having established a role for aIPg neurons in female aggression, we used two complementary methods to discover additional cells involved in regulating this behavior. First, we used behavioral screens to identify other cell types that could drive female aggression when activated. Second, we used the aIPg neurons as an entry point for EM-based circuit mapping. As described below, both approaches converged on the same set of cells.

### aIPg and pC1d are two key groups of neurons involved in female aggressive behaviors

It is reasonable to expect that other neurons in the circuit or parallel pathways could also induce aggression upon activation. To identify such neurons, we took a strategy analogous to that used by geneticists to answer the question of how many distinct genes produce a particular phenotype when mutated. In their experimental approach, individual mutations are placed into complementation groups after performing a genetic screen large enough to sample all the genes. In this way, the number of different genes that can give rise to the phenotype under study when mutated can be estimated (see for example, Nüsslein-Volhard and Wieschaus, 1980). To identify further cell types that could also increase female aggression, we analyzed the results of a previous screen of over 2,000 GAL4 lines, in which cell types are expressed in multiple lines (Robie et al., 2017). We generated split-GAL4 hemi-driver lines using the enhancers from this screen’s top hits for increased female-female chasing behavior and crossed them to each other to reveal the presence of shared cell types (Figure 4; Figure 4-figure supplement 1). Strikingly, 13 of the top 14 hits based upon their behavioral score could be accounted for by just two cell types: aIPg and pC1d, one of the five cell types found in the pC1 cluster of cells in females. Members of this pC1 cluster are involved in egg laying and copulation (Wang et al., 2020). While such screens could miss cell types with less penetrant activation phenotypes and would be insensitive to cell types that functioned to inhibit aggression, these results imply a central role for aIPg and pC1d in female aggression.

**Figure 4.**
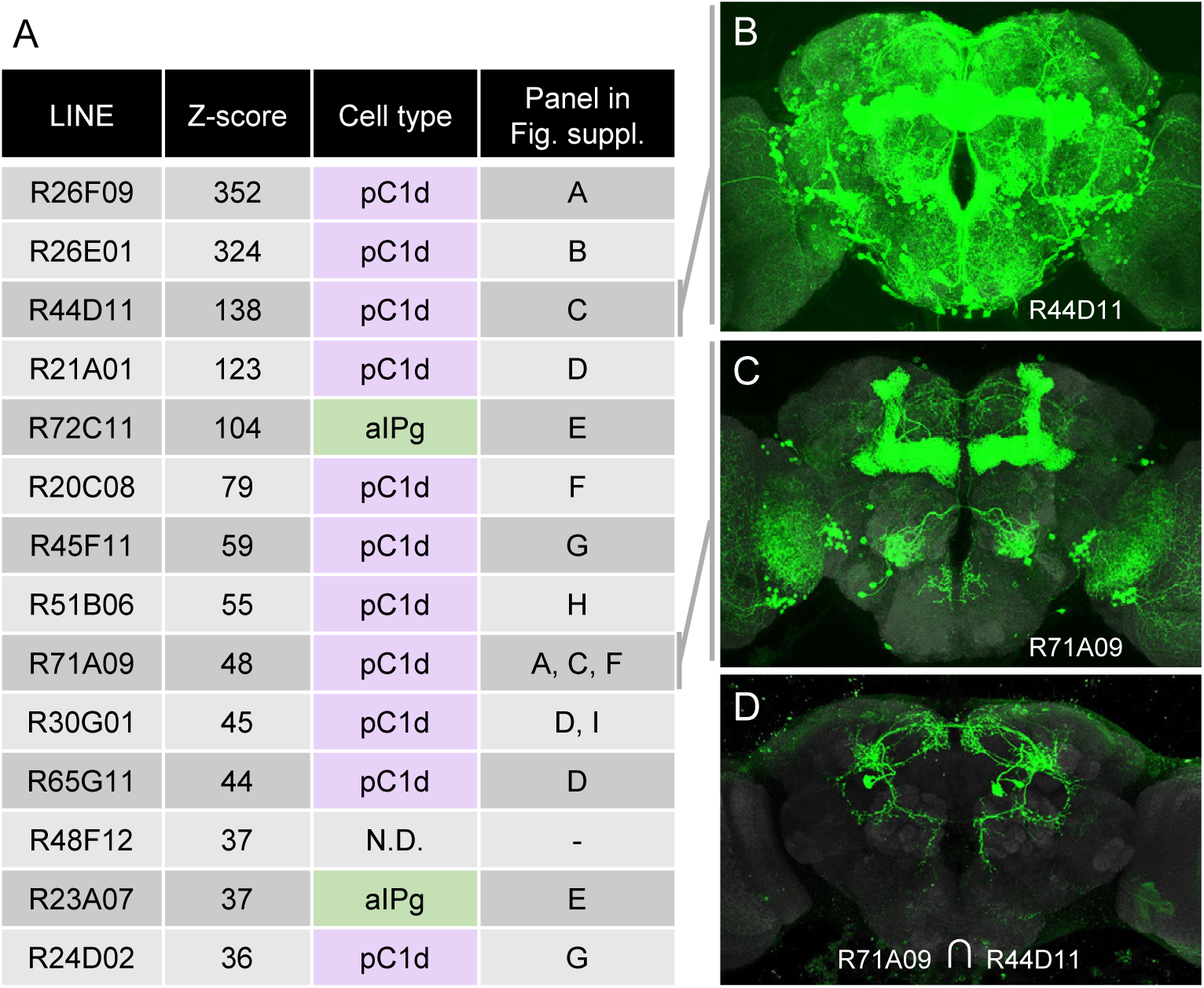
An unbiased screen suggests that aIPg1 – 3 and pC1d are key nodes for gating female-female aggression. (A) The fourteen top hits for female-female chasing from an unbiased activation screen of 2,204 generation 1 GAL4 lines are listed along with their Z-scores for this behavior as determined by Robie et al. (2017). Also shown is the relevant cell type we concluded from our intersectional analysis (Figure 4-figure supplement 1) to be present in each line. N.D., no cell type reproducibly detected, suggesting that R48F12 may not share a common cell type with any of the other 13 lines. The final column in the table refers to the panels in Figure 4-figure supplement 1 where results supporting the stated conclusion are shown. (B, C) The expression patterns of the two indicated GAL4 lines (Jenett et al. 2012). The images shown were taken from the database at www.janelia.org/gal4-gen1, where the expression patterns of the other lines listed in A can also be found. (D) The expression pattern of a split-GAL4 line made by intersecting these two enhancers; a cell with the morphology of pC1d can be seen.

The five cell types that make up the pC1 group express *dsx* and have been collectively implicated in female receptivity, oviposition, male courtship, and both male and female aggression (Fathy, 2016; Hoopfer et al., 2015; Ishii et al., 2020; Palavicino-Maggio et al., 2019; Rideout et al., 2010; Scheffer et al., 2020; Wang et al., 2020; Wohl et al., 2020; Zhou et al., 2014). In previous work, it has proven difficult to elucidate the relative contribution of the five different pC1 cell types present in females to these behaviors. pC1d cells are revealed in intersections using the enhancers from 11 of the top 14 hits from the Robie et al. (2017) screen (Figure 4; Figure 4-figure supplement 1). However, nearly all of these intersections contain at least one other pC1 cell type, leaving open the possibility that inducing aggression requires a combination of multiple pC1 cell types.

### Activation of pC1d alone, but not pC1e or pC1a – c, promotes female aggressive behaviors

To address the role of individual pC1 cell types in female aggression, we generated split-GAL4 lines that drive expression in either pC1d or pC1e as well as lines containing both cell types (Figure 5A, Figure 6A, Figure 5-figure supplements 1 and 2, and Figure 6-figure supplement 1 and 2). We also used a split-GAL4 line that labels pC1a – c (provided by K. Wang and B. Dickson). No expression was observed in males in the majority of the lines used (Figure 5-figure supplement 1C and Figure 6-figure supplement 1C). The cells labeled in males in two of the lines were not morphologically similar to pC1d or pC1e and these lines were not used for behavioral analysis in males. Transcriptional profiling of pC1d and pC1e confirmed that the cells in females expressed *dsx* and genes associated with use of acetylcholine (Figure 1-figure supplement 5), as previously described (Palavicino-Maggio et al., 2019; Rezával et al., 2016; Rideout et al., 2010; Zhou et al., 2014). Our data suggest that these cells might also express *fru* (Figure 1-figure supplement 5).

**Figure 5.**
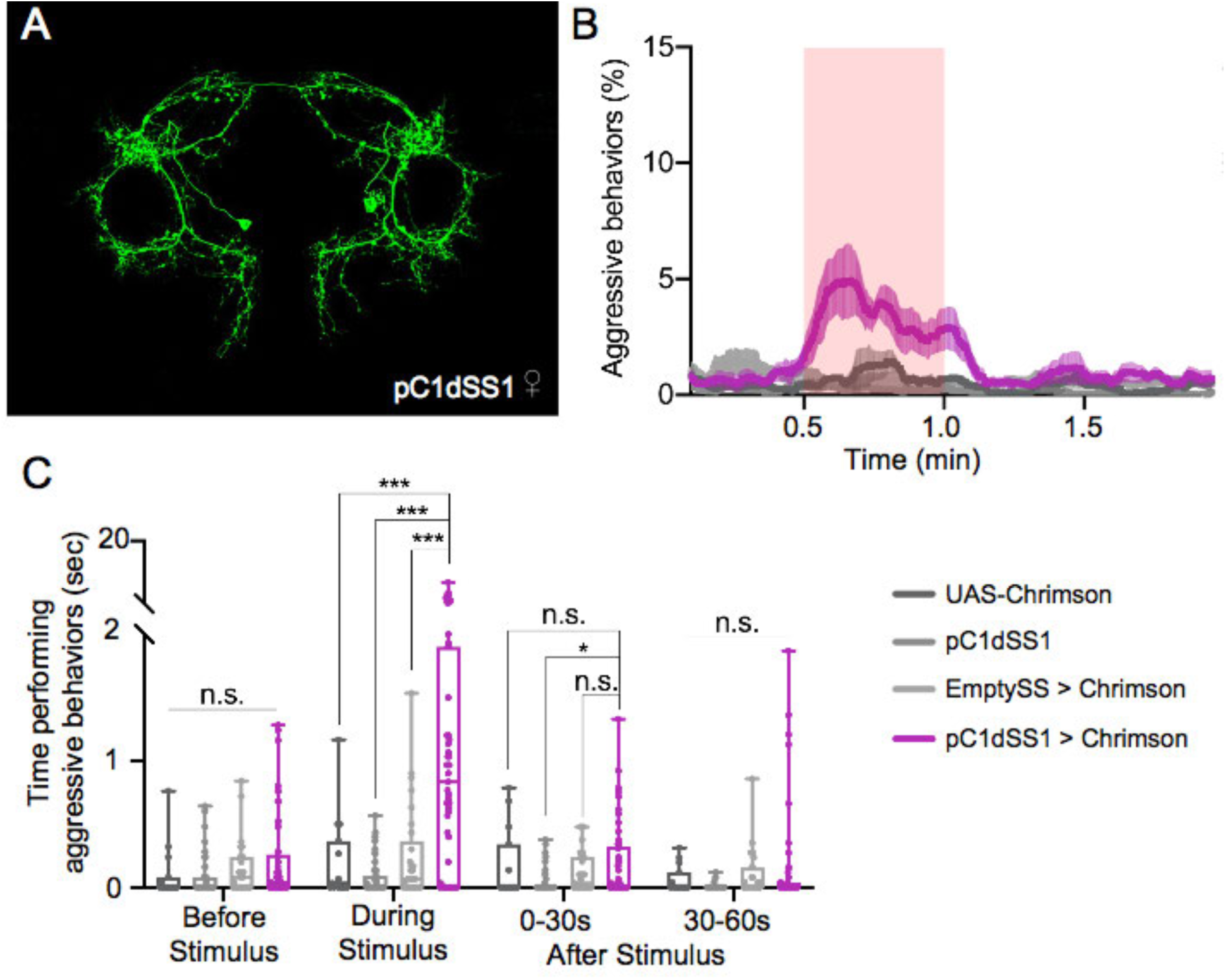
pC1d significantly increases aggressive social interactions in female flies. (A) Maximum intensity projection (63x) image of the central brain of a female from the pC1dSS1 split-GAL4 line crossed with 20xUAS-CsChrimson::mVenus and stained with anti-GFP antibody. Images of the complete brain and ventral nerve cord of a female and male of the same genotype are shown in Figure 5-figure supplement 1A – C. (B) Percentage of flies engaging in aggressive behaviors over the course of a 2-minute trial during which a 30-second 0.4 mW/mm^2^ continuous light stimulus was delivered. Mean is represented as a solid line and shaded error bars represent variation between experiments. Each experiment included approximately 15 flies. 20xUAS-CsChrimson, n = 3 experiments; pC1dSS1, n = 3 experiments; EmptySS > 20xUAS-CsChrimson, n = 5 experiments; pC1dSS1 > 20xUAS-CsChrimson, n = 5 experiments. (C) Total time an individual spent performing aggressive behaviors during each of four 30-second periods: prior to, during, immediately following, and 30-60 sec after the stimulus. Points represent individual flies. 20xUAS-CsChrimson, n = 15 flies; pC1dSS1, n = 48 flies; EmptySS > 20xUAS-CsChrimson, n = 29 flies; pC1dSS1 > 20xUAS-CsChrimson, n = 53 flies. Box-and-whisker plots show median and interquartile range (IQR); whiskers show range. A two-way ANOVA with a Tukey’s multiple comparisons post-hoc test was used for statistical analysis. Asterisk indicates significance from 0: *p<0.05; ***p<0.001; n.s., not significant.

**Figure 6.**
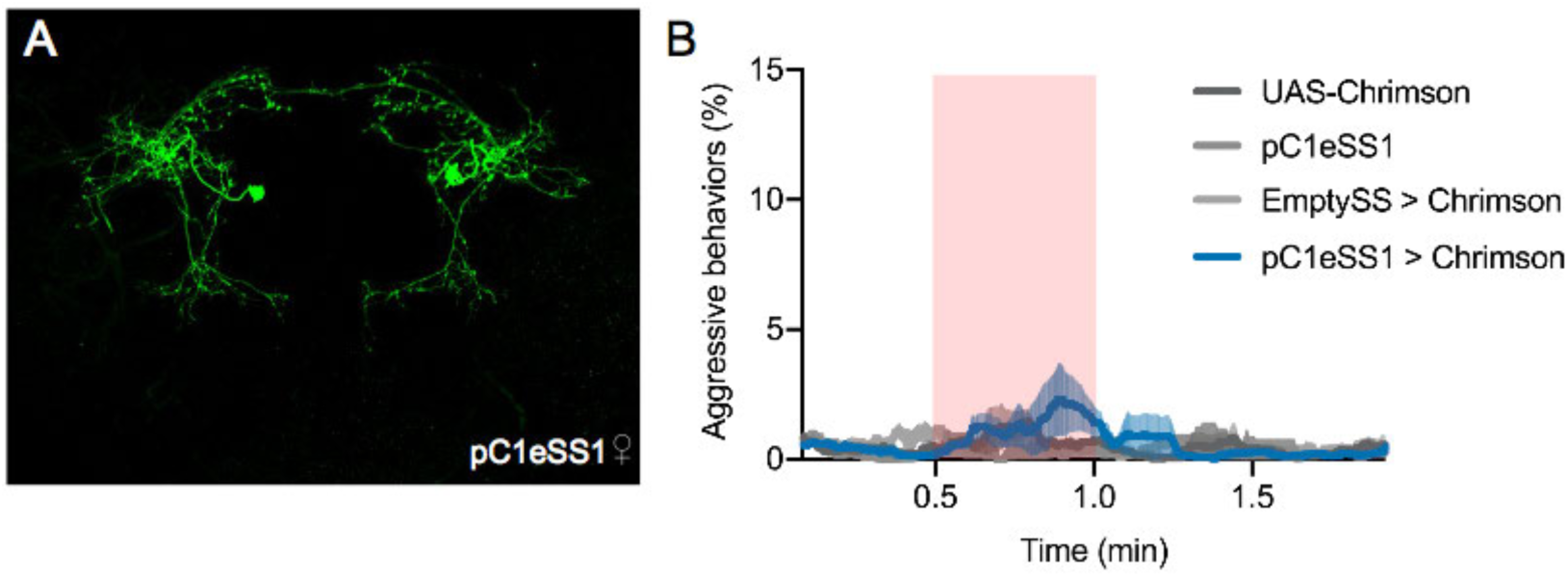
pC1e does not significantly increases aggressive social interactions in female flies. (A) Maximum intensity projection (63x) image of the central brain of a female from the pC1eSS1 line crossed with 20xUAS-CsChrimson::mVenus and stained with anti-GFP antibody. Images of the complete brain and ventral nerve cord of a female and male of the same genotype are shown in Figure 6-figure supplement 1A-C. (B) Percentage of flies engaging in aggressive behaviors over the course of a 2-minute trial during which a 30-second 0.4 mW/mm^2^ continuous light stimulus was delivered. Red shading indicates the stimulus period. The mean is represented as a solid line and shaded error bars represent variation between experiments. Each experiment included approximately 15 flies. 20xUAS-CsChrimson, n = 3 experiments; pC1eSS1, n = 2 experiments; EmptySS > 20xUAS-CsChrimson, n = 3 experiments; pC1eSS1 > 20xUAS-CsChrimson, n = 3 experiments.

Using the same stimulation parameters as we used for lines labeling aIPg neurons, optogenetic activation of pC1d alone increased the percentage of aggressive flies (Figure 5B, Figure 5-figure supplement 3A, Figure 5-figure supplement 4, and Video 5). These aggressive interactions included fencing and head butting as observed upon aIPg activation (Video 6). pC1d stimulation increased touching and chasing; however, there were no sharp decreases in walking (Figure 5-figure supplement 3B – D). Similar results were seen with additional split-GAL4 lines labelling pC1d (Figure 5-figure supplement 4), while behavioral changes were not observed upon stimulation of males (Figure 5-figure supplement 5A – B). Additional controls in which all *trans*-retinal was omitted from the food did not show an elevation in aggression following light onset (Figure 5-figure supplement 6A – B). Increasing the intensity and frequency of stimulation did heighten behavior (Figure 5-figure supplement 7A, Figure 5-figure supplement 8A – C). However, changes in the expression level of CsChrimson were, unexpectedly, inversely correlated with behavior (Figure 5-figure supplement 7B – C), suggesting that higher effector expression levels might be detrimental to cell function.

Notably, we did not find significant continued changes in behavior following the stimulation period for any of the frequencies tested using pC1d, in contrast to the persistence of aggression observed after similar activation of aIPg neurons (Figure 1-figure supplement 10A, B, D and Figure 5-figure supplement 8A, B, D). As the behaviors induced by activation of aIPg neurons were largely independent of external conditions, we tested whether pC1d behavioral effects were similarly resistant to changes in the target or arena. pC1d stimulation resulted in similar attack latency and time spent performing aggressive behaviors irrespective of whether the opponent was a wild-type female or male (Figure 5-figure supplement 9A – C). While activation promoted aggression, no differences were observed following pC1d inactivation with tetanus toxin (Figure 5-figure supplement 10). These results suggest that pC1d neurons are not essential for female aggression; however, we have not independently confirmed the degree of effectiveness of the tetanus toxin inactivation.

In contrast to pC1d, stimulation of lines containing pC1e alone did not significantly alter any of the behaviors we assayed (Figure 6B, Figure 6-figure supplement 1A – E, Figure 6-figure supplement 2A – D, Figure 6-figure supplement 3A – B). Changes in the effector expression levels, light power density, and frequency did not significantly alter behavior upon activation of pC1e (Figure 6-figure supplement 4A – D), suggesting that the absence of behavioral changes was not due to the stimulation parameters. Analysis of lines containing both pC1d and pC1e exhibited similar levels of behavior to containing pC1d alone, implying that pC1d and pC1e do not act synergistically (Figure 5-figure supplement 4A – B). Likewise, activation of pC1a – c did not change the percentage of flies displaying aggression (Figure 6-figure supplement 5A – C). Taken together, our results suggest that pC1d, but not pC1e or pC1a – c, acts as a significant facilitator of female aggression.

### aIPg and pC1d neurons form reciprocal connections and only share a few overlapping pre- or post-synaptic connections

The generation of the full adult female brain (FAFB) electron microscopic (EM) image set (Zheng et al., 2018) and the connectome of the hemibrain (Scheffer et al., 2020) allowed us to use EM level connectomics to determine the structure of the circuit(s) that contained the cells identified through our behavioral studies. First, we identified the cells in EM volumes that correspond to those observed in our aIPg split-GAL4 lines. We began this work in FAFB, before the availability of the hemibrain dataset. First, the fiber bundle in each hemisphere that contained the neurons corresponding to the ∼32 cells of the aIP-g lineage based on cell body and soma tract position in the brain (Cachero et al., 2010) was identified. We sufficiently traced the axon and major dendritic arbors of all these cells to determine if they matched the morphologies of the neurons observed in our split-GAL4 lines (Figure 7). While the hemibrain connectome was being generated, we were able to identify the corresponding aIPg cells (Figure 7 – figure supplement 1) as well as those of the pC1 cluster, allowing us to participate in the effort to improve the accuracy of their reconstruction and analyze their morphology in greater detail. The results of these reconstructions are in the v1.0 connectome (neuprint.janelia.org) reported in Scheffer et al. (2020). Based on morphological differences, most notably in their SMP and SIP arbors, and later confirmed by their distinct connectivity, we separated the aIPg neurons into 3 distinct types, aIPg type 1, aIPg type 2, and aIPg type 3 (Figure 8 and Videos 7 and 8). Light microscopy images confirmed the presence of these aIPg types within our split-GAL4 lines (Figure 1-figure supplement 2).

**Figure 7.**
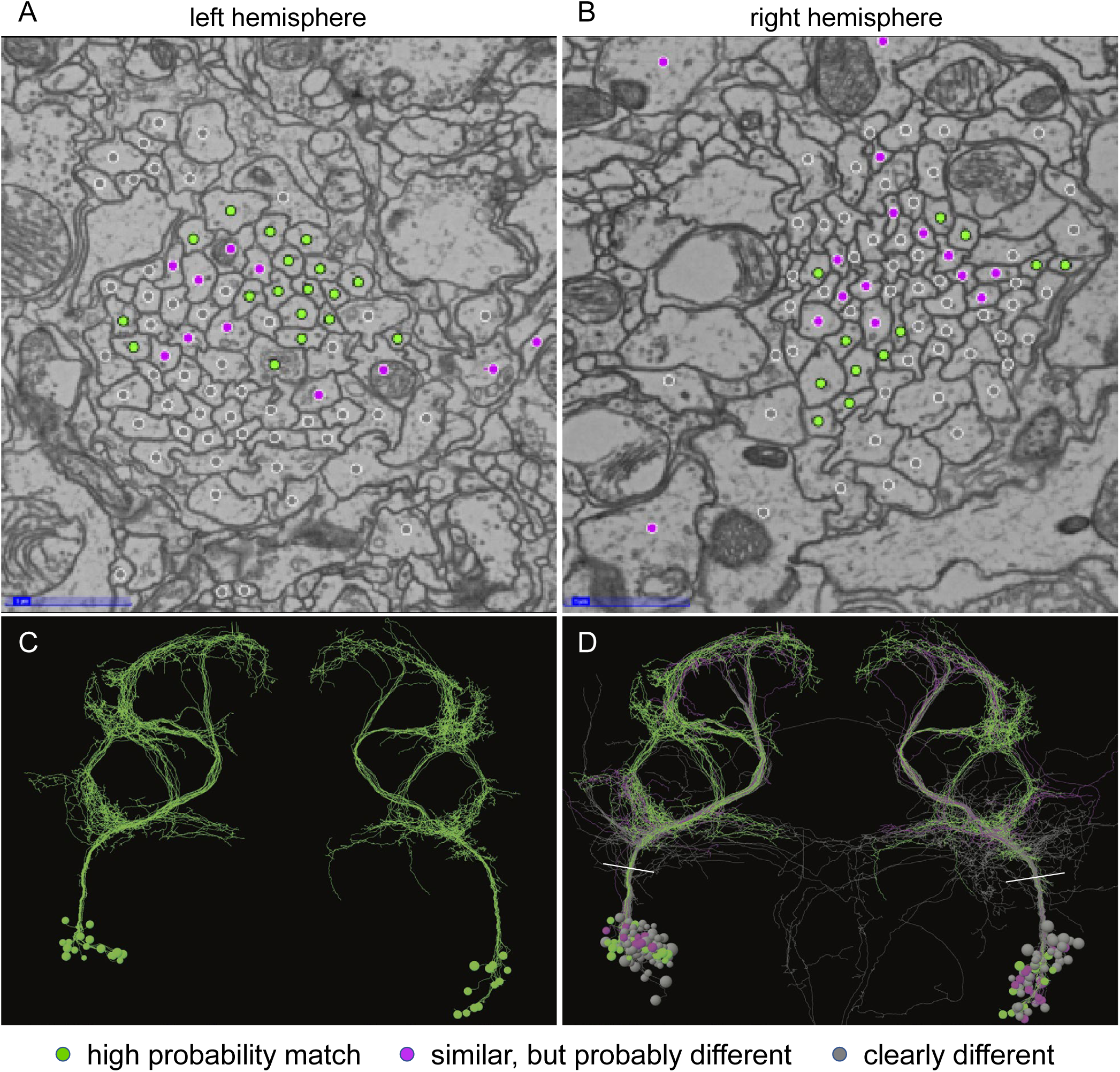
Identification of aIPg neurons in the FAFB dataset. (A – B) Images of an area of the left and right hemisphere of an EM section from the FAFB dataset containing the fiber tracts of the 32 putative aIP-g neurons described by Cachero et al. (2010). A dot has been placed in each axon, color-coded to reflect the degree of similarity of its morphology, revealed by manual tracing and visual inspection, to the aIPg neurons contained in our split-GAL4 lines. Grey represents neurons whose morphology clearly differed, magenta represents neurons whose morphology were similar but differed in one or more branches, and green represents neurons that we judged to correspond to those in our split-GAL4 lines. Note that, as we often observe in our split-GAL4 lines (see Figure 1-figure supplement 1), the number of neurons differed between hemispheres; in this case, the left hemisphere had 17 green cells, while the right hemisphere had only 12. (C – D) Skeleton rendering of the traced aIPg neurons, colored based on their similarity to the aIPg neurons identified in our split-GAL4 lines. Panel C shows only cells we judged to correspond to those in our split-GAL4 lines, while D shows all traced cells. White lines in D indicate the approximate plane of A – B. Tracing of grey neurons was stopped when it was clear they did not match our split-GAL4 lines; therefore, their arbors are likely to be incomplete in these images.

**Figure 8.**
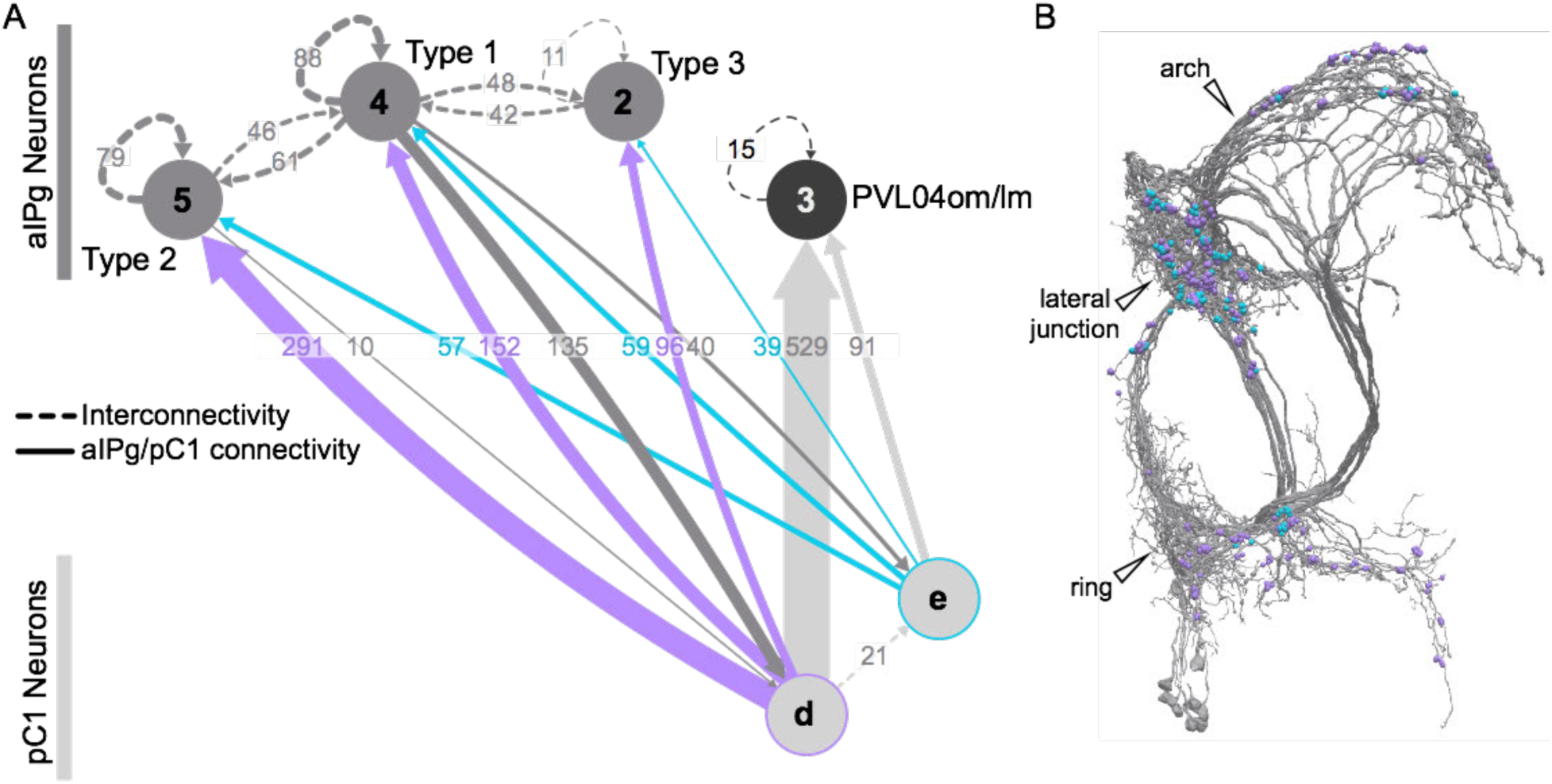
Reciprocal interactions between aIPg and pC1 neurons identified by EM connectomics. (A) Interconnectivity between pC1d, pC1e, aIPg Type 1, aIPg Type 2, aIPg Type 3, and PVL04om/lm neurons thresholded at 10 synapses. PVL04om/lm neurons also derive from the aIP-g lineage. Synapse number is noted on each arrow and dashed lines represent interconnectivity within the aIPg Type 1, Type 2 or Type 3, PVL04om/lm or pC1 neurons. Number within circle represents the number of neurons within the cell type; circles without numbers have only one neuron per brain hemisphere. Colored arrows depict synapses from pC1d (purple) and pC1e (blue) to aIPg types. (B) Electron microscopy rendering of all 11 Type 1, 2 and 3 aIPg neurons seen in the hemibrain with color-coded dots at the sites of synapses from pC1d (purple) and pC1e (blue); these synapses are concentrated in the lateral junction above the peduncle of the mushroom body. See Video 7 for more detail on interconnectivity between aIPg neuron types and Video 8 for detail on the connections between aIPg and pC1 neurons.

From analyzing the connectivity of aIPg neurons, we identified two pC1 cell types, pC1d and pC1e, among their top six presynaptic inputs. We are confident that these pC1 cells correspond to those observed in our split-GAL4 lines (Figure 5-figure supplement 2, Figure 6-figure supplement 2) as no other cells with similar morphology were found in the hemibrain volume. These light to EM level assignments are also consistent with those reported by Wang et al (2020) for light microscopy-FAFB correspondences. No connections above the threshold of 10 synapses were identified in the hemibrain between aIPg neurons and the other three cell types in the pC1 cluster, pC1a – c (Figure 8-figure supplement 1).

Another group of cells in the hemibrain volume, PVL04om/lm, is also heavily innervated by pC1d (Figure 8A). These cells bear a morphological resemblance to aIPg neurons, but they have distinct projections in the SMP. Moreover, PVL04om/lm neurons do not directly connect with the aIPg type 1, type 2, or type 3 neurons and two of their top inputs include pC1a and pC1c, which do not provide input to the aIPg neurons (Figure 8A and Figure 8-figure supplement 1). Analysis of our split-GAL4 lines indicates that PVL04om/lm cells are unlikely to be consistently present, especially in split-GAL4 lines, such as aIPgSS3, that have expression in a smaller number of cells. Consistent with this interpretation, we have not observed PVL04om/lm cells in our MultiColor FlpOut (MCFO) analyses of the aIPg lines. Their similar morphology combined with the inherent variability of the cell number in our split-GAL4 lines, even between the brain hemispheres of a single individual, means that we cannot formally rule out the possibility of PVL04om/lm cells contributing to activation phenotypes seen with our aIPg split-GAL4 lines. Nevertheless, PVL04om/lm neurons were not included in further analysis and examination of the role of these cells awaits split-GAL4 lines that specifically express in them.

Type 1 aIPg neurons provide strong reciprocal feedback onto pC1d (Figure 8A – B and Video 9), but types 2 and 3 do not. There are also extensive reciprocal connections between aIPg neurons of types 1 and 2 and between types 1 and 3, but not between types 2 and 3 (Figure 8A and Video 8).

Among the aIPg neurons’ six strongest pre-synaptic inputs, as judged by synapse number, we found that pC1d and pC1e rank first and sixth, respectively (Figure 9 and Video 10). A neuron previously implicated in oviposition, oviIN (Wang et al. 2020), ranked eleventh among aIPg inputs, based on synapses number (Figure 9 and Supplementary Table 2). The second and third strongest inputs, SCB014 and the ADM07t, also provide presynaptic input to pC1d, although it is a lower percentage of their total output at 1.89% from SCB014 and 0.22% from the ADM07t group to pC1d, compared to 7.7% and 8.6% to the aIPg neurons, respectively (Figure 9A, Video 10, Supplementary Table 2 and 3). OviIN also forms connections with both aIPg and pC1d, but aside from these three cell types aIPg and pC1d do not share any strong pre-synaptic inputs (Figure 9A – B, 10A – B, Video 10 and 12; note that ADM07t falls below the cutoff for being displayed and for being considered a significant connection in Figure 10A and in Supplementary Table 3). Only one of the top downstream targets of the aIPg neurons, ADM01r, is also a major downstream target of pC1d (Figure 9C – D and 10C – D, Video 11 and 13, Supplemental Table 2 and 3). However, again we found that the connectivity strengths differ considerably: 1.13% of ADM01r’s input comes from pC1d, while 14.42% is provided by aIPg neurons (Figure 9C and 10C, Video 11, Supplemental Table 2 – 3).

**Figure 9.**
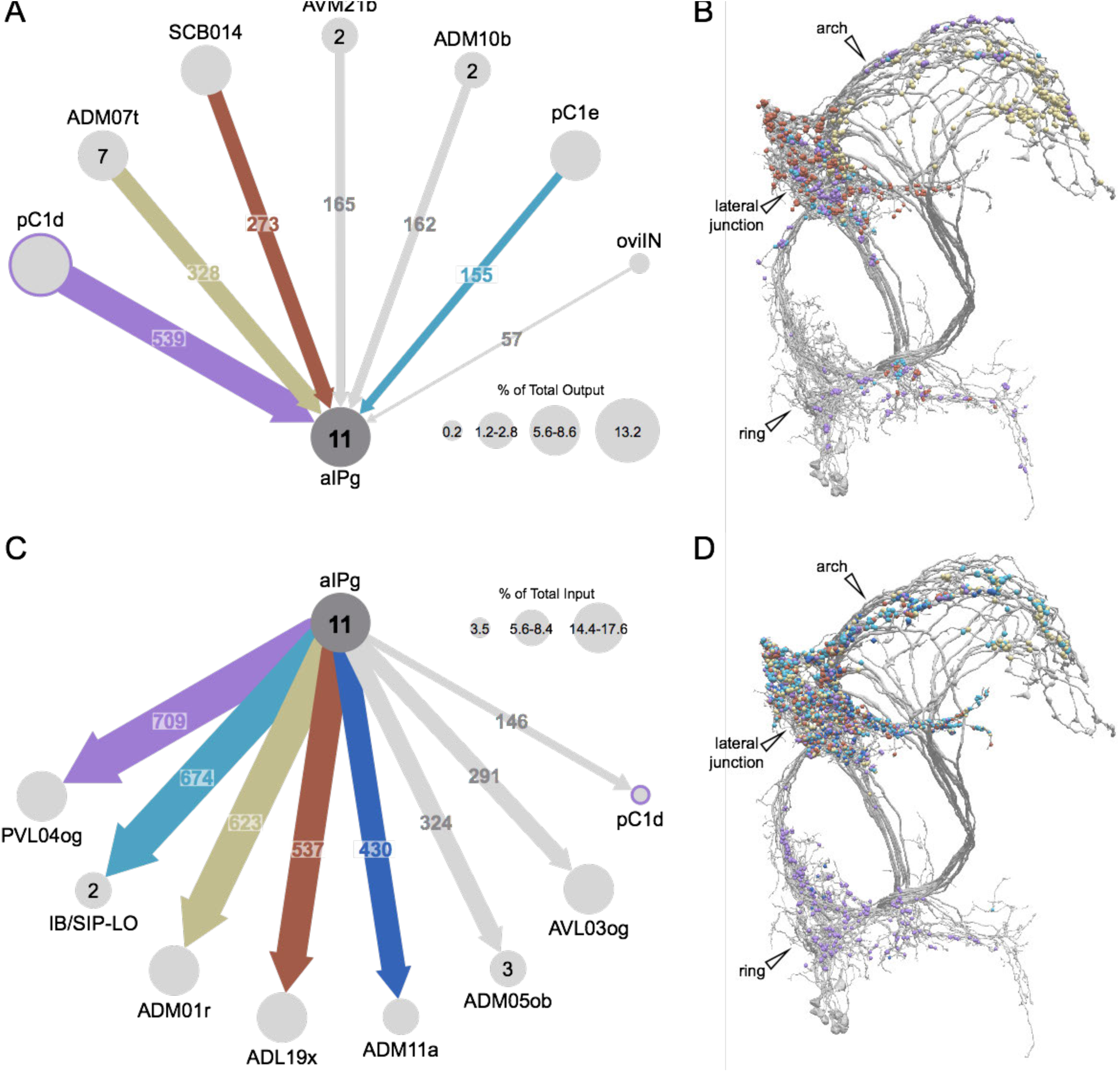
Major inputs to and outputs from aIPg neurons. (A) Inputs to the 11 aIPg neurons in the right hemisphere thresholded at 150 synapses (inputs to all 11 aIPg neurons pooled). OviIN was included due to its involvement in other female behaviors. The size of the circles representing input neurons indicates the percentage of their output (estimated by synapse number) that goes to aIPg neurons. (B) Positions on the aIPg arbors of post-synaptic sites where these connections occur, color coded to match diagram in (A). While SCB014 forms a large number of synapses to all aIPg neuron types in the lateral junction, the majority of the synapses from the ADM07t neurons are on type 1 and 3 neurons and occur within the arch of the lateral protocerebral complex (LPC). (C) Post-synaptic outputs of aIPg neurons, thresholded at 290 synapses; pC1d at 146 synapses is included to show reciprocal connections. The size of the circles representing the downstream targets of aIPg indicates the percentage of their input (estimated by synapse number) that comes from aIPg neurons. (D) Positions on the aIPg arbors of the presynaptic sites where these connections occur are shown, color coded to match diagram in (C). See Video 9 for details on the inputs to aIPg neurons and Video 10 for details on their outputs.

**Figure 10.**
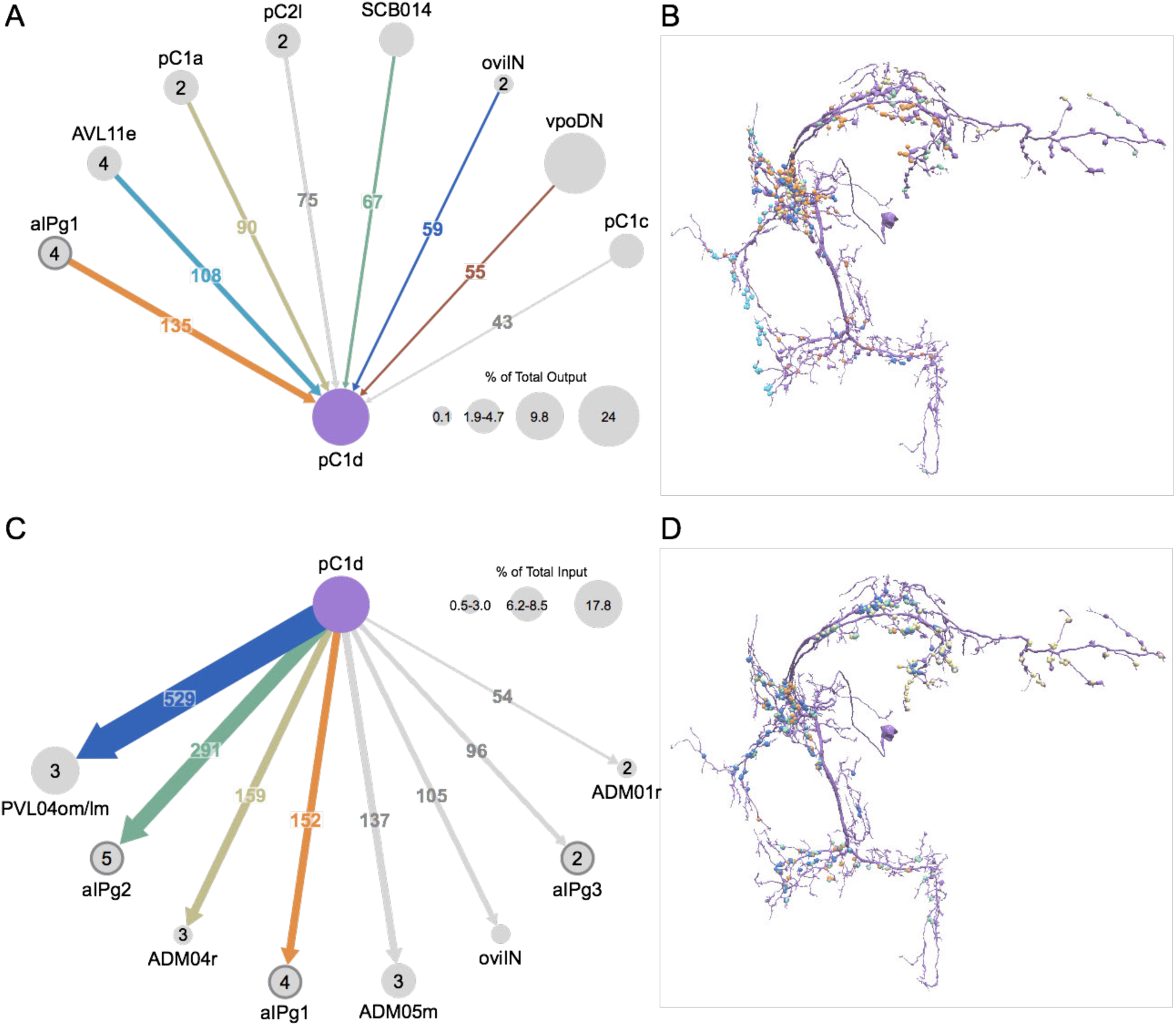
Major inputs to and outputs from pC1d. (A) Inputs to the single right-hemisphere pC1d neurons thresholded at 90 synapses with other neurons of interest included. The size of the circles representing input neurons indicates the percentage of their output (estimated by synapse number in the hemibrain volume) that goes to pC1d; note that vpoDN is a descending interneuron with most of its outputs likely to lie outside the volume. aIPg neurons are indicated with a dark grey outline. (B) Positions on the pC1d arbor of post-synaptic sites, color coded to match diagram in (A). (C) Post-synaptic outputs of pC1d thresholded at 90 synapses with other neurons of interest included. The size of the circles representing output neurons indicates the percentage of their input (estimated by synapse number) that comes from pC1d. aIPg neurons are indicated with a dark grey outline. (D) Positions on the pC1d arbors of the presynaptic sites where these connections occur are shown, color coded to match diagram in (C). See Video 11 for details on the inputs to pC1d and Video 12 for details on pC1d’s outputs.

Inputs to pC1d include several cell types implicated in oviposition and receptivity (Wang et al., 2020), such as pC1a and vpoDN, a neuron that controls vaginal plate opening (Wang et al 2020b; Figure 10A – B, Video 12). For example, pC1d receives 24.02% of the vpoDN’s output in the central brain as judged by synapse number (Figure 7A, Supplemental Table 3) (Wang et al. 2020b). Interestingly, synapses from aIPg type 1 neurons and pC1a are located at widely separated sites on the pC1d arbor (Figure 10B and Video 12), suggesting a potential spatial separation of the circuit connections underlying these two social behaviors. These connections may play a role in the flies’ ability to display one action at a time.

## Discussion

The circuits that govern aggression in *Drosophila* are known to be sexually dimorphic and are poorly understood in females. In this paper, we described female aggressive behaviors, uncover key components of the underlying neuronal circuits, develop genetic reagents to manipulate these neurons, and map their connections using EM-level connectomics. Specifically, we discovered the involvement of a subset of the aIPg lineage, a collection of cell types not previously implicated in social behaviors, in mediating female aggressive social interactions. Optogenetic activation with the channelrhodopsin CsChrimson dramatically increased aggression in lines labelling this subset of aIPg neurons, while inactivation diminished these actions. Analysis using EM-level connectomics revealed strong connectivity between these aIPg neurons and two members of the pC1 cluster, a group of related cell types previously linked with social behaviors (Hoopfer et al., 2015; Palavicino-Maggio et al., 2019; Scheffer et al., 2020; Wang et al., 2020; Zhou et al., 2014). In particular, pC1d is the top pre-synaptic input to aIPg neurons, devoting ∼13% of its output synapses to them. Behavioral tests using split-GAL4 lines cleanly labeling pC1d demonstrated its ability to increase female aggression. In contrast, we found no evidence for the involvement of the other four pC1 cell types, pC1a – c and pC1e, in aggression; however, we found that pC1e makes synapses onto aIPg neurons raising the possibility that this cell type plays a role that was not revealed by our behavioral assays.

### aIPg neurons mediate female aggressive behaviors

Innate behaviors, including aggression, have been proposed to result from the interplay of external stimuli and the internal state of the animal (Lorenz, 1963; Tinbergen, 1951). Neuronal populations that have the ability to bypass normally required sensory cues to induce aggressive behavior have been previously identified by experimentally activating select subsets of neurons in *Drosophila*. For example, male-specific neurons expressing *Drosophila* tachykinin (DTK) act as a hub mediating aggressive behaviors when activated and have been proposed to encode higher levels of motivation (Anderson, 2016; Asahina, 2017; Asahina et al., 2014; Hashikawa et al., 2018, 2017; Hoopfer, 2016). As aIPg appears to be both necessary and sufficient for performing a high level of female aggression, we consider this population to be performing a mediator function (Gregg, 2003; Siegel et al., 1999).

Neurons that provide input to aIPg could act as facilitators adjusting the degree of behavior or conveying specific sensory information important for its initiation and execution. For example, pC1d activation was able to drive aggressive behaviors but was not essential for these social interactions, indicating it may act as a facilitator. Stimulation of pC1e neurons alone did not result in aggressive phenotypes under the conditions tested, or increase the aggression observed when activated with pC1d. However, multiple external factors required for aggression in wild-type females were not assessed in our activation experiments. For example, diet and the presence of yeast in the arena are known to be important factors determining the level of female aggression (Ueda and Kidokoro, 2002). Experiments examining the influence of such conditions on the circuit and behavioral dynamics will be needed to evaluate pC1e’s potential role. As circuits can perform more than one behavioral output, pC1e and the rest of this circuit could also be involved in other interactions that will need to be studied using additional assays.

### aIPg type 1, type2 and type3 neurons differ in their pre- and post-synaptic connections

The aIPg neurons we studied are three distinct cell types that differ in morphology and connectivity. Our efforts to derive split-GAL4 lines specific for each of these cell types have been unsuccessful, which has limited our ability to explore their individual roles. Nor have we yet performed physiological experiments that might reveal distinct features of the responses of these neurons when the circuit is activated. Nevertheless, we propose that the three aIPg types found in our split-GAL4 lines make distinct contributions to the phenotypes we observed, as there are many differences in their inputs and outputs (Figure 11). For example, while pC1d makes strong connections onto all three aIPg types, it only receives strong recurrent feedback from type 1 neurons. Similarly, SCB014, which is the third top input to the aIPg neurons, predominately sends its projections to type 2 neurons. There are likewise many differences in downstream targets, some of which connect to descending neurons (Figure 11). Generating split-GAL4 lines that specifically target such cell types will allow us to perform experiments to explore their roles in aggressive behaviors.

**Figure 11.**
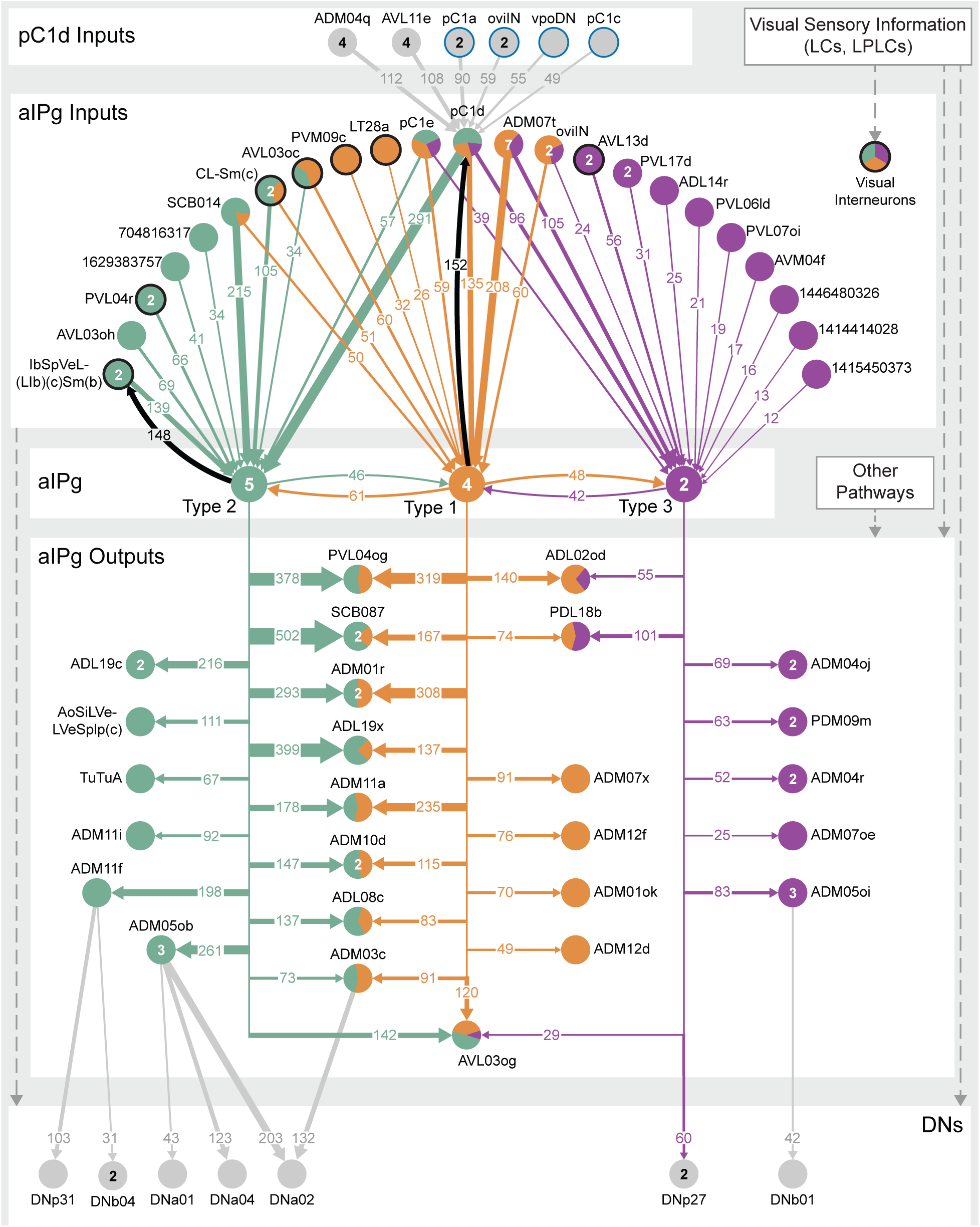
Proposed circuit underlying female aggressive behaviors. A circuit diagram showing the key neuronal pathways we uncovered in our behavioral and connectomics analyses. The numbers in arrows represent synapse number. We propose that pC1d facilitates female aggressive behaviors through aIPg type 1 (orange), type 2 (green), and type 3 neurons (purple), which act as a mediator of these social interactions. The aIPg type 1 neurons, based on their connections with the other two types and feedback onto pC1d, appear to be important for the recurrency within the circuit. We propose that visual information enters the circuit through interneurons (black outline) that innervate aIPg neurons (see Figure 11-figure supplement 1) or at later nodes in the circuit. Like pC1d, other neuronal populations (color-coded by the proportion of their synapses on each aIPg type) also provide different proportions of their output to the aIPg type 1, aIPg type 2 and aIPg type 3 subtypes. The EM dataset further identified differences in the post-synaptic targets of aIPg type 1, aIPg type 2 and aIPg type 3 neurons, which are shown color-coded by the relative proportion of their synaptic input they receive from each aIPg type. Additionally, while only aIPg type 3 neurons directly connect with a descending interneuron (DN), we identified four post-synaptic targets of the aIPg type 1, aIPg type 2 and aIPg type 3 neurons that form synapses onto DNs, which may function as motor outputs of the circuit. The top four inputs to pC1d and three other neurons of interest are shown. These include neurons implicated in other female behaviors (blue outline), including oviposition (oviIN, pC1a) and mating (vpoDN), suggesting communication between the circuits underlying these interactions. Inputs to the aIPg neurons were thresholded at 6 synapses per aIPg neuron, resulting in collective thresholds of 24, 30, and 12 synapses to aIPg type 1, aIPg type 2, and aIPg type 3, respectively. Outputs of the aIPg neurons were similarly adjusted based on the aIPg type, with 48, 60, and 24 synapses used as the collective thresholds for aIPg type 1, aIPg type 2, and aIPg type 3, respectively. Connections with DNs were thresholded at 30 synapses. Connections between IbSpVeL-(LIb)(c)Sm(b), a strong reciprocal target of aIPg type 2, and DNa03 (334 synapses) and DNb01 (32 synapses), as well as connections between pC1d and DNa01 (40 synapses) are not shown.

### Activation of aIPg neurons generated a persistent internal state resulting in continued aggressive social interactions

In both males and female flies, aggression-related behaviors have been found to extend beyond the stimulation period of certain neuronal cell types (Deutsch et al., 2020; Hoopfer et al., 2015), suggesting a persistent internal state. We did observe head butting and fencing behaviors after the cessation of the optogenetic stimulus when lines labeling aIPg cells were used. Additionally, the extent of these behaviors depended on stimulus conditions. In contrast, levels of post-stimulus aggressive behavior similar to those observed in aIPg lines were not found when activating pC1d or pC1e, or a combination of these two cell types. Persistence of aggression after stimulating a combination of pC1d and pC1e has been reported following a 5 minute stimulation paradigm (Deutsch et al., 2020) rather than the 30 second stimulation we used. It is possible that such persistence is the result of downstream activation of aIPg neurons. However, it is important to keep in mind that the stimulus conditions required to induce persistent aggression are more intense than those needed to generate high levels of aggression during the stimulus period itself and the ethological relevance of persistence in female aggression is unclear. Female aggressive bouts range from 10 to 30 seconds depending on their mated status as opposed to an average of 80 seconds for single-housed males (Bath et al., 2017; Yuan et al., 2014). Additionally, female flies do not appear to establish dominance, and aggressive behaviors are not found to escalate over the course of a trial (Nilsen et al., 2004; Vrontou et al., 2006). While some persistence is needed to maintain aggressive behaviors during bouts lasting tens of seconds, it is unclear what role longer persistence would play. Consistent with this view, we observed that aggressive behaviors took only a few 10s of seconds to return to baseline levels after the stimulation of aIPg neurons.

Multiple recurrent circuit motifs that could provide the basis for persistent neural activity over this time scale were revealed by our connectomics analyses. In particular, the feedback between aIPg type 1 neurons and pC1d, and recurrent connections between the aIPg neurons themselves, are obvious candidates as all of these neurons are cholinergic and thus putatively excitatory. The higher levels of stimulation required to induce persistence raises the possibility that this phenotype is an experimental artefact of non-physiological levels of stimulation of those same circuits. Nevertheless, it seems likely that more subtle changes in brain state persist, and the circuit mechanisms of such changes remain an open question. For example, our RNA profiling experiments raise the possibility of a role for the neuropeptide sNPF.

Changes in brain state, such as those occurring during aggressive encounters, also influence sensory processing. For example, olfactory cues paired with a male fly winning an aggressive encounter become associated with reward (Kim et al., 2018). We observed that females quickly altered their speed and direction following aIPg stimulation, appearing to approach other flies. These behaviors raise the possibility that visual signals influence the “decision” to be aggressive. For example, certain features of the target, including its size and distance, might be important to drive aggression through aIPg neurons. Another non-exclusive hypothesis is that aggression includes behaviors that rely on vision, such that aIPg activation recruits orientation circuits and other pathways also used by different types of social interactions. Connections were identified in the hemibrain between visual interneurons and all three types of the aIPg neurons (Figure 11 and Figure 11-figure supplement 1), which would be in a position to convey information about the target and salient aspects of the environment. Specific types of lobular columnar cells convey different information (Keleş and Frye, 2017; Wu et al., 2016), and these inputs differ amongst the aIPg types. For example, optogenetic activation of LC10 can elicit behaviors resembling reaching (Wu et al, 2016) and components of courtship behavior (Ribeiro et al., 2018), and these neurons connect through an interneuron to aIPg type 1 and type 2. Additionally, visual information could be integrated at later nodes within the circuit, as has been shown for P9 descending neurons involved in mediating courtship behavior (Bidaye et al., 2019).

### Interactions between pC1d and neurons implicated in other female social behaviors

Our analysis of the pC1d and aIPg also revealed connections to neurons known to be involved in other female-specific behaviors, including oviposition. Links between oviposition and female aggression have been observed in field work on other *Drosophila* species, with increased aggressive behaviors occurring on egg laying sites (Shelly, 1999). Supporting this close relationship between the behaviors, pC1d receives innervation from pC1a, a cell type involved in the egg laying pathway (Wang et al., 2020). Interestingly, Wang et al. (2020) noted differences in the depolarization of pC1d and pC1e following activation of the sex-peptide abdominal ganglion (SAG) neuron in the sex-peptide pathway. While these are not as extensive as seen in pC1a, they provide a potential neural basis for the reported effect of mating status on female aggressive behavior (Bath et al., 2020, 2018; Ueda and Kidokoro, 2002). Further supporting the close tie between these two social behaviors, we found that a very high proportion of the synaptic output of vpoDN in the central brain goes to pC1d and oviIN innervates both pC1d and aIPg neurons (Wang et al., 2020a,b). Mating behavior and aggression also overlaps at the P1/pC1 level in male flies, raising the possibility of a common node in both sexes (Hoopfer et al., 2015; Wang et al., 2020). Due to their connectivity, we expect aIPg, pC1d, and pC1e to be involved in a range of social interactions in addition to aggression.

**Table 1.** Split-GAL4 lines

| Name | Number | AD Parent | DBD Parent |
| --- | --- | --- | --- |
| emptySS | emptySS | BPp65ADZp in attP40 | BPZpGDBD in attP2 |
| alPgSS1 | SS36564 | VT064565-pBPp65ADZpUw in attP40 | VT043699-pBPZpGdbdUw in attP2 |
| alPgSS2 | SS36551 | VT043699-pBPp65ADZpUw in attP40 | R11A07-pBPZpGdbdUw in attP2 |
| alPgSS3 | SS32237 | R11A07-pBPp65ADZpUw in attP40 | VT043700-pBPZpGdbdUw in attP2 |
| alPgSS4 | SS47478 | R72C11-pBPp65ADZpUw in VK00027 | R11A07-pBPZpGdbdUw in attP2 |
| alPgSS5 | SS56964 | R11A07-pBPp65ADZpUw in attP40 | R72C11-pBPZpGdbdUw in attP2 |
| pC1dSS1 | SS56987 | R35C10-pBPp65ADZpUw in JK73A | R71A09-pBPZpGdbdUw in attP2 |
| pC1dSS2 | SS57598 | R71A09-pBPp65ADZpUw in JK73A | R26F09-pBPZpGdbdUw in attP2 |
| pC1dSS3 | SS43274 | VT25602-pBPp65ADZpUw in attP40 | VT002064-pBPZpGdbdUw in attP2 |
| pC1dSS4 | SS59331 | R35C10-pBPp65ADZpUw in JK73A | VT002063-pBPZpGdbdUw in attP2 |
| pC1eSS1 | SS59336 | R35C10-pBPp65ADZpUw in JK73A | R60G08-pBPZpGdbdUw in attP2 |
| pC1eSS2 | SS39313 | R60G08-pBPp65ADZpUw in attP40 | VT002063-pBPZpGdbdUw in attP2 |
| pC1eSS3 | SS59433 | R60G08-pBPp65ADZpUw in attP40 | R35C10-pBPZpGdbdUw in attP2 |
| pC1a-c | SS75230 | R18A05-pBPp65ADZpUw in attP40 | dsx-ZpGal4DBD/TM6B |

### Concluding remarks

The female fly must integrate information from sensory cues about the external world—such as the presence of other individuals, food and egg laying sites—with internal states—such as hunger and mating status—when weighing when to initiate or abandon aggressive interactions. Our work provides a foundation for future studies aimed at understanding the complex social behavior of aggression by identifying key neuronal cell types, placing them in the context of a larger neuronal circuit, and providing tools for their manipulation.

## Acknowledgements

We thank Drs. U. Heberlein, D. J. Anderson, V. Chiu, K. Longden, K. Wang, F. Wang, A. Nern, and T. Wolff as well as the Janelia community for their helpful suggestions during the course of this work and their comments on the manuscript. We thank the Janelia Project Technical Resources team (led by Gudrun Ihrke) for assistance with the manual correction of automated fly trajectories (Rebecca Arruda and NC) and performing neuronal segmentation of confocal images (Claire Managan). We also thank the Quantitative Genomics facility at Janelia for performing RNA profiling of the aIPg, pC1d, and pC1e lines, M. Eddison and J. Simon for their help in setting up the inactivation experiments and Igor Siwanowicz and Wyatt Korff for their assistance with the high-speed videos. We thank the Fly Light team for generating the images of GAL4 expression patterns and the connectome annotation team for their help with proofreading.

## Materials and Methods

### Fly stocks

All experiments used virgin female flies unless otherwise stated. Flies were reared on standard cornmeal molasses food at 25°C and 50% humidity. For optogenetic activation experiments, flies were reared in the dark on standard food supplemented with retinal (Sigma-Aldrich, St. Louis, MO), 0.2 mM all *trans*-retinal prior to eclosion and 0.4 mM all *trans*-retinal post eclosion. Hemi-driver lines were created using gateway cloning as previously described (Dionne et al. 2018).

In addition to the split-GAL4 lines (see Table 1), the following fly strains were used: (1) 20XUAS-CsChrimson-mVenus in attP18 (Klapoetke et al., 2014); (2) w; Enhancer1-ADp65 (attP40); Enhancer2-ZpGAL4DBD (attP2) referred to as EmptySS and is described previously (Pfeiffer et al., 2010; Luan et al., 2006; Aso et al., 2014); (3) MCFO-1 (Nern et al., 2015) (4) UAS-GFP: pJFRC2-10XUAS-IVS-mCD8::GFP in attP2 (Pfeiffer et al., 2010); (5) UAS-impTNT-HA; (6) (w-); UAS-TNT-E (Sweeney et al. 1995).

### Optogenetic activation behavioral testing

Groups of 13 – 18 group-housed virgin female flies (5 – 10 days post-eclosion) were tested at 25°C and 50% relative humidity in a 127 mm circular arena with a center depth of 3.5 mm as described previously (Robie et al. 2017, Wu et al. 2014). Flies were loaded into the arena using an aspirator. For activation of neurons expressing CsChrimson, the arena was uniformly illuminated with 617 nm LEDs (Red-Orange LUXEON Rebel LED - 122 lm; Luxeon Star LEDs, Brantford, Canada) at the power density specified in the figure legend. Unless otherwise stated, all trials were performed under white-light illumination. For each trial, flies were acclimatized to the area for 30 s prior to the delivery of a single constant stimulus lasting 30 s. Pulse stimulation at 0.1 mW/mm^2^ was given in 30 s intervals with an inter-stimulus interval of 30 to 60 s. The pulse width was kept constant at 10 ms while the pulse number and period varied. Videos were recorded from above using a camera (ROHS 1.3 MP B and W Flea3 USB 3.0 Camera; Point Grey, Richmond, Canada) with an 800 nm long pass filter (B and W filter; Schneider Optics, Hauppauge, NY) at 30 frames per second and 1024 x 1024 pixel resolution. Flies were tracked using Ctrax (Branson et al. 2009) followed by automated classification of behavior with JAABA classifiers (see Robie et al. (2017) or Supplementary file 1 for performance summary). For experiments examining individual behaviors, the identities of the fly tracks were manually corrected using the FixErrors GUI available with Ctrax. To validate classifiers, we manually scored a separate group of ground-truth videos for head butting and fencing using the ‘ground-truthing’ mode in JAABA. Using the balanced random feature, set of randomly selected frames from the ground-truth videos was manually labeled as positive or negative for the behavior of interest on a frame-by-frame basis.

Activation experiments detailed in Figure 3 and Figure 5 supplement 9 were performed in 16 mm diameter x 12 mm high aggression chambers using same LEDs and camera setup as detailed above. For these experiments, two flies (genotypes and sex specified in figure) were introduced into the arena with an aspirator. Flies were acclimatized to the arena for 30 s prior to delivery of a single constant stimulus lasting 30 s. Flies were tracked using the Caltech FlyTracker (http://www.vision.caltech.edu/Tools/FlyTracker/) followed by automated classification of behavior with a JAABA classifier for head butting and fencing behaviors (see Supplementary file 1 for performance summary).

For high speed videos, two flies were loaded into an arena using cold anesthesia and an aspirator. The arena was illuminated with a ring light and a LED gooseneck. Constant stimulation was provided from above during the recording at 0.45 mW/mm^2^ with a 625 nm light source. Videos were recorded with a Photon Fastcam Mini camera at 1000 frames per second.

Receptivity assays were performed as detailed in Wang et al. (2020). Briefly, one virgin female and one wild-type single housed virgin male were transferred into a 10 mm diameter x 2 mm height arena through aspiration. Flies were videotaped under white-light illumination for 30 min and photostimulated at 0.4 mW/mm^2^ as described by Wang et al. (2020). The copulation latency was examined manually.

### Inactivation behavioral testing

Virgin female flies were group-housed at a density of 20 – 40 females per vial at 22°C on dextrose media (79 g agar, 275 g yeast, 520 g cornmeal, 1100 g Dextrose, 87.5 mL 20% Tegosept, 20 mL Proprionic Acid in 11000 mL of water) for 21 – 28 days. Female flies were single housed for 5 – 7 days and subsequently starved on 1% agarose in water for 20 – 24 hours prior to testing. Rearing and housing were performed in a light cycling incubator set for 12/12 hr light/dark cycle. All inactivation experiments were run at 25°C and 40% humidity and performed with two female flies per arena.

Assays were performed in previously described acrylic multi-chamber aggression arenas (Hoopfer et al. 2015, Kim et al. 2018, Asahina et al. 2014). Each circular arena measured 16 mm diameter x 12 mm high and was coated with Insect-a-Slip (Bioquip Products, Rancho Dominguez, CA) on the walls and SurfaSil Siliconizing Fluid (Thermo Fisher Scientific, Waltham, MA) on the clear acrylic top plate to confine the flies to the bottom plate. The floor of the arenas was composed of a uniform layer (∼1 mm thick) of apple juice-agarose food (2.5% (w/v) sucrose and 2.25% (w/v) agarose in apple juice) that contained a ∼1 mm spot of live yeast (Fleischmann, Cincinnati, Ohio) placed in approximately the center of each arena. Flies were illuminated from beneath with visible light and recorded with ambient overhead room lighting. Flies were introduced into the chamber with gentle aspiration through a hole in the top plate and allowed to acclimatize for 30 s – 1 min prior to recording. For automated analysis, flies were tracked using the Caltech FlyTracker (http://www.vision.caltech.edu/Tools/FlyTracker/) followed by automated classification of behavior with a JAABA classifier for head butting and fencing behaviors (see Supplementary file 1 for performance summary). For manual analysis, videos were blinded and scored for head butting and fencing behaviors using JWatcher software (http://www.jwatcher.ucla.edu/). Data were reported for each pair tested for manually annotated analysis.

### Immunohistochemistry and imaging

Dissection and immunohistochemistry of fly brains were carried out as previously described (Jenett et al., 2012, Aso et al. 2019). Each split-GAL4 line was crossed to the same Chrimson effector used for behavioral analysis. Full step-by-step protocols can be found at https://www.janelia.org/project-team/flylight/protocols.

For single cell labelling of aIPg neurons, we used the MultiColor FlpOut (MCFO) technique (Nern et al., 2015). For MultiColor FlpOut (MCFO) experiments, the MCFO stock was crossed to a split-GAL4 line. Flies were collected after eclosion, transferred to a new food vial and incubated in a 37°C water bath for 20–25 min. These flies were dissected and underwent whole-mount immunohistochemistry and confocal imaging (Dolan et al. 2018).

All imaging was performed on LSM 700, 710 and 780 confocal microscopes (Zeiss) using ZEN software with a custom MultiTime macro. For all images except those in Figure 1 – figure supplement 1, Figure 5 – figure supplement 1, and Figure 6 – figure supplement 1, the macro was programmed to automatically select appropriate laser power and/or gain for each sample, resulting in independent image parameters between samples. The images in Figure 1 – figure supplement 1, Figure 5 – figure supplement 1, and Figure 6 – figure supplement 1 were captured on the same LSM 710 microscope with all imaging parameters held fixed to maximize comparability.

### Connectomics Analysis

Data collection and processing is specified in Zheng et al. (2018) for the FAFB dataset and Scheffer et al. (2020) for the hemibrain dataset. Hemibrain data was queried using NeuPrint and v1.0 of the connectome (neuprint.janelia.org). Visualizations of neuronal morphologies from the hemibrain dataset were generated in NeuTu. Thresholds were used in order to limit the number of neurons in the figures to those connections with the most synapses. The specific thresholds used are given in each figure. In Figures 9 – 11, neurons of interest that fell below the thresholds indicated were also included. A complete list of synaptic connections can be found in NeuPrint.

For anatomical videos made in Blender (https://www.blender.org/), neuron and ROI meshes were pulled directly from DVID using a set of python scripts (https://github.com/connectome-neuprint/neuVid). Adobe Premier Pro and Adobe Illustrator were used to add text and diagrams to the videos, respectively. Recordings for the narrations were performed using Camtasia.

### Statistics

No statistical methods were used to pre-determine sample size. Sample size was based on previous literature in the field and experimenters were not blinded in most conditions as almost all data acquisition and analysis were automated. However, inactivation experiments in which manual quantification was performed were blinded. All are biological replicates and behavioral data are representative of at least 2 independent trials per experiment.

For each experiment, the experimental and control flies were collected, treated and tested at the same time. A Mann–Whitney *U* test or Kruskal–Wallis test and Dunn’s post hoc test was used for statistical analysis. Comparisons with more than one variant were first analyzed using two-way ANOVA. All statistical analysis was performed using Prism Software (GraphPad, version 7). P values are indicated as follows: ****p < 0.0001; ***p < 0.001; **p < 0.01; and *p < 0.05. See Supplementary file 5 for exact *p*-values for each figure.

For boxplots, lower and upper whiskers represent 1.5 × IQR of the lower and upper quartiles, respectively; boxes indicate lower quartile, median, and upper quartile, from bottom to top. When all points are shown, whiskers represent range and boxes indicate lower quartile, median, and upper quartile, from bottom to top. Shaded error bars on graphs are presented as mean ± s.e.m.

## Supplementary Figures

**Figure 1 – figure supplement 1.**
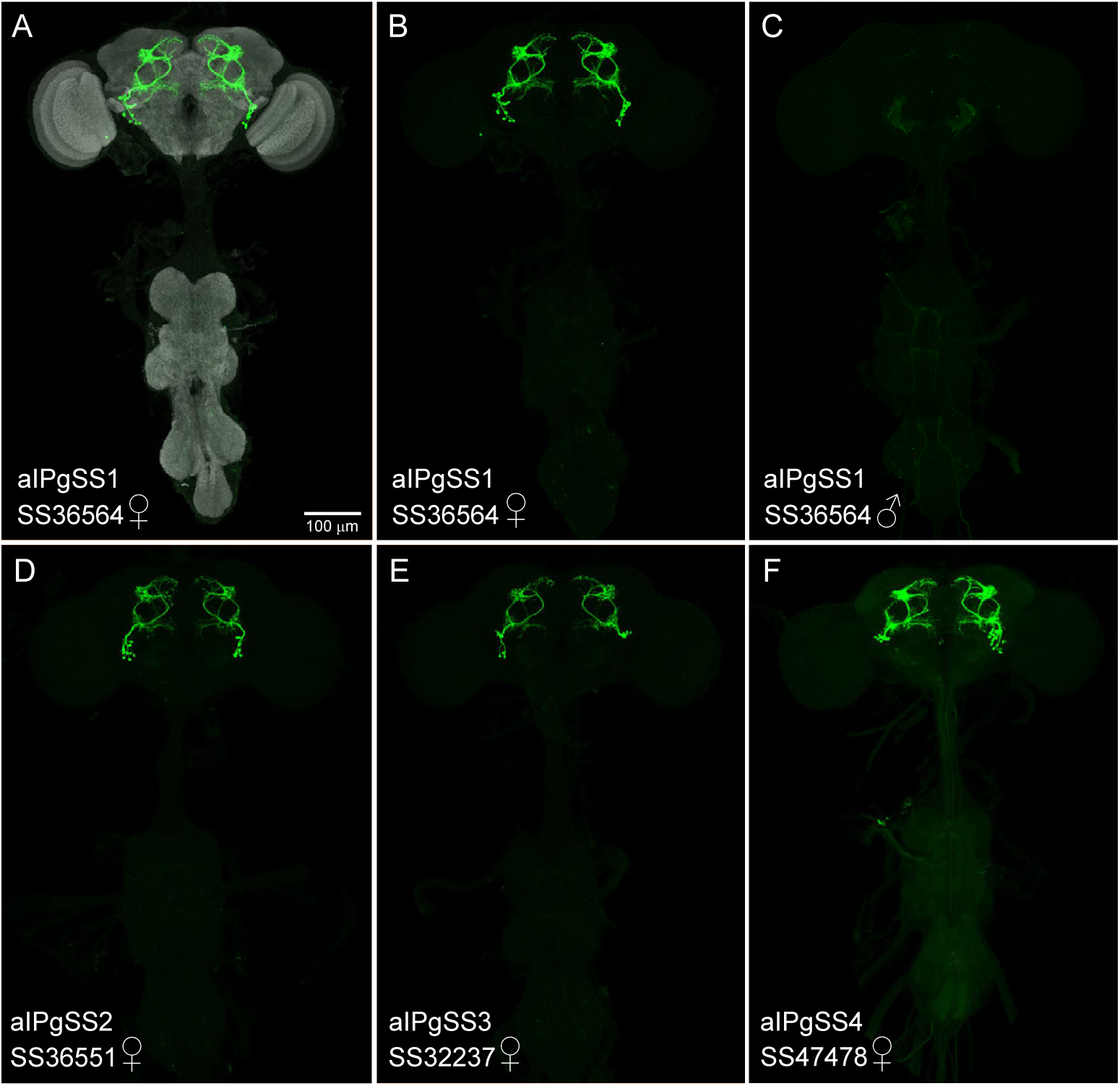
Expression patterns of aIPg split-GAL4 lines. (A – F) Maximum intensity projections (20x) of the brains and ventral nerve cords of the indicated split-GAL4 lines crossed to 20xUAS-CsChrimson::mVenus and stained with anti-GFP antibody are shown. As the images are visualizations of the optogenetic effector itself, these images indicate the relative expression levels of the effector in the different split-GAL4 lines. Sex of the imaged brain is indicated as is the stock name of split-GAL4 line. The scale bar shown in A applies to panels A – F. (D’ – F’) Enlargements of the central brain of the images shown in D – F. (A – C) The SS36564 (aIPgSS1) expression pattern is shown; the neuropil reference channel is shown in A (gray). SS36564 expresses in 11.3±1.5 cells per female brain hemisphere (n = 8). Note the lack of expression in aIPg neurons in the male central nervous system (C) imaged under identical conditions to that of the female (B). (D, D’) Expression pattern of SS36551 (aIPgSS2) in females. SS365551 expresses in 8.9±1.4 cells per female brain hemisphere (n = 8). A few cells with distinct, but related morphology are observed in males. (E, E’) Expression pattern of SS32237 (aIPgSS3) in females. SS32237 expresses in 7.8±0.89 cells per female brain hemisphere (n = 8). A few cells with distinct, but related morphology are observed in males. (F, F’). Expression pattern of SS47478 (aIPgSS4) in females; this line expresses in 12.8±1.8 cells per female brain hemisphere (n = 8). No expression was seen in males.

**Figure 1 – figure supplement 2.**
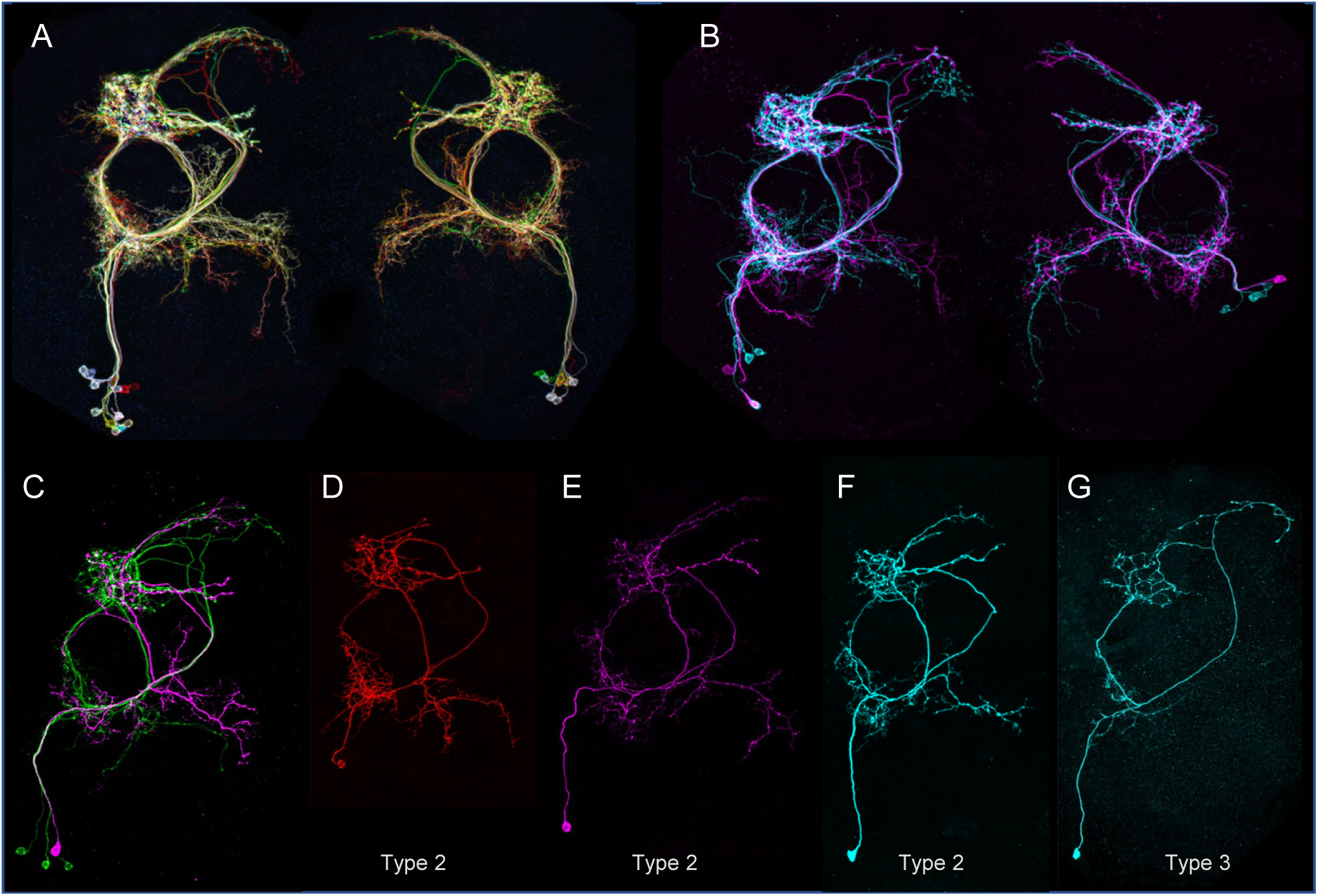
Morphologies of individual aIPg type 1 – 3 neurons. The images in all panels were generated by stochastic labeling of aIPg split-GAL4 lines using the MultiColor FlipOut method (Nern et al. 2015). (A, B) Images from SS36564 (aIPgSS1). Both brain hemispheres, and multiple aIPg cells, are shown. (C, E, F, G) Images from SS36551 (aIPgSS2). One brain hemisphere is shown. (D) Image from SS47478 (aIPgSS4). One brain hemisphere is shown. (D, E, F) show examples of type 2 aIPg cells, while (G) shows a type 3 aIPg cell.

**Figure 1 – figure supplement 3.**
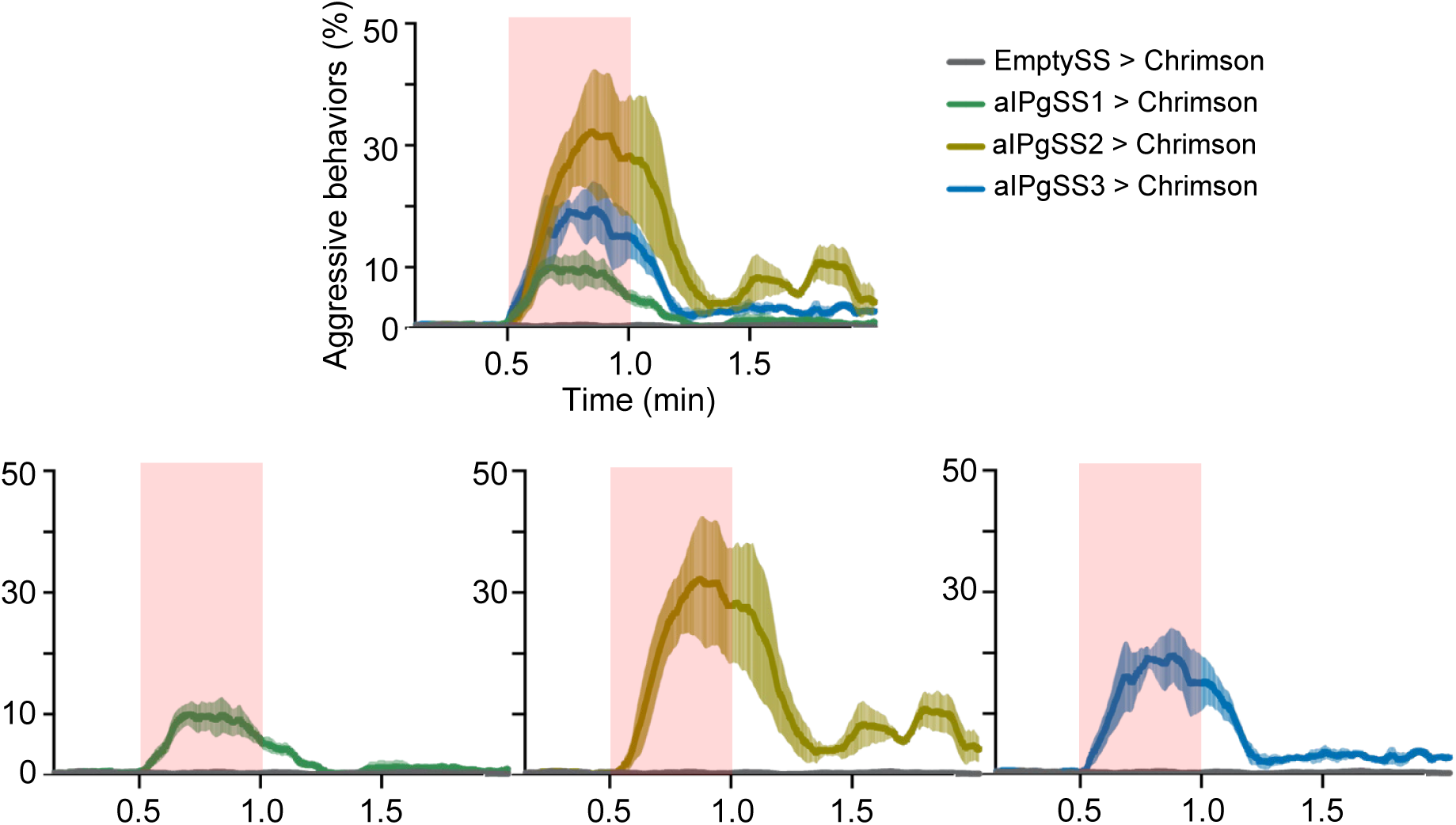
Additional aIPg split-GAL4 lines also induce aggressive behavior. Percentage of flies engaging in aggressive behaviors over the course of a 2-minute trial during which a 30-second 0.1 mW/mm^2^ continuous light stimulus was delivered is shown for three different split-GAL4 lines that drive expression in the aIPg cells. The expression pattern of these three aIPgSS lines are shown in Figure 1 – figure supplement 1. The red shading indicates the stimulus period. The mean is represented as a solid line and shaded error bars represent variation between experiments. For clarity, the individual plots for each line and the EmptySS > Chrimson are also shown. Each experiment included approximately 15 flies. EmptySS > Chrimson, n = 7 experiments; aIPgSS1 > Chrimson, n = 5 experiments; aIPgSS2 > Chrimson, n = 3 experiments; aIPgSS3 > Chrimson, n = 4 experiments.

**Figure 1 – figure supplement 4.**
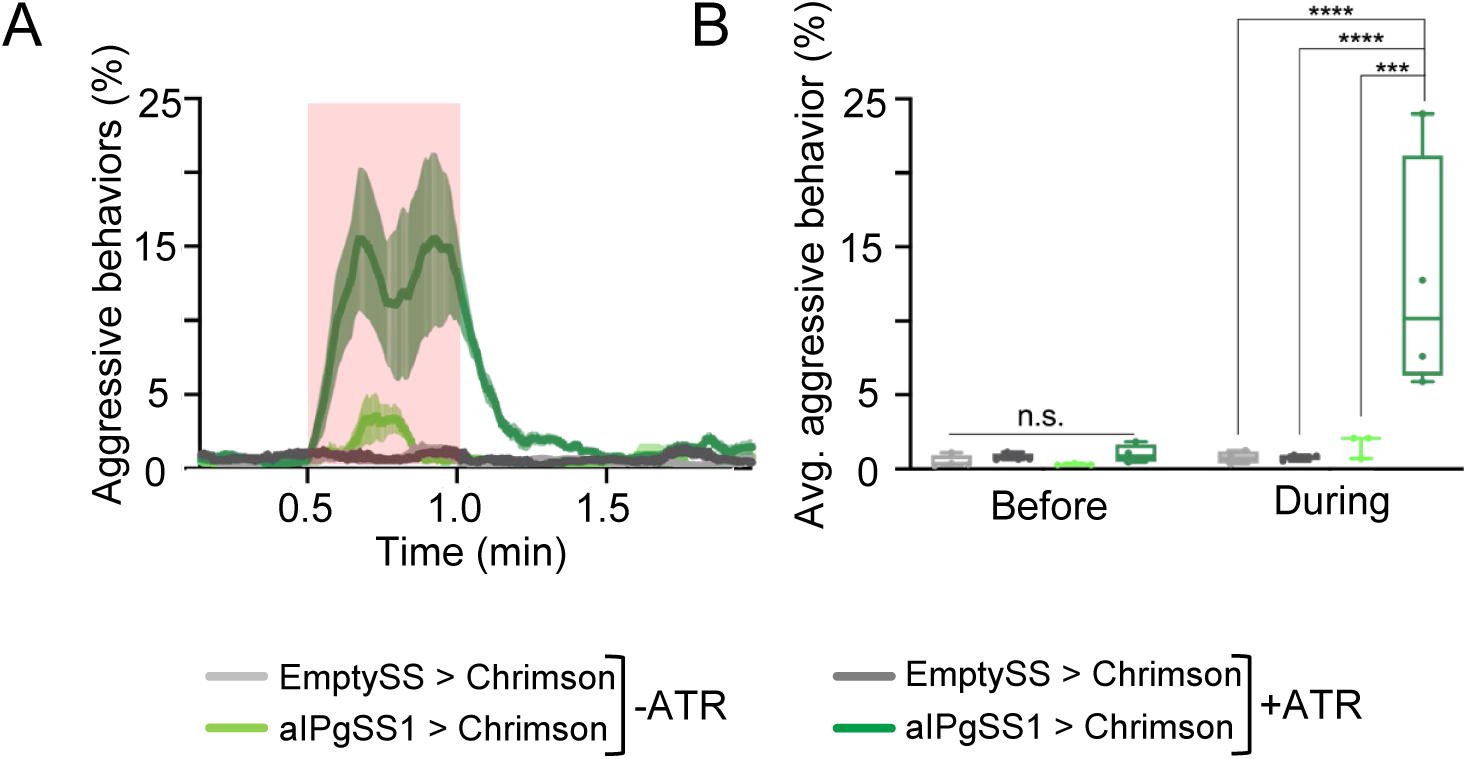
Optogenetic activation of aggression with Chrimson requires on feeding all *trans*-retinal. (A) Percentage of flies that engaged in aggressive behaviors. Flies were raised with food supplemented with all *trans*-retinal (+ATR) or not (-ATR). A 30-second 0.1 mW/mm^2^ continuous light stimulus was delivered 30 sec after the start of a 2-minute trial as indicated by the red shading. The mean is represented as a solid line and shaded error bars represent variation between experiments. (B) Average percentage of flies engaging in aggressive behaviors over the 30 second period prior to or during the stimulus delivery in A. Points represent separate experiments consisting of approximately 15 flies. +ATR: EmptySS > Chrimson, n = 4 experiments; aIPgSS1 > Chrimson, n = 4 experiments;-ATR: EmptySS > Chrimson, n = 4 experiments; aIPgSS1 > Chrimson, n = 3 experiments. Box-and-whisker plots show median and interquartile range (IQR); whiskers show range. A two-way ANOVA with a Tukey’s multiple comparisons post-hoc test was used for statistical analysis. Asterisk indicates significance from 0: ***p<0.001; n.s, not significant.

**Figure 1 – figure supplement 5.**
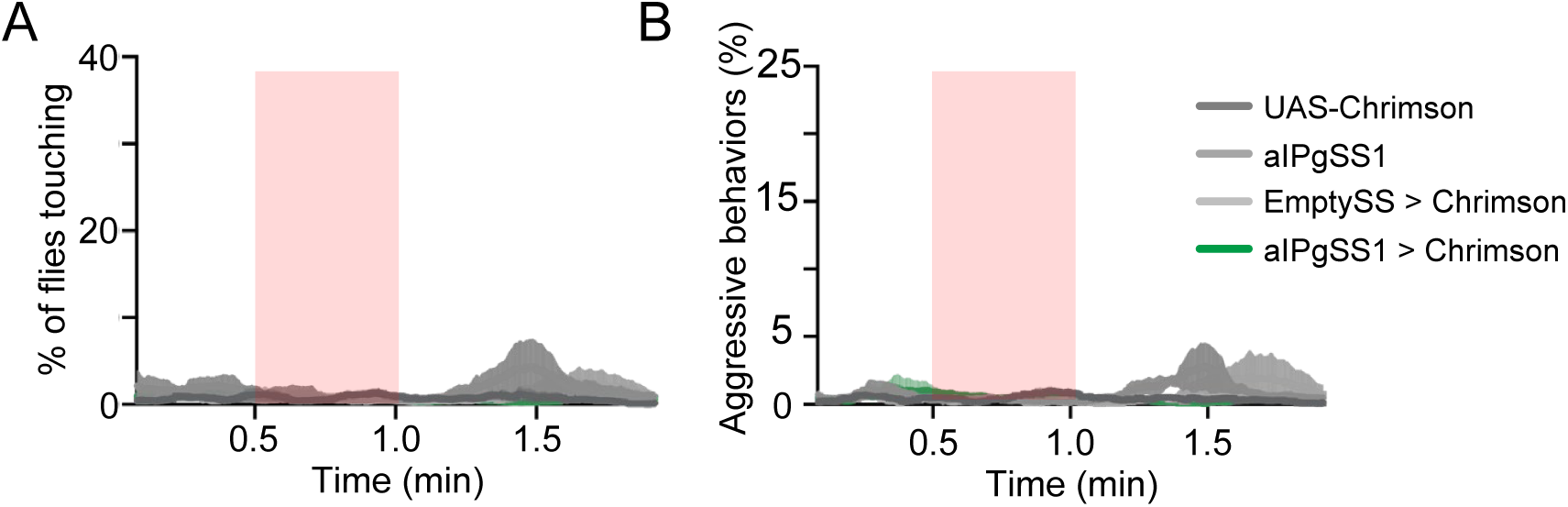
Optogenetic stimulation of aIPgSS1>Chrimson males does not result in aggressive behavior. (A – B) Percentage of male flies touching (A) or performing aggressive behaviors (fencing and head butting) (B) over the course of a 2-minute trial during which a 30-second 0.4 mW/mm^2^ continuous light stimulus was delivered as indicated by the red bar. The mean is represented as a solid line and shaded error bars represent variation between experiments. Each experiment included approximately 15 flies. 20xUAS-CsChrimson, n = 5 experiments; aIPgSS1, n = 4 experiments; EmptySS > 20xUAS-CsChrimson, n = 3 experiments; aIPgSS1 > 20xUAS-CsChrimson, n = 6 experiments.

**Figure 1 – figure supplement 6.**
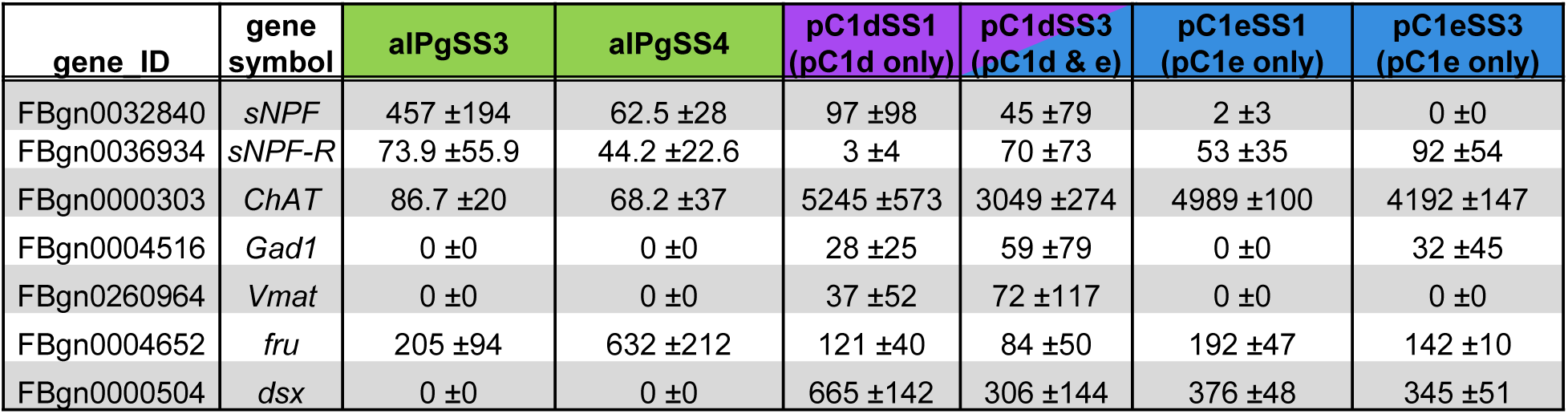
aIPg neurons are cholinergic, fru^+^ and sNPF^+^; pC1 neurons are cholinergic and dsx^+^. Expression (TPM) of genes related to neurotransmitters and sex determination pathways, *sNPF, sNPF-R*, *dsx* and *fru.* Cells were dissociated from dissected brains and sorted on the basis of their expression of the indicated split-GAL4 driver: aIPgSS3 (n = 2), aIPgSS4 (n = 3), pC1dSS1 (n = 3), pC1dSS3 (n = 6), pC1eSS1 (n = 3), and pC1eSS3 (n = 3). Cell sorting and RNA profiling was done as described in Aso et al. (2019). The mean and standard deviation of the indicated number of biological repeats are shown. Complete transcript data were deposited to NCBI Gene Expression Omnibus [author note: submission will be done while paper is under review].

**Figure 1 – figure supplement 7.**
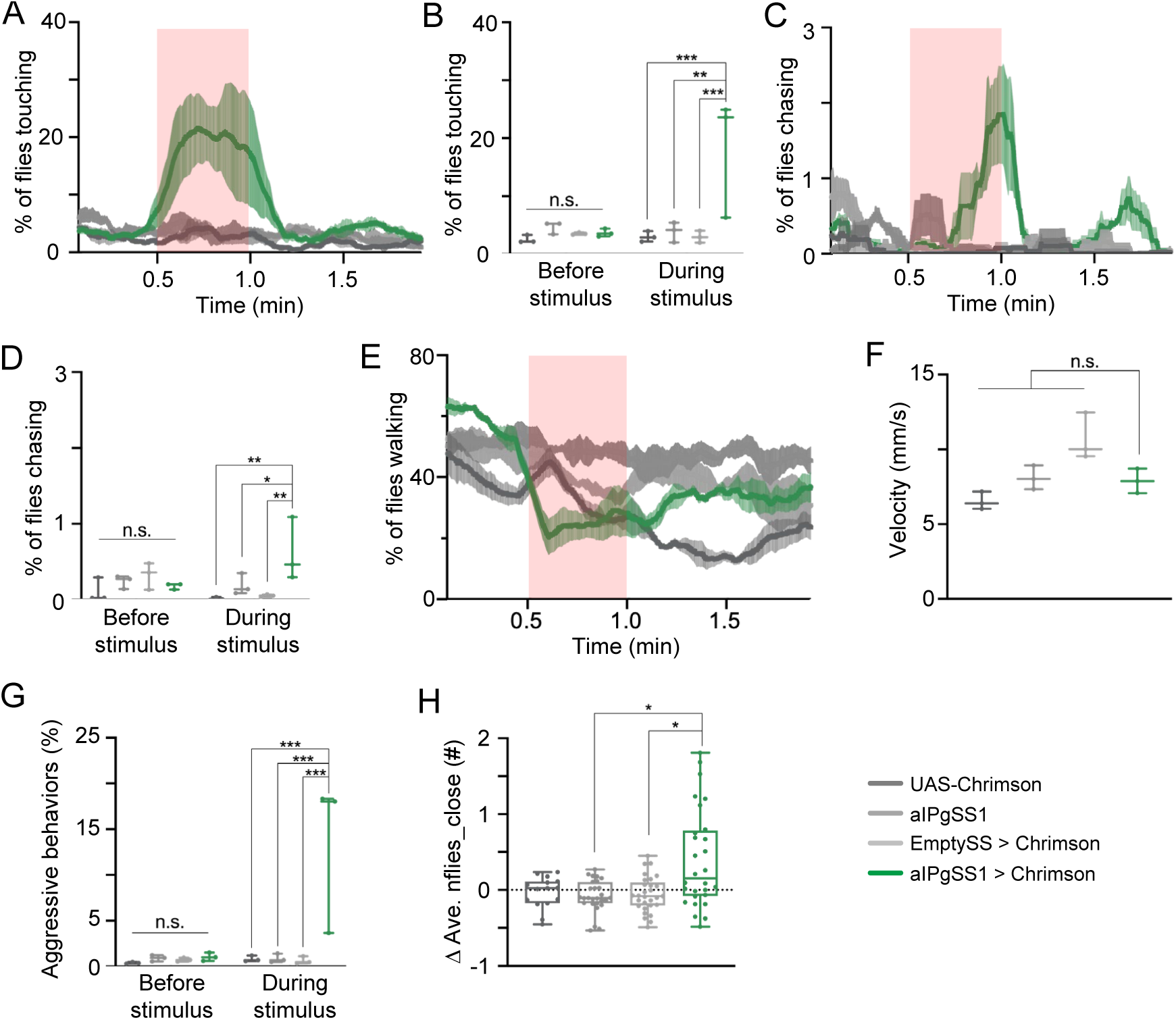
Changes in behavioral metrics in females following activation of aIPgSS1 neurons. (A, C, E) Percentage of female flies touching (A), chasing (C), or walking (E) over the course of a 2-minute trial during which a 30-second 0.4 mW/mm^2^ continuous light stimulus was delivered. (B, D) Average percentage of flies touching (B) or chasing (D) over the 30-second period prior to or during the stimulus delivery in A or C, respectively. (F) Velocity over the course of a 2-minute trial including the stimulus period. (G) Average percentage of flies engaging in aggressive behaviors over the 30-second period prior to or during the stimulus delivery in Figure 1D. (A – G) 20xUAS-CsChrimson, n = 3 experiments; aIPgSS1, n = 3 experiments; EmptySS > 20xUAS-CsChrimson, n = 3 experiments; aIPgSS1 > 20xUAS-CsChrimson, n = 3 experiments. (H) Difference in the average number of flies within two body lengths between the 30-seconds prior to and during the stimulus. 20xUAS-CsChrimson, n = 15 flies; aIPgSS1, n = 27 flies; EmptySS > 20xUAS-CsChrimson, n = 29 flies; aIPgSS1 > 20xUAS-CsChrimson, n = 28 flies. Points represent separate experiments consisting of approximately 15 flies (B, D, E, G) or an individual fly (H). Analyses were performed on the same data set used in Figure 1. Box-and-whisker plots show median and interquartile range (IQR); whiskers show range. A two-way ANOVA with a Tukey’s multiple comparisons post-hoc test was used for statistical analysis. Asterisk indicates significance from 0: *p<0.05; **p<0.01; ***p<0.001; n.s, not significant.

**Figure 1 – figure supplement 8.**
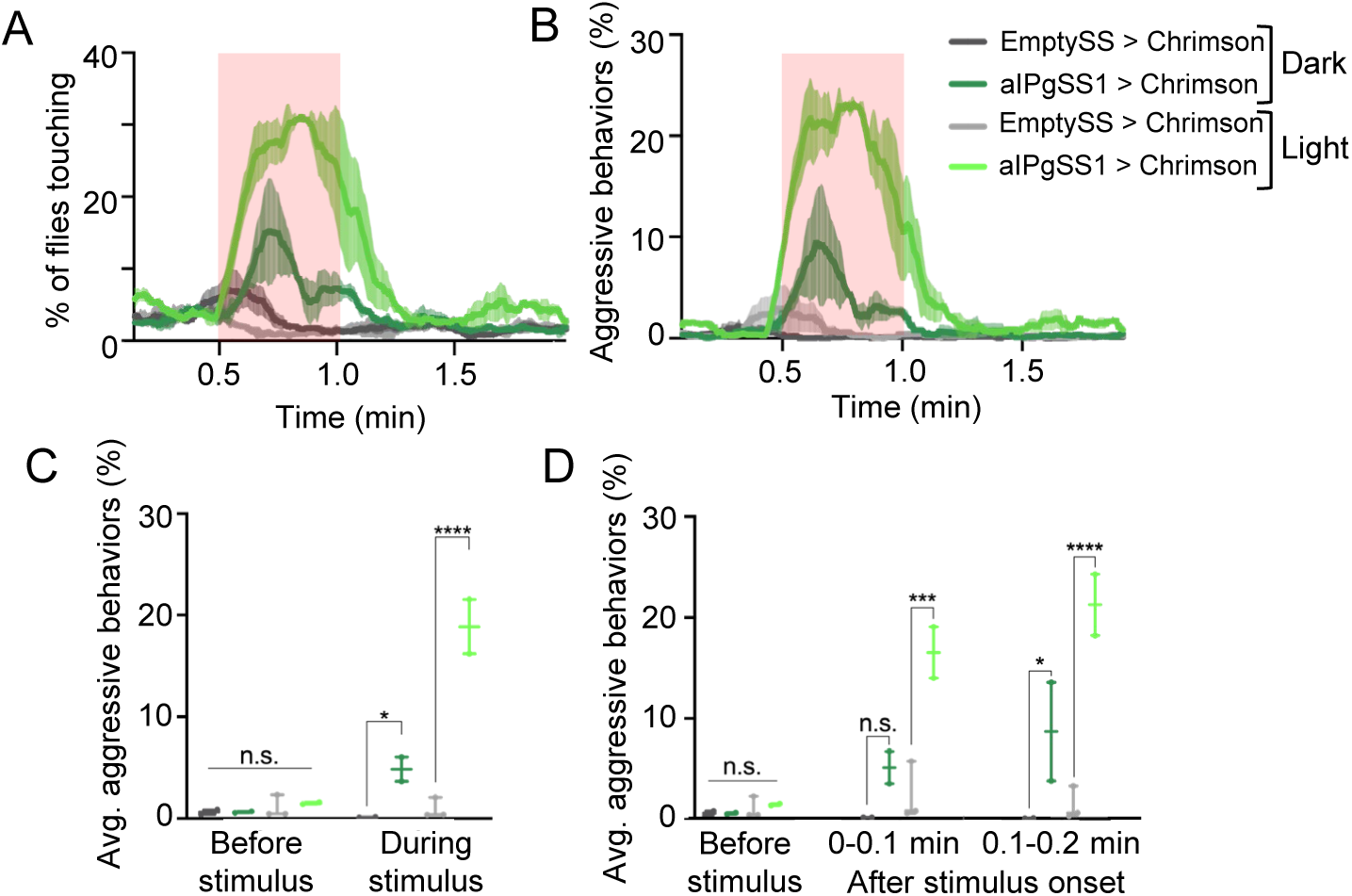
Comparison of the results of aIPgSS1 stimulation in light and dark conditions. (A – B) Percentage of female flies touching (A) or engaging in aggressive behaviors (B) over the course of a 2-minute trial during which a 30-second 0.1 mW/mm^2^ continuous light stimulus was delivered. (C) Average percentage of flies engaging in aggressive behaviors over the 30-second period prior to or during the stimulus delivery in B. (D) Average percentage of flies engaging in aggressive behaviors over the 30-second period prior to, the first 6 seconds, or the second 6 seconds during the stimulus delivery in B. Points in C and D represent separate experiments consisting of approximately 15 flies. Light: EmptySS > Chrimson, n = 3 experiments; aIPgSS1 > Chrimson, n = 2 experiments; Dark: EmptySS > Chrimson, n = 2 experiments; aIPgSS1 > Chrimson, n = 2 experiments. Box-and-whisker plots show median and interquartile range (IQR); whiskers show range. A two-way ANOVA with a Tukey’s multiple comparisons post-hoc test was used for statistical analysis. Asterisk indicates significance from 0: *p<0.05; ***p<0.001; ****p<0.0001; n.s., not significant.

**Figure 1 – figure supplement 9.**
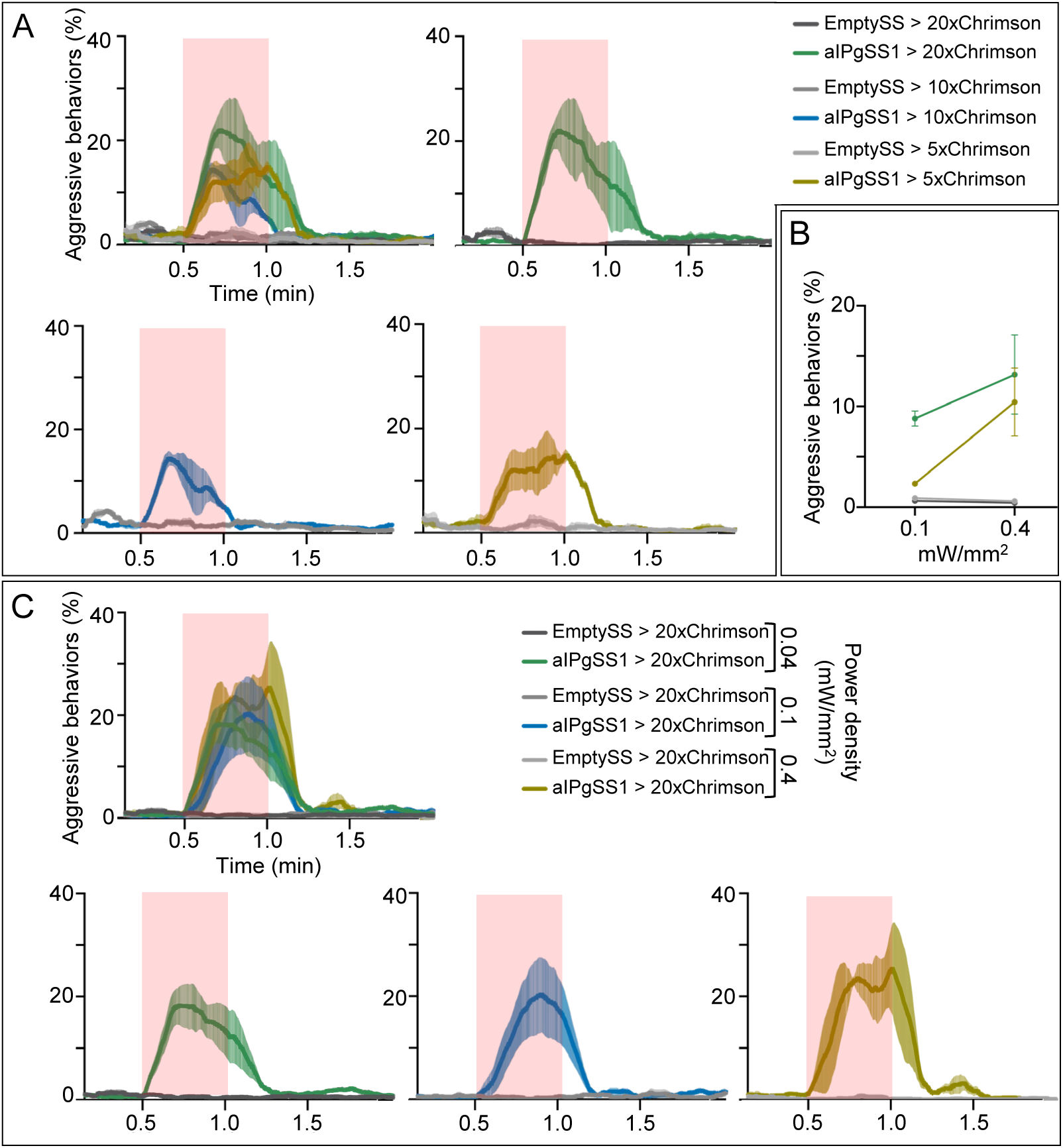
Behavioral effects of effector strength and stimulus delivery. (A) Percentage of flies that carry 5x, 10x, or 20x UAS-Chrimson effectors that engage in aggressive behaviors over the course of a 2-minute trial during which a 30-second 0.1 mW/mm^2^ continuous light stimulus was delivered as indicated by the red shading. Individual curves are shown for comparison and each experiment included approximately 15 flies. EmptySS > 20xUAS-Chrimson, n = 2 experiments; aIPgSS1 > 20xUAS-Chrimson, n = 3 experiments; EmptySS > 10xUAS-Chrimson, n = 3 experiments; aIPgSS1 > 10xUAS-Chrimson, n = 4 experiments; EmptySS > 5xUAS-Chrimson, n = 3 experiments; aIPgSS1 > 5xUAS-Chrimson, n = 3 experiments. (B) Average percentage of flies with either the 5x or 20x UAS-Chrimson effector that perform aggressive behaviors over the 30-second period during the stimulus delivery. Each experiment included approximately 15 flies. 0.1 mW/mm^2^: EmptySS > 5xUAS-Chrimson (light gray), n = 6 experiments; aIPgSS1 > 5xUAS-Chrimson (gold), n = 5 experiments; EmptySS > 20xUAS-Chrimson (dark gray), n = 6 experiments; aIPgSS1 > 20xUAS-Chrimson (green), n = 8 experiments; 0.4 mW/mm^2^: EmptySS > 5xUAS-Chrimson, n = 6 experiments; aIPgSS1 > 5xUAS-Chrimson, n = 9 experiments; EmptySS > 20xUAS-Chrimson, n = 3 experiments; aIPgSS1 > 20xUAS-Chrimson, n = 5 experiments. Bars are mean +/-S.E.M. (C) Percentage of flies engaging in aggressive behaviors over the course of a 2-minute trial during which a 30-second 0.4, 0.1, or 0.04 mW/mm^2^ continuous light stimulus was delivered. Individual curves are shown for comparison and each experiment included approximately 15 flies. 0.04 mW/mm^2^: EmptySS > Chrimson, n = 5 experiments; aIPgSS1 > Chrimson, n = 6 experiments; 0.1 mW/mm^2^: EmptySS > Chrimson, n = 4 experiments; aIPgSS1 > Chrimson, n = 4 experiments; 0.4 mW/mm^2^: EmptySS > Chrimson, n = 3 experiments; aIPgSS1 > Chrimson, n = 3 experiments.

**Figure 1 – figure supplement 10.**
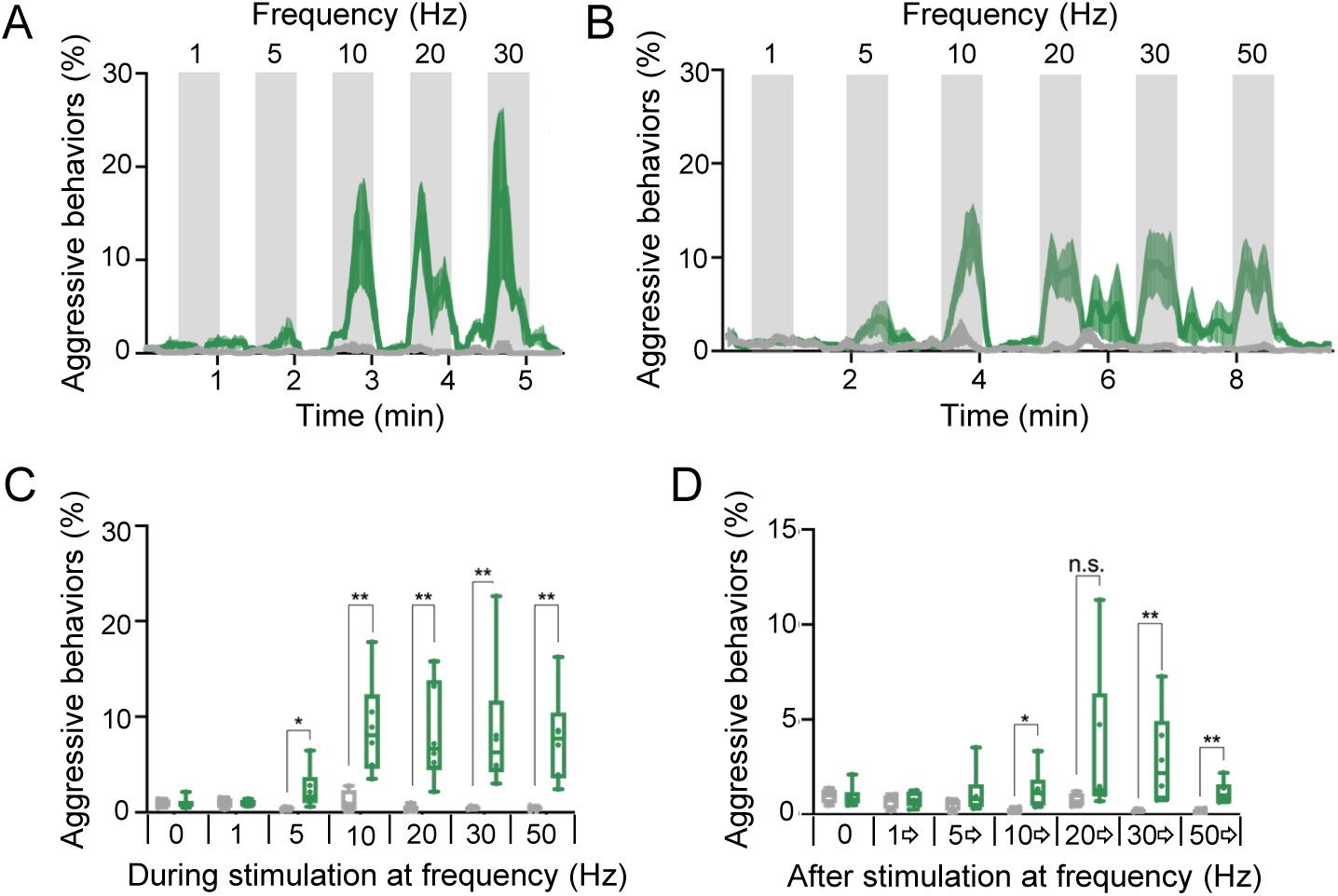
Higher frequency optogenetic stimulation increases the persistence of aggressive behaviors. (A – B) Blocks of 30-second photostimulation (grey bars) with increasing stimulation frequency separated by 30-second (A) or 60-second (B) intervals were delivered sequentially to females. Light was delivered at 0.1mW/mm^2^ with at a 10-ms pulse width, but with increasing pulse frequency. Numbers above the grey bars correspond to the frequency over the 30-second stimulus period. The pulse period and pulse number during each period was as follows: 1000ms, 30; 200ms, 150; 100ms, 300; 50ms, 600; 33ms, 909; 20ms, 1500. aIPgSS1 > 20xUAS-Chrimson is shown in green; EmptySS > 20xUAS-Chrimson is shown in gray. The mean is represented as a solid line and error bars represent variation between experiments. Each experiment included approximately 15 flies. A: EmptySS > 20xUAS-Chrimson, n = 3 experiments; aIPgSS1 > 20xUAS-Chrimson, n = 3 experiments; B: EmptySS > 20xUAS-Chrimson, n = 4 experiments; aIPgSS1 > 20xUAS-Chrimson, n = 5 experiments. (C) Average percentage of flies performing aggressive behaviors over the 30-second period during stimulus delivery in B. (D) Average percentage of flies performing aggressive behaviors over the 30-second period before 1 Hz, the 60-second periods between subsequent stimuli, and the 30-seconds after 50 Hz stimulation in B. Points indicate separate experiments. Box-and-whisker plots show median and interquartile range (IQR); whiskers show range. Mann-Whitney *U*-tests were used for statistical analysis. Asterisk indicates significance from 0: *p<0.05; **p<0.01; n.s., not significant.

**Figure 2 – figure supplement 1.**
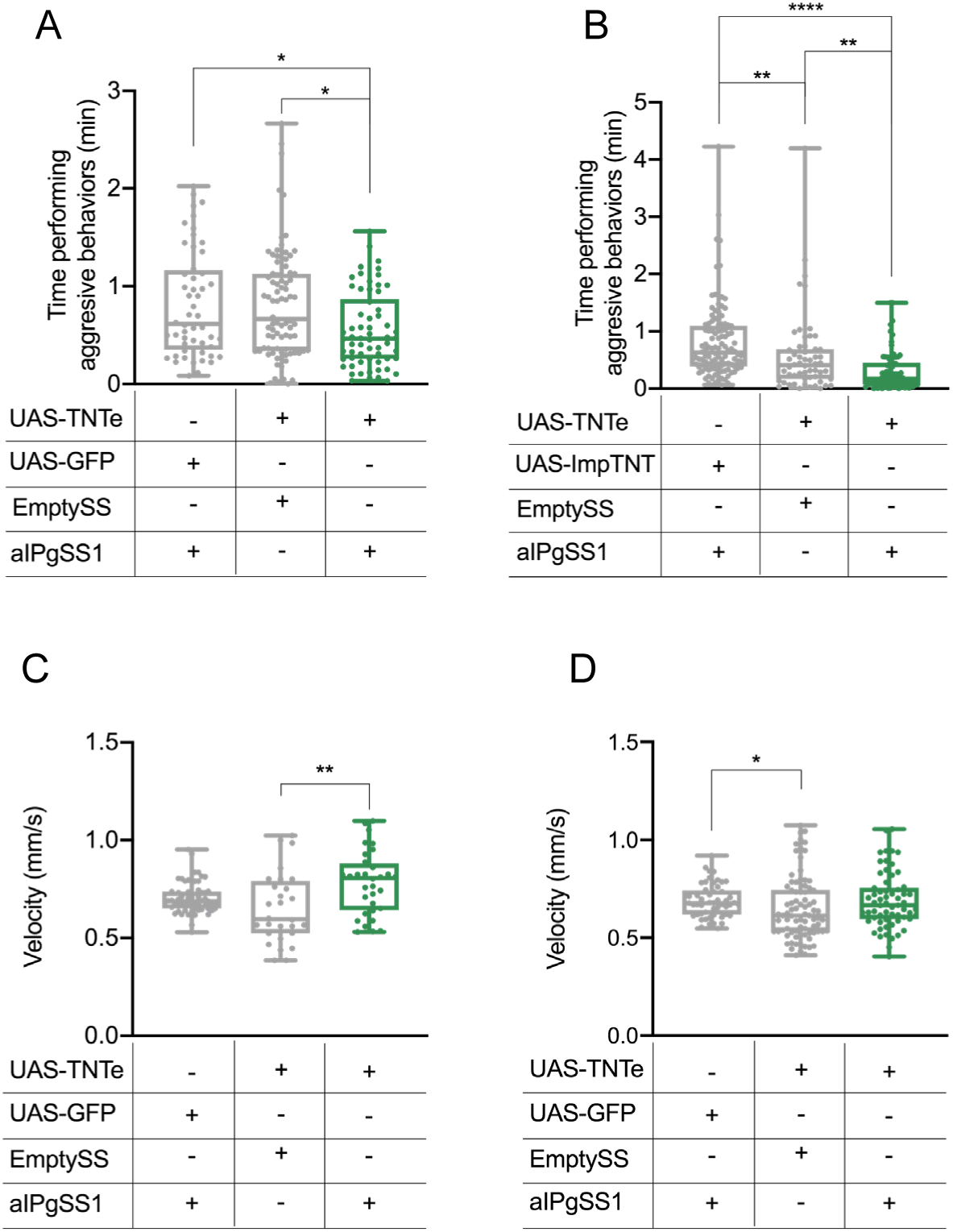
aIPg inactivation reproducibly decreases aggressive behaviors but not velocity. (A – B) Total time an individual spent performing aggressive behaviors over a 30-minute trial. Plots in A and B each represent the pooled data from 4 and 5 testing days, respectively, of separate biological repeats. The experiments in A and B differ in that a different negative control UAS line was used: UAS-GFP in A and UAS-ImpTNT in B; ImpTNT is an inactive form of TNT (see Methods). Experiments were scored using an automated classifier generated with JAABA (see Methods). Points indicate individual flies. A: aIPgSS1 > UAS-GFP, n = 52 flies; EmptySS > UAS-TNTe, n = 82 flies; aIPgSS1 > UAS-TNTe, n = 62 flies; B: aIPgSS1 > UAS-ImpTNTe, n = 110 flies; EmptySS > UAS-TNTe, n = 56 flies; aIPgSS1 > UAS-TNTe, n = 74 flies. (C) Average velocity over the 30-minute trial shown in Figure 2. Points indicate individual flies. aIPgSS1 > UAS-GFP, n = 54 flies; EmptySS > UAS-TNTe, n = 28 flies; aIPgSS1 > UAS-TNTe, n = 30 flies. (D) Average velocity over the 30-minute trial in A. Box-and-whisker plots show median and interquartile range (IQR); whiskers show either 1.5 × IQR of the lower and upper quartiles. Kruskal–Wallis and Dunn’s post hoc tests were used for statistical analysis. Asterisk indicates significance from 0: *p<0.05; **p<0.01; ****p<0.0001.

**Figure 2 – figure supplement 2.**
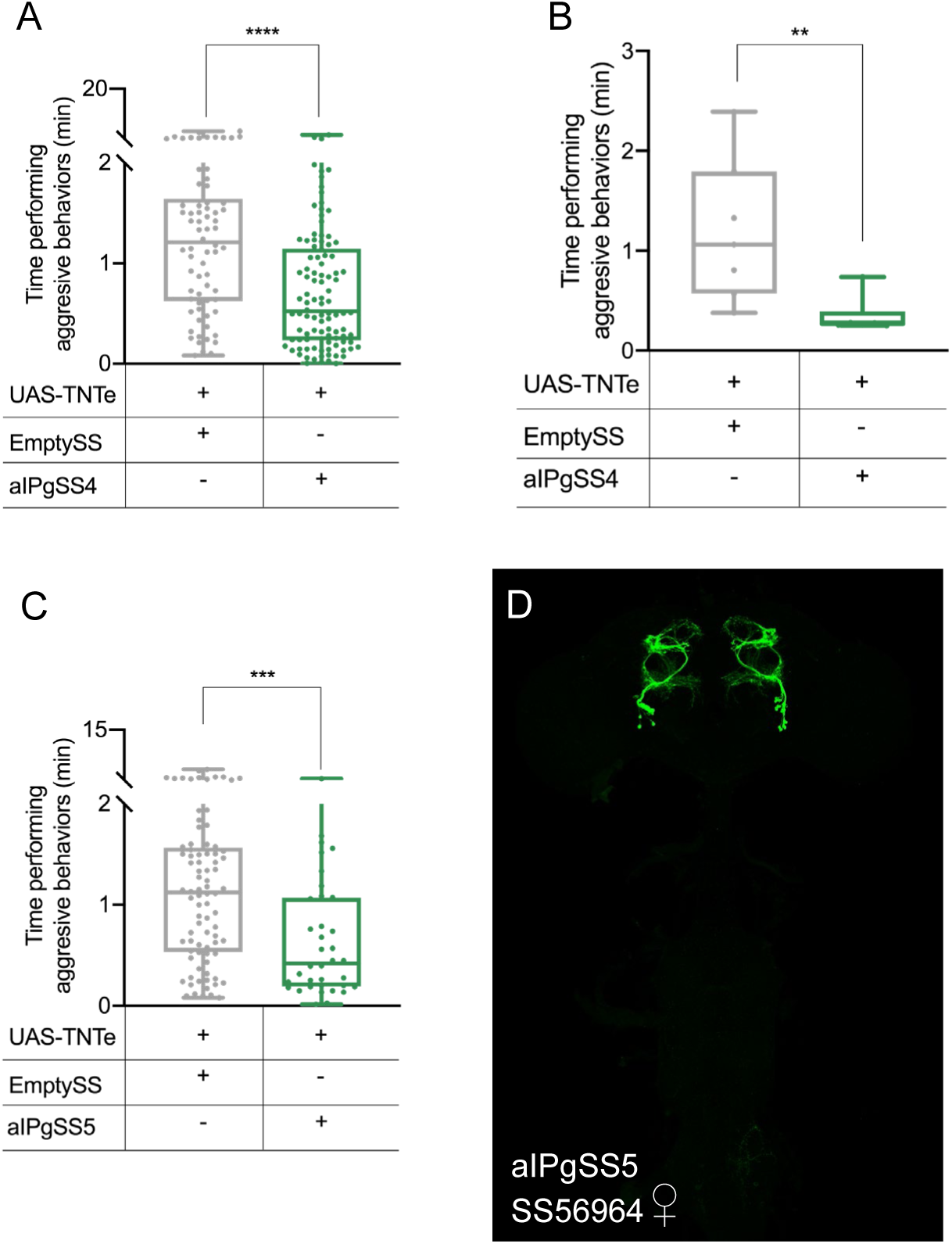
Inactivation of aIPg neurons using additional split-GAL4 lines also decreases aggressive behaviors. (A – C) Total time an individual spent performing aggressive behaviors over a 30-minute trial. Experiments in A-C were pooled from 3, 2 and 2 testing days, respectively. Experiments in B were manually scored with JWatcher software, whereas the experiments in A and C were scored using an automated classifier generated with JAABA (see methods). Points indicate individual flies. A: EmptySS > UAS-TNTe, n = 70 flies; aIPgSS4 > UAS-TNTe, n = 102 flies; B: EmptySS > UAS-TNTe, n = 7 pairs; aIPgSS4 > UAS-TNTe, n = 6 pairs; C: EmptySS > UAS-TNTe, n = 84 flies; aIPgSS5 > UAS-TNTe, n = 36 flies. (D) Expression pattern of SS56964 (aIPgSS5); this line expresses in 15.8±0.4 cells per female brain hemisphere (n = 6). Box-and-whisker plots show median and interquartile range (IQR); whiskers show either 1.5 × IQR of the lower and upper quartiles. Kruskal– Wallis and Dunn’s post hoc tests were used for statistical analysis. Asterisk indicates significance from 0: ***p<0.001; ****p<0.0001.

**Figure 3 – figure supplement 1.**
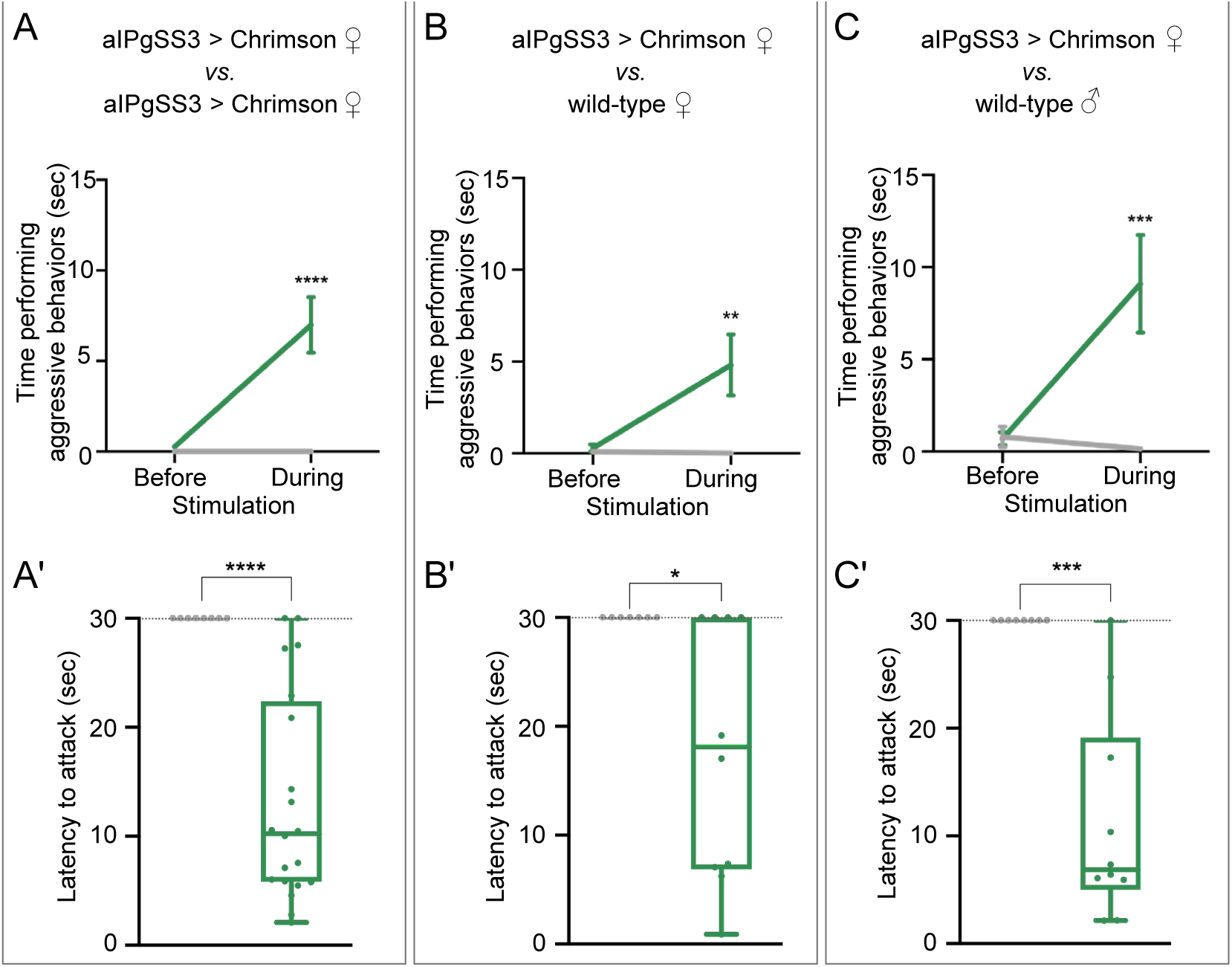
Increased aggression against wild-type females and males reproduced with a second aIPgSS line. (A – C) Total time an individual spent performing aggressive behaviors in a 16 mm arena over the 30 second period prior to or during stimulation. The plots refer only to the behavior of aIPgSS3 > Chrimson females and each arena contained only two flies: (A) Two aIPgSS3 > Chrimson females; (B) an aIPgSS3 > Chrimson female and a wild-type (Canton-S) female; and (C) a aIPgSS3 > Chrimson female and a wild-type (Canton-S) male. The green line shows the stated genotype; the gray line shows the results when EmptySS > Chrimson was used instead of aIPgSS3 > Chrimson. (A’ – C’) Amount of time during a 30 second 0.1 mW/mm^2^ continuous stimulation period until first aggressive encounter. Points indicate individual flies. Dotted lines indicate the end of the trial and error bars are mean +/-S.E.M. Box-and-whisker plots show median and interquartile range (IQR); whiskers show either 1.5 × IQR of the lower and upper quartiles. A’: EmptySS > 20xUAS-Chrimson, n = 16 flies; aIPgSS3 > 20xUAS-Chrimson, n = 18 flies; B’: EmptySS > 20xUAS-Chrimson, n = 5 flies; aIPgSS3 > 20xUAS-Chrimson, n = 7 flies; C’: EmptySS > 20xUAS-Chrimson, n = 8 flies; aIPgSS3 > 20xUAS-Chrimson, n = 10 flies. A Mann-Whitney *U* (A’ – C’) post hoc test or a two-way ANOVA with a Sidak’s multiple comparisons post-hoc test (A – C) was used for statistical analysis. Asterisk indicates significance from 0: *p<0.05; **p<0.01; ****p<0.0001; n.s., not significant.

**Figure 3 – figure supplement 2.**
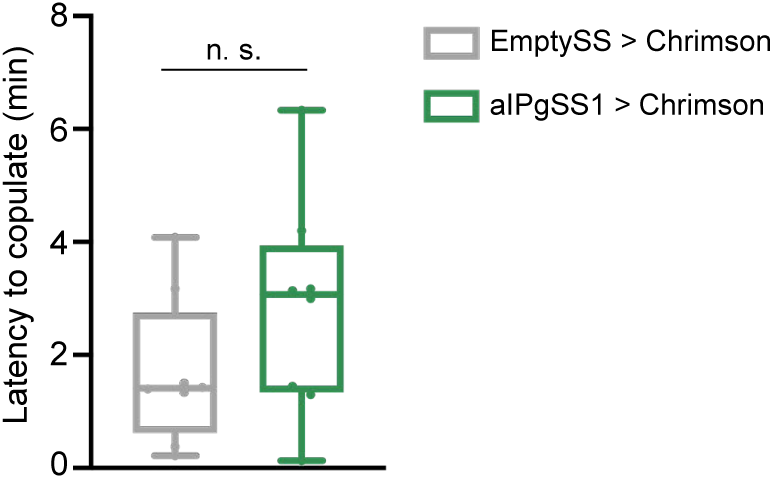
aIPg activation does not significantly alter copulation latency. Amount of time until first copulation event following a 0.4 mW/mm^2^ continuous stimulus. Virgin females were added into a 10 mm arena containing virgin wild type (Canton-S) males immediately prior to the stimulus onset with each stimulus lasting 10 minutes. Points indicate individual flies. EmptySS > 20xUAS-Chrimson, n = 8 flies; aIPgSS1 > 20xUAS-Chrimson, n = 8 flies. Box-and-whisker plots show median and interquartile range (IQR); whiskers show either 1.5 × IQR of the lower and upper quartiles. A Mann-Whitney *U* post hoc test was used for statistical analysis. n.s., not significant.

**Figure 4 – figure supplement 1.**
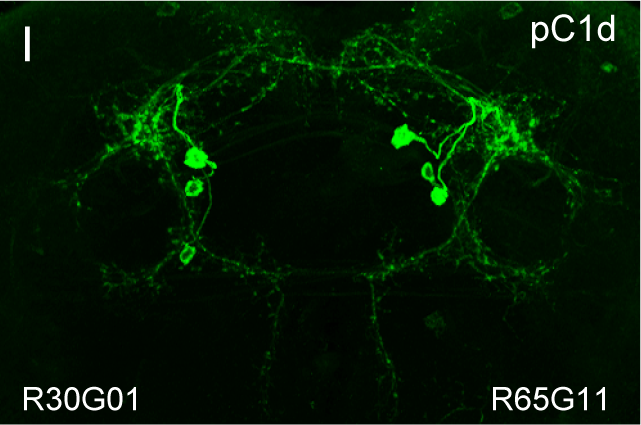
(A – I) Each panel shows the intersection of the indicated enhancers; the enhancer in the lower left was used to drive the split-GAL4 AD domain and the enhancer in the lower right drove the DBD domain. We observed either the pC1d or aIPg cell type in each intersection. Nearly all of the pC1d containing intersections also show expression in pC1e or other members of the pC1 cell type group.

**Figure 5 – figure supplement 1.**
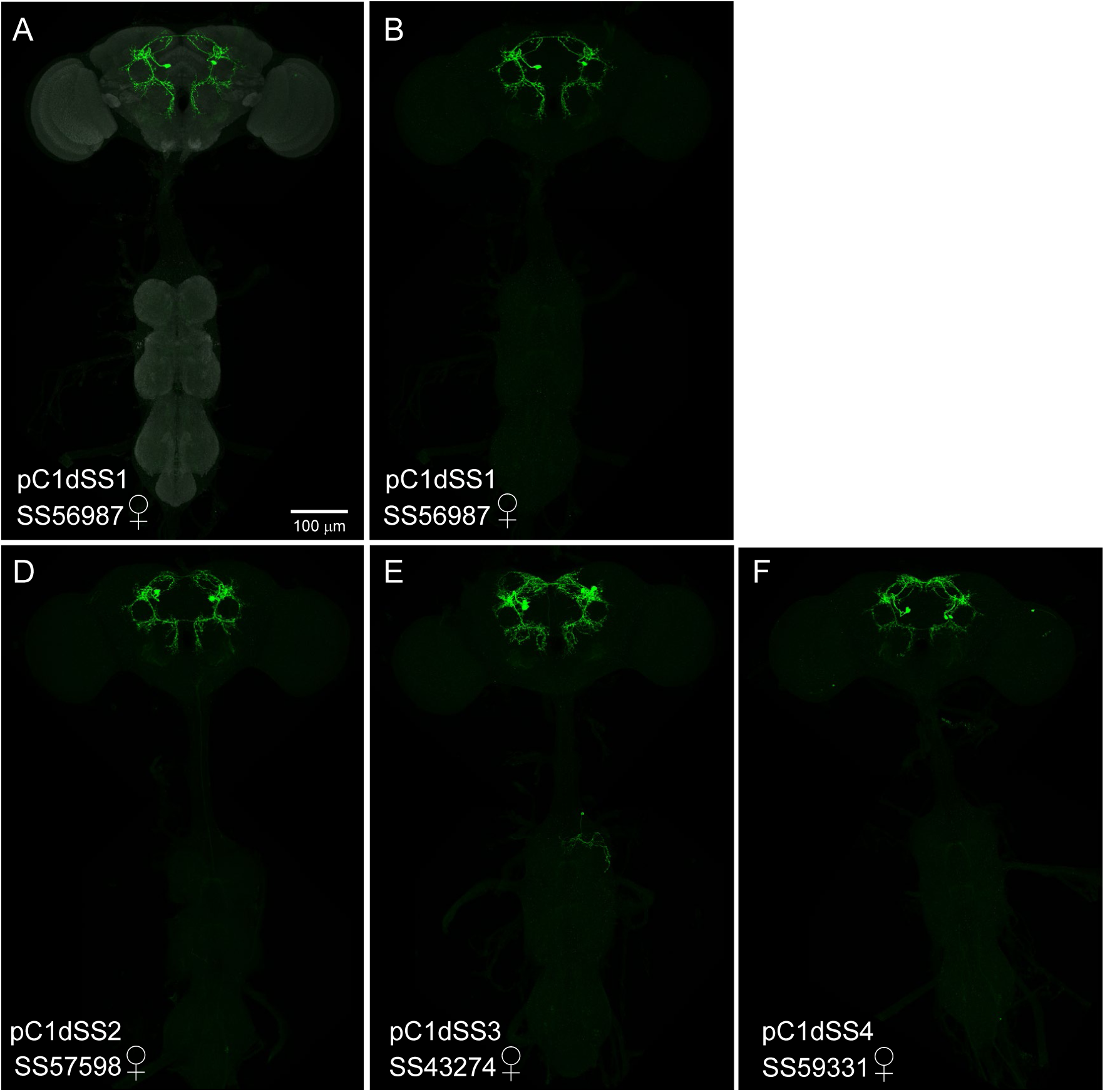
Expression patterns of pC1d split-GAL4 lines. (A – F) 20X maximum intensity projections of the brains and ventral nerve cords of the indicated split-GAL4 lines crossed to 20xUAS-CsChrimson::mVenus. As we are visualizing the optogenetic effector itself, these images serve as a way to compare the relative expression levels of the effector in the different split-GAL4 lines. Sex of the imaged brain is indicated. The scale bar shown in A applies to panels A – F. (D’ – F’) Enlargements of the central brain of the images shown in D – F. (A – C) The SS56987 (pC1dSS1) expression pattern is shown; the neuropil reference channel shown in A (gray). Note that no expression in pC1d neurons is seen in the male nervous system which was imaged under identical conditions to the female nervous system shown in B. (D, D’) Expression pattern of SS57598 (pC1dSS2). (E, E’) Expression pattern of SS43274 (pC1dSS3). Note that this line expresses in both pC1d and pC1e. This line shows expression in males in a cell type clearly distinct from pC1d. Arrows indicate cell bodies. (F, F’). Expression pattern of SS59331 (pC1dSS4). Note that this line expresses in both pC1d and pC1e. Arrows indicate cell bodies.

**Figure 5 – figure supplement 2.**
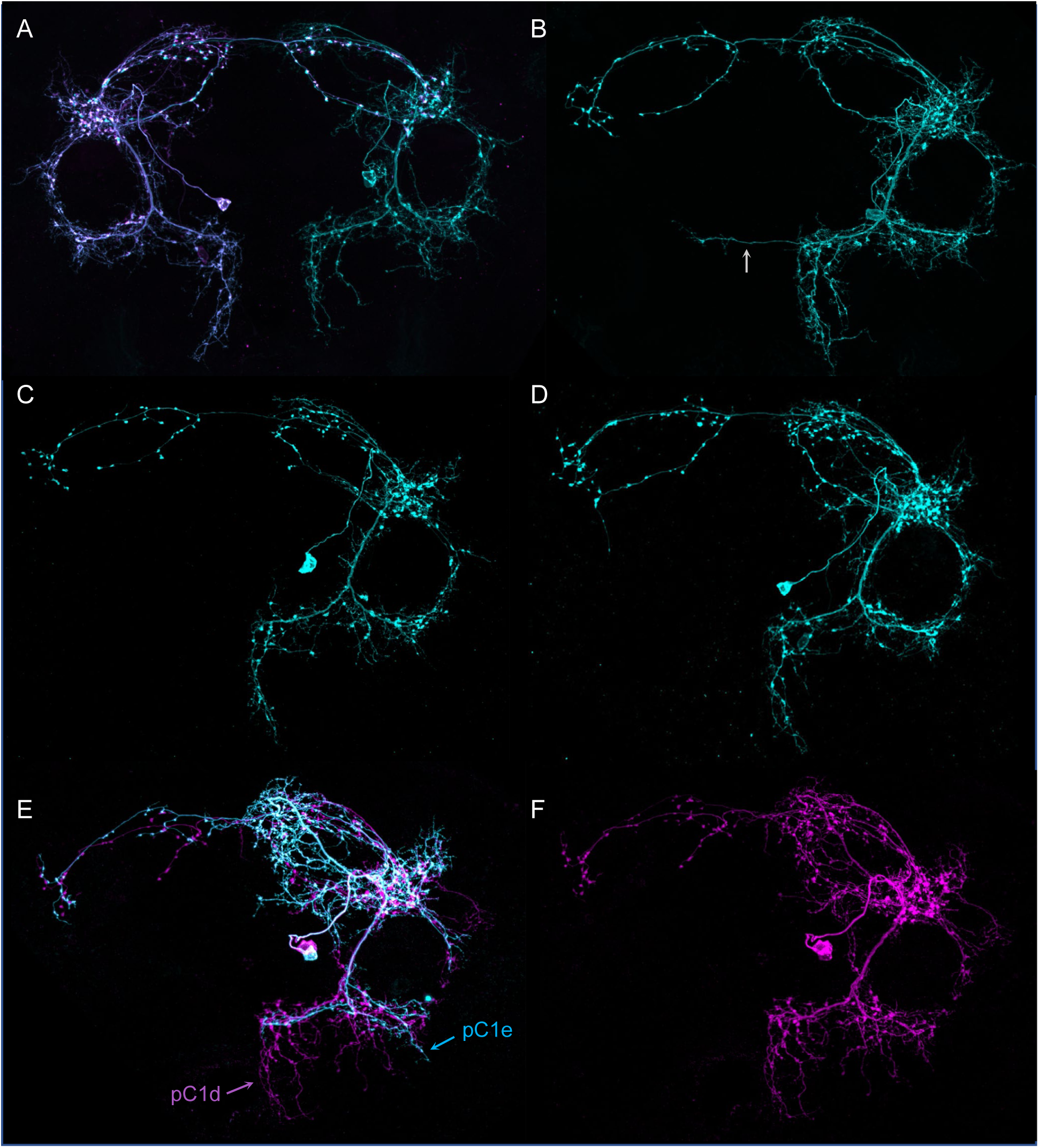
Morphologies of individual pC1d neurons. The images in all panels were generated by stochastic labeling of pC1d split-GAL4 lines using the MultiColor FlipOut method (Nern et al. 2015). Both brain hemispheres are shown. (A – D) Images from SS56987 (pC1dSS1). The arrow in (B) highlights a process that crosses the midline which is often seen in pC1d cells; see (D’) and (F’) in Figure 5 for additional examples. (A) The right and left hemisphere pC1d cells are shown. (E, F) Images from SS43274 (pC1dSS3).

**Figure 5 – figure supplement 3.**
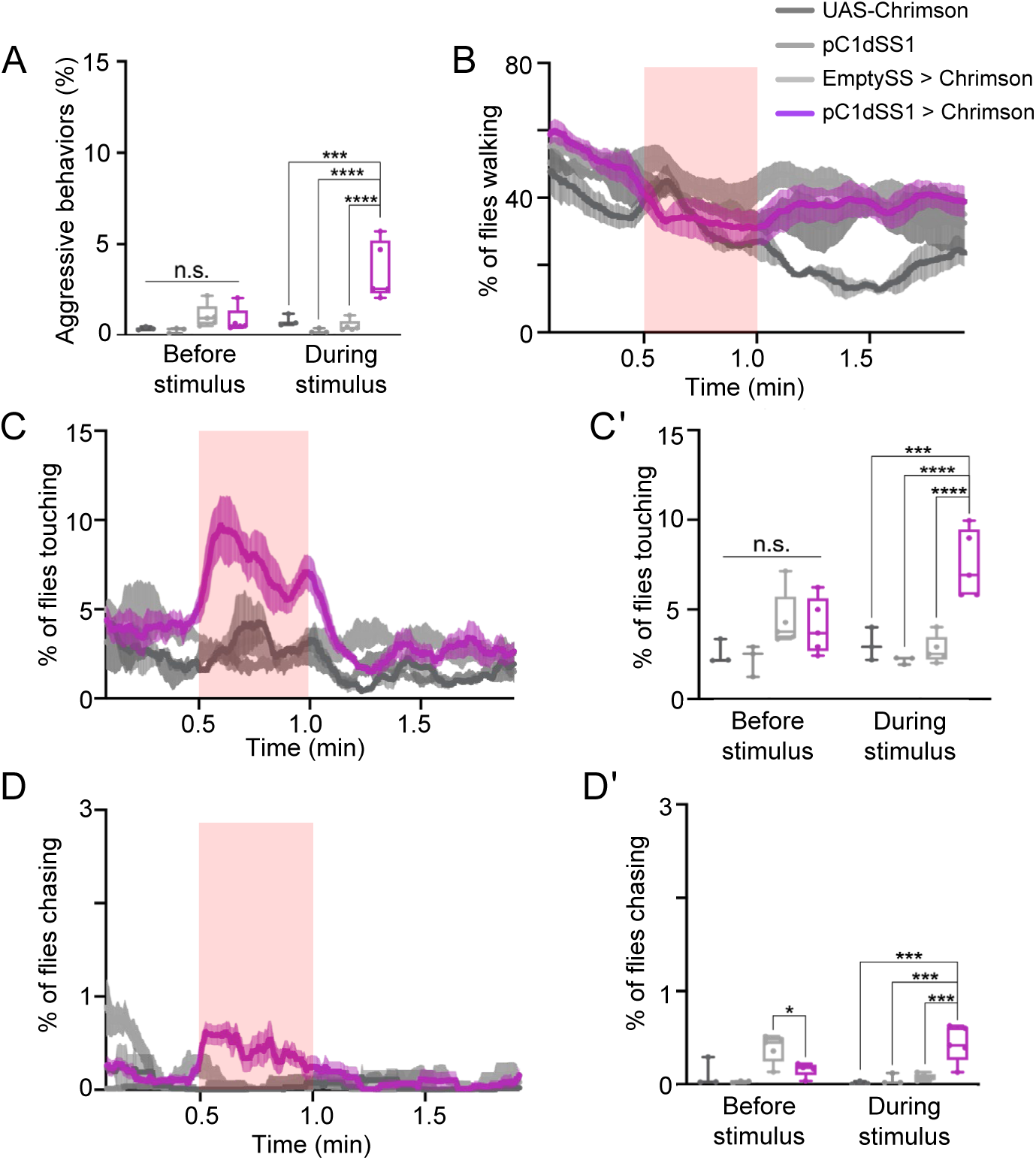
Behavioral characterization of female flies after pC1d activation. Behaviors were measured over the course of a 2-minute trial during which a 30-second 0.4 mW/mm^2^ continuous light stimulus was delivered (A) average percentage of flies engaging in aggressive behaviors over the 30-second period prior to or during stimulus delivery. (B, C, D) Percentage of female flies walking (B), touching (C), or chasing (D). (C’, D’) Average percentage of flies touching (C’) or chasing (D’) over the 30-second period prior to or during the stimulus delivery in C or D, respectively. Points represent separate experiments consisting of approximately 15 flies. 20xUAS-CsChrimson, n = 3 experiments; pC1dSS1, n = 3 experiments; EmptySS > 20xUAS-CsChrimson, n = 5 experiments; pC1dSS1 > 20xUAS-CsChrimson, n = 5 experiments. Box-and-whisker plots show median and interquartile range (IQR); whiskers show range. A two-way ANOVA with a Tukey’s multiple comparisons post-hoc test was used for statistical analysis. Asterisk indicates significance from 0: *p<0.05; ***p<0.001; ****p<0.0001; n.s., not significant.

**Figure 5 – figure supplement 4.**
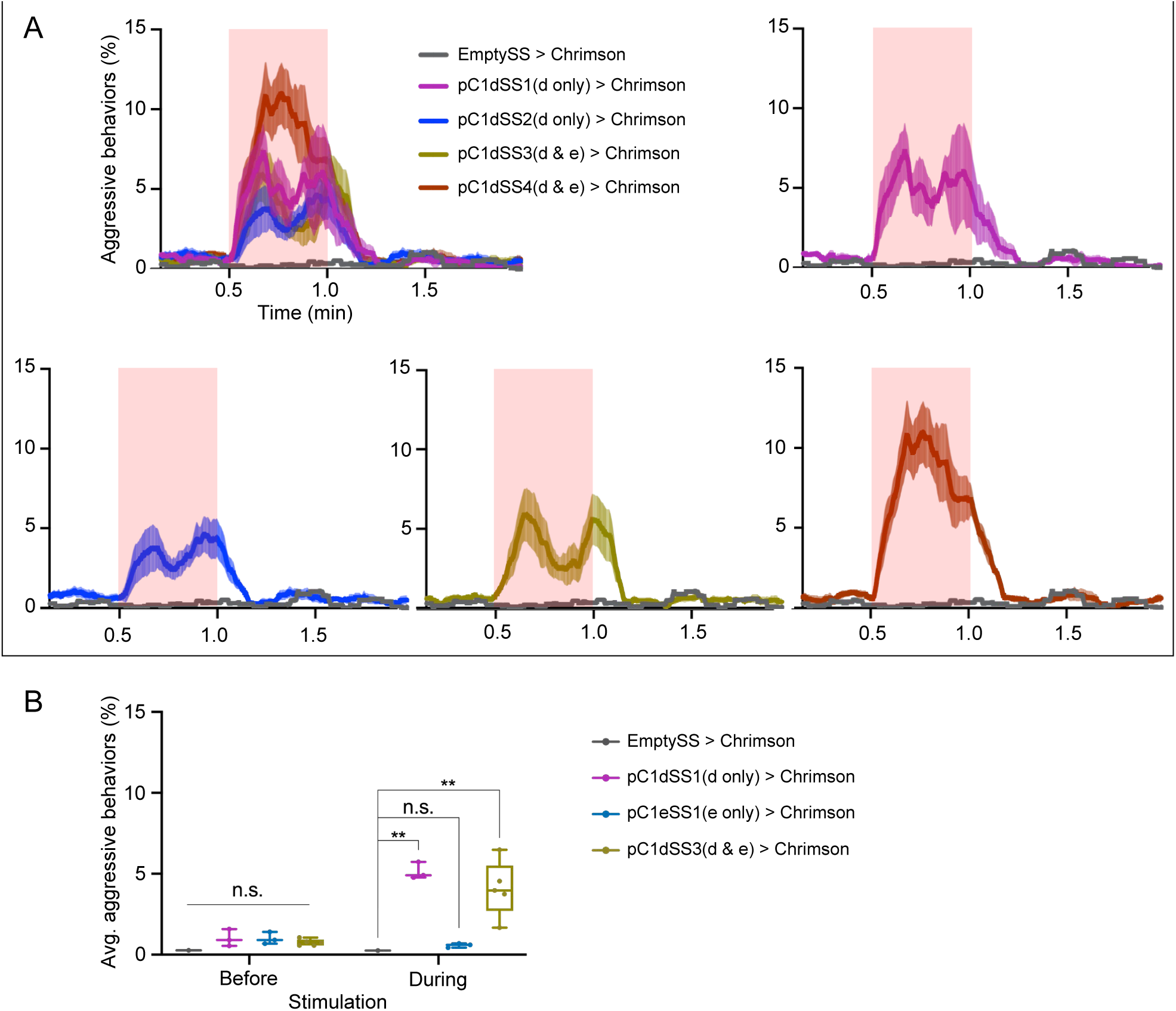
Optogenetic activation of additional lines labeling pC1d split-GAL4 lines display similar behavioral results to pC1dSS1. (A) Percentage of flies engaging in aggressive behaviors over the course of a 2-minute trial during which a 30-second 0.1 mW/mm^2^ continuous light stimulus was delivered. Red shading indicates the stimulus period. The mean is represented as a solid line and error bars represent variation between experiments. Note that two pC1d split-GAL4 lines label just pC1d and two others label both pC1d and pC1e, as indicated. Individual curves are shown for comparison. Each experiment included approximately 15 flies. EmptySS > Chrimson, n = 1 experiment; pC1dSS1 > Chrimson, n = 3 experiments; pC1dSS2 > Chrimson, n = 6 experiments; pC1dSS3 > Chrimson, n = 5 experiments; pC1dSS4 > Chrimson, n = 5 experiments. (B) Average percentage of flies engaging in aggressive behaviors over the 30 second period prior to or during the delivery of a 0.1 mW/mm^2^ stimulus. Points represent separate experiments consisting of approximately 15 flies. EmptySS > Chrimson, n = 1 experiment; pC1dSS1 > Chrimson, n = 3 experiments; pC1eSS1 > Chrimson, n = 3 experiments; pC1dSS3 > Chrimson, n = 5 experiments. Box-and-whisker plots show median and interquartile range (IQR); whiskers show range. A two-way ANOVA with a Tukey’s multiple comparisons post-hoc test was used for statistical analysis. Asterisk indicates significance from 0: **p<0.01; n.s., not significant.

**Figure 5 – figure supplement 5.**
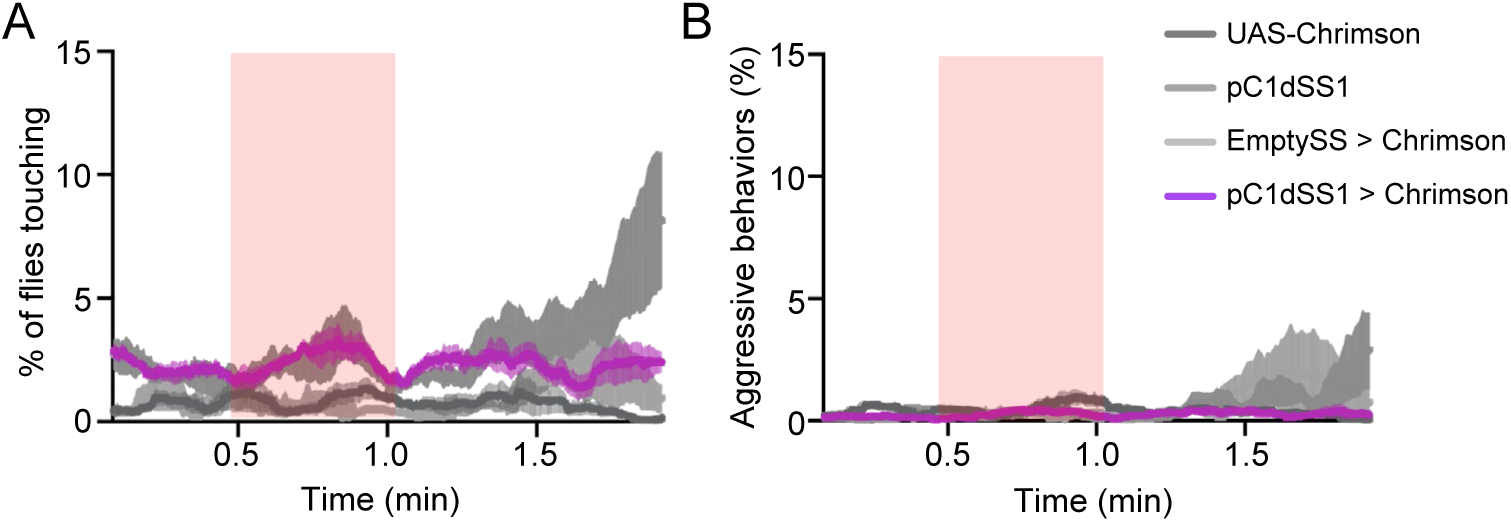
Optogenetic stimulation of pC1dSS1>Chrimson males does not result in aggressive behavior. (A – B) Percentage of male flies touching (A) or performing aggressive behaviors (B) over the course of a 2-minute trial during which a 30 second 0.4 mW/mm^2^ continuous light stimulus was delivered. The mean is represented as a solid line and error bars represent variation between experiments. Each experiment included approximately 15 flies. 20xUAS-CsChrimson, n = 5 experiments; pC1dSS1, n = 4 experiments; EmptySS > 20xUAS-CsChrimson, n = 3 experiments; pC1dSS1> 20xUAS-CsChrimson, n = 5 experiments.

**Figure 5 – figure supplement 6.**
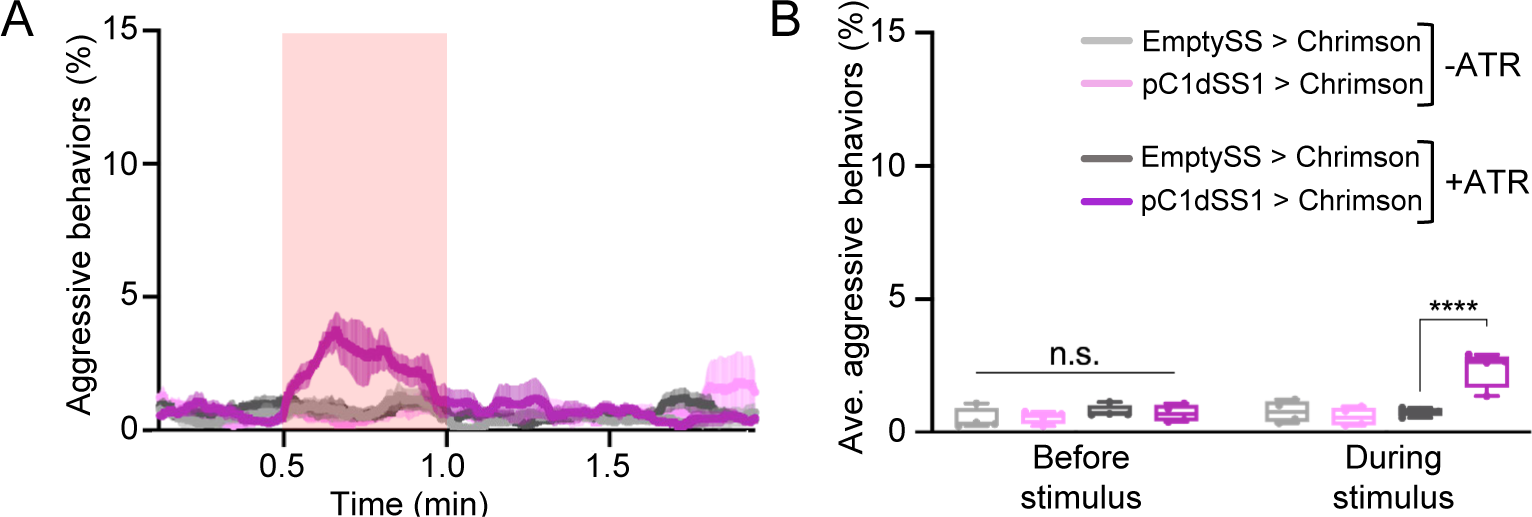
Optogenetic activation of aggression depends on feeding all *trans*-retinal. (A) Percentage of flies that engaged in aggressive behaviors. Flies were raised with food supplemented with all *trans*-retinal (+ATR) or not (-ATR). A 30-second 0.1 mW/mm^2^ continuous light stimulus was delivered 30 sec after the start of a 2-minute trial as indicated by the red shading. The mean is represented as a solid line and error bars represent variation between experiments. Each experiment included approximately 15 flies. (B) Average percentage of flies engaging in aggressive behaviors over the 30-second period prior to or during the stimulus delivery in A. Points represent separate experiments consisting of approximately 15 flies. +ATR: EmptySS > Chrimson, n = 4 experiments; pC1dSS1 > Chrimson, n = 4 experiments; -ATR: EmptySS > Chrimson, n = 4 experiments; pC1dSS1> Chrimson, n = 4 experiments. Box-and-whisker plots show median and interquartile range (IQR); whiskers show range. A two-way ANOVA with a Tukey’s multiple comparisons post hoc test was used for statistical analysis. Asterisk indicates significance from 0: ****p<0.0001.

**Figure 5 – figure supplement 7.**
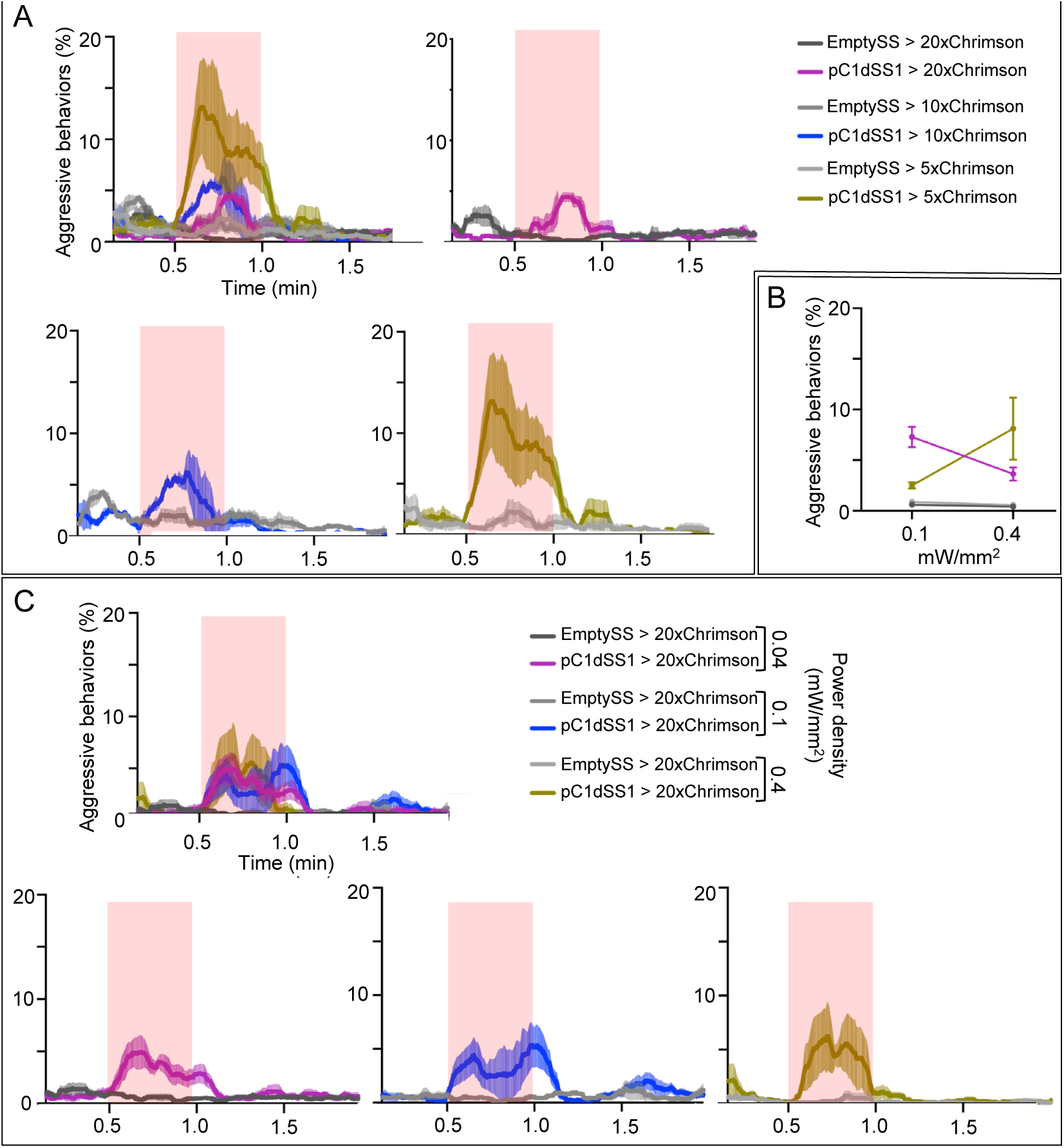
Behavioral effects of stimulus delivery and effector strength. (A) Percentage of flies engaging in aggressive behaviors over the course of a 2-minute trial during which a 30-second 0.1 mW/mm^2^ continuous light stimulus was delivered to flies carrying 5XUAS, 10xUAS, and 20xUAS effectors. Individual curves are shown for comparison and each experiment included approximately 15 flies. EmptySS > 20xUAS-Chrimson, n = 2 experiments; pC1dSS1 > 20xUAS-Chrimson, n = 2 experiments; EmptySS > 10xUAS-Chrimson, n = 2 experiments; pC1dSS1 > 10xUAS-Chrimson, n = 2 experiments; EmptySS > 5xUAS-Chrimson, n = 3 experiments; pC1dSS1 > 5xUAS-Chrimson, n = 3 experiments. (B) Average percentage of flies engaging in aggressive behaviors over the 30-second period during stimulus delivery. Each experiment included approximately 15 flies. 0.1 mW/mm^2^: EmptySS > 5xUAS-Chrimson (light gray), n = 6 experiments; pC1dSS1 > 5xUAS-Chrimson (gold), n = 6 experiments; EmptySS > 20xUAS-Chrimson (dark gray), n = 6 experiments; pC1dSS1 > 20xUAS-Chrimson (purple), n = 7 experiments; 0.4 mW/mm^2^: EmptySS > 5xUAS-Chrimson, n = 6 experiments; pC1dSS1 > 5xUAS-Chrimson, n = 2 experiments; EmptySS > 20xUAS-Chrimson, n = 3 experiments; pC1dSS1 > 20xUAS-Chrimson, n = 5 experiments. Bars are mean +/- S.E.M. (C) Percentage of flies engaging in aggressive behaviors over the course of a 2-minute trial during which a 30-second 0.4, 0.1, or 0.04 mW/mm^2^ continuous light stimulus was delivered. Individual curves are shown for comparison and each experiment included approximately 15 flies. 0.04 mW/mm^2^: EmptySS > Chrimson, n = 5 experiments; pC1dSS1 > Chrimson, n = 5 experiments; 0.1 mW/mm^2^: EmptySS > Chrimson, n = 4 experiments; pC1dSS1 > Chrimson, n = 4 experiments; 0.4 mW/mm^2^: EmptySS > Chrimson, n = 3 experiments; pC1dSS1 > Chrimson, n = 3 experiments.

**Figure 5 – figure supplement 8.**
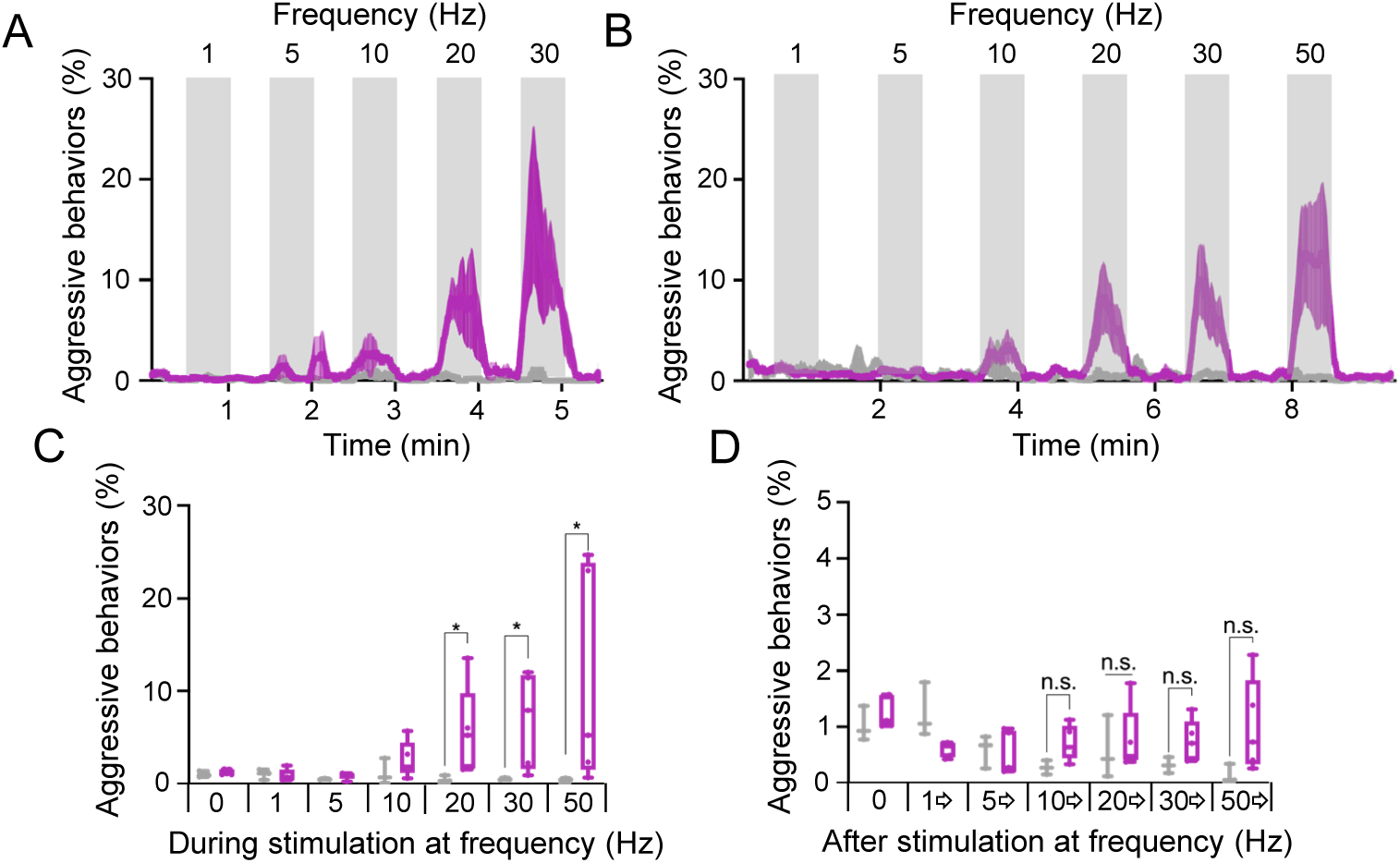
Behavioral effects of the frequency of optogenetic stimulation. (A – B) Blocks of 30-second photostimulation (grey bars) with increasing stimulation frequency separated by 30 (A) or 60 (B) second intervals were delivered sequentially to pC1dSS1females. Light was delivered at 0.1mW/mm^2^ with at a 10-ms pulse width, but with increasing pulse frequency. Numbers above the grey bars correspond to frequency over the 30-second stimulus period. The pulse period and pulse number during each period was as follows: 1000ms, 30; 200ms, 150; 100ms, 300; 50ms, 600; 33ms, 909; 20ms, 1500. pC1dSS1 > 20xUAS-Chrimson is shown in purple; EmptySS > 20xUAS-Chrimson is shown in gray. The mean is represented as a solid line and shaded error bars represent variation between experiments. Each experiment included approximately 15 flies. A: EmptySS > 20xUAS-Chrimson, n = 3 experiments; pC1dSS1 > 20xUAS-Chrimson, n = 3 experiments; B: EmptySS > 20xUAS-Chrimson, n = 3 experiments; pC1dSS1 > 20xUAS-Chrimson, n = 5 experiments. (C) Average percentage of flies performing aggressive behaviors over the 30-second period during stimulus delivery in B. (D) Average percentage of flies performing aggressive behaviors over the 30-second period before 1 Hz, the 60-second periods between subsequent stimuli, and the 30-seconds after 50 Hz stimulation in B. Points indicate separate experiments. Box-and-whisker plots show median and interquartile range (IQR); whiskers show range. Mann-Whitney *U*-tests were used for statistical analysis. Asterisk indicates significance from 0: *p<0.05; n.s., not significant.

**Figure 5 – figure supplement 9.**
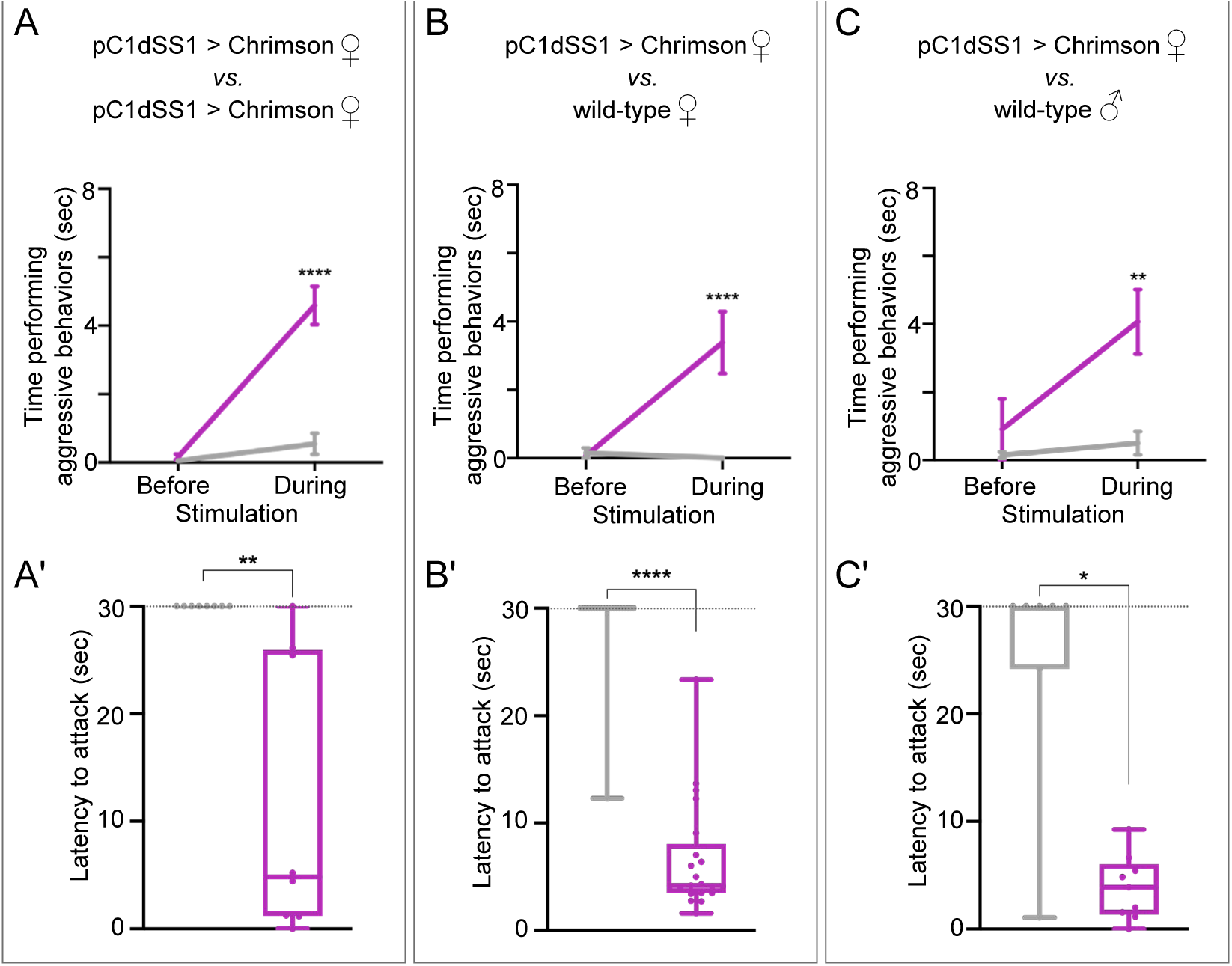
pC1d activation also increases aggression against wild-type females and males. (A – C) Total time an individual spent performing aggressive behaviors over the 30 second period prior to or during stimulation, see Figure 3 legend for detail. (A’ – C’) Amount of time during a 30 second 0.1 mW/mm^2^ continuous stimulation period until first aggressive encounter. Points indicate individual flies. Dotted lines at 30 sec indicate the end of the trial and error bars are mean +/- S.E.M. A: EmptySS > 20xUAS-Chrimson, n = 22 flies; pC1dSS1 > 20xUAS-Chrimson, n = 22 flies; B: EmptySS > 20xUAS-Chrimson, n = 8 flies; pC1dSS1 > 20xUAS-Chrimson, n = 8 flies; C: EmptySS > 20xUAS-Chrimson, n = 7 flies; pC1dSS1 > 20xUAS-Chrimson, n = 8 flies. Box-and-whisker plots show median and interquartile range (IQR); whiskers show either 1.5 × IQR of the lower and upper quartiles. A two-way ANOVA with a Sidak’s multiple comparisons post-hoc test (A – C) or Mann-Whitney *U* (A’ – C’) post hoc test was used for statistical analysis. Asterisk indicates significance from 0: *p<0.05; **p<0.01; ****p<0.0001.

**Figure 5 – figure supplement 10.**
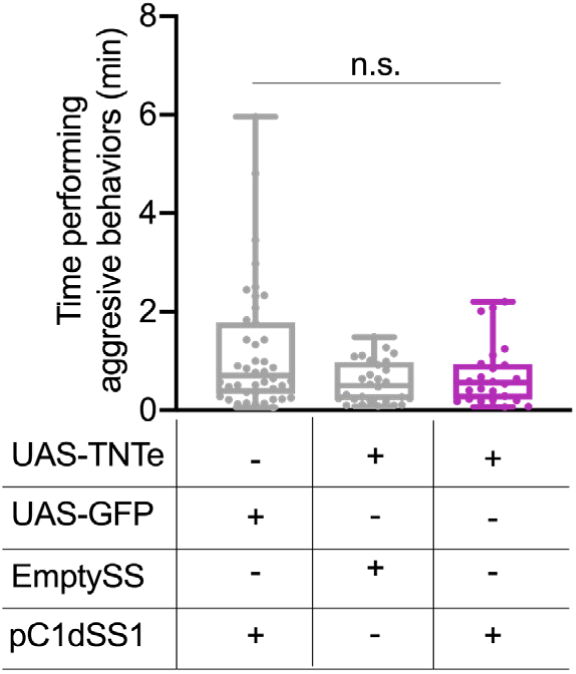
pC1d inactivation did not significantly diminish aggressive behavior. Total time an individual spent performing aggressive behaviors over a 30-minute trial. Points indicate individual flies. pC1dSS1 > UAS-GFP, n = 42 flies; EmptySS > UAS-TNTe, n = 28 flies; pC1dSS1 > UAS-TNTe, n = 24 flies. Box-and-whisker plots show median and interquartile range (IQR); whiskers show either 1.5 × IQR of the lower and upper quartiles. Kruskal–Wallis and Dunn’s post hoc tests were used for statistical analysis. n.s., not significant.

**Figure 6 – figure supplement 1.**
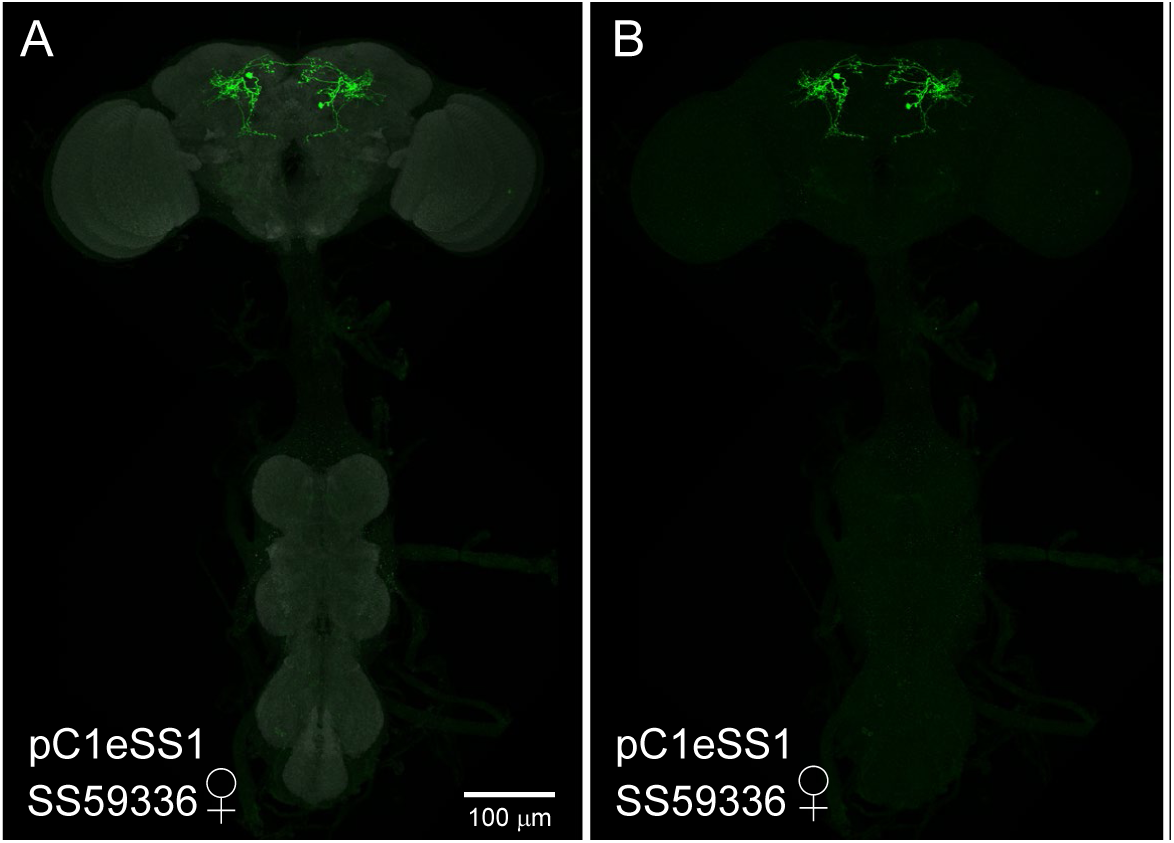
Expression patterns of pC1e split-GAL4 lines. (A – E) 20X maximum intensity projections of the brains and ventral nerve cords of the indicated split-GAL4 lines crossed to 20xUAS-CsChrimson::mVenus. Because we are visualizing the optogenetic effector itself, these images serve as a way to compare the relative expression levels of the effector in the different split-GAL4 lines. The gender of the imaged brain is indicated. The scale bar shown in A applies to panels A-E. (D’, E’) Enlargements of the central brain of the images shown in D and E. (A – C) The SS59336 (pC1eSS1) expression pattern is shown; the neuropil reference channel is shown in A (gray). Note that no expression in aIPg neurons is seen in the male nervous system (C) which was imaged under identical conditions to the female nervous system shown in B. (D, D’) Expression pattern of SS39313 (pC1eSS2). This line shows expression in males in a cell type clearly distinct from pC1e (not shown). (E, E’) Expression pattern of SS59433 (pC1eSS3).

**Figure 6 – figure supplement 2.**
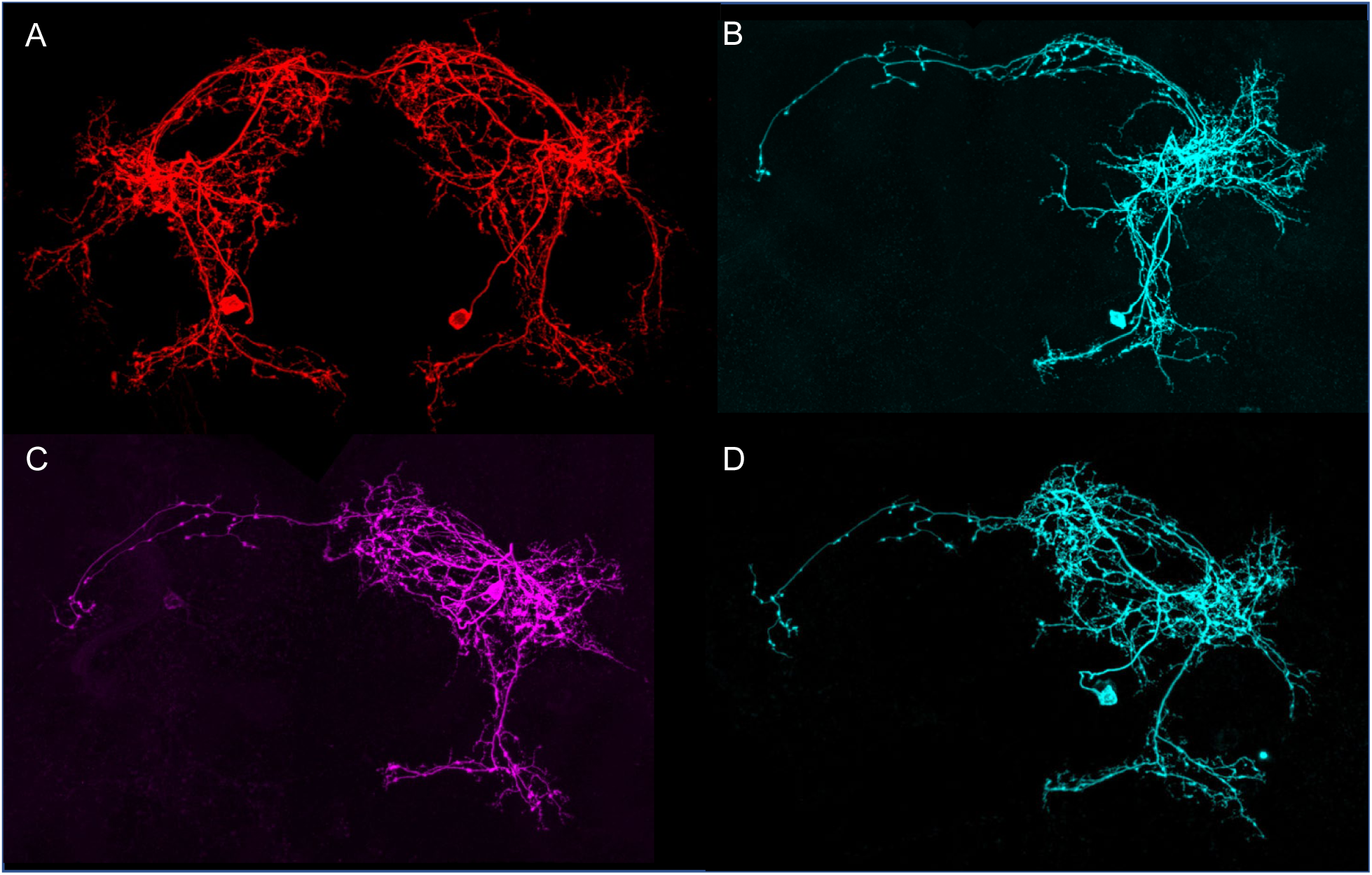
Morphologies of individual pC1e neurons. The images in all panels were generated by stochastic labeling of pC1e split-GAL4 lines using the MultiColor FlipOut method (Nern et al. 2015). (A, B) Images from SS59336 (pC1eSS1). (A) The right and left hemisphere pC1e cells are shown. (C, D) Images from SS43274 (pC1dSS3). Note this line contains both pC1d and pC1e cells. See Figure 5 – figure supplement 2 for pC1d cells.

**Figure 6 – figure supplement 3.**
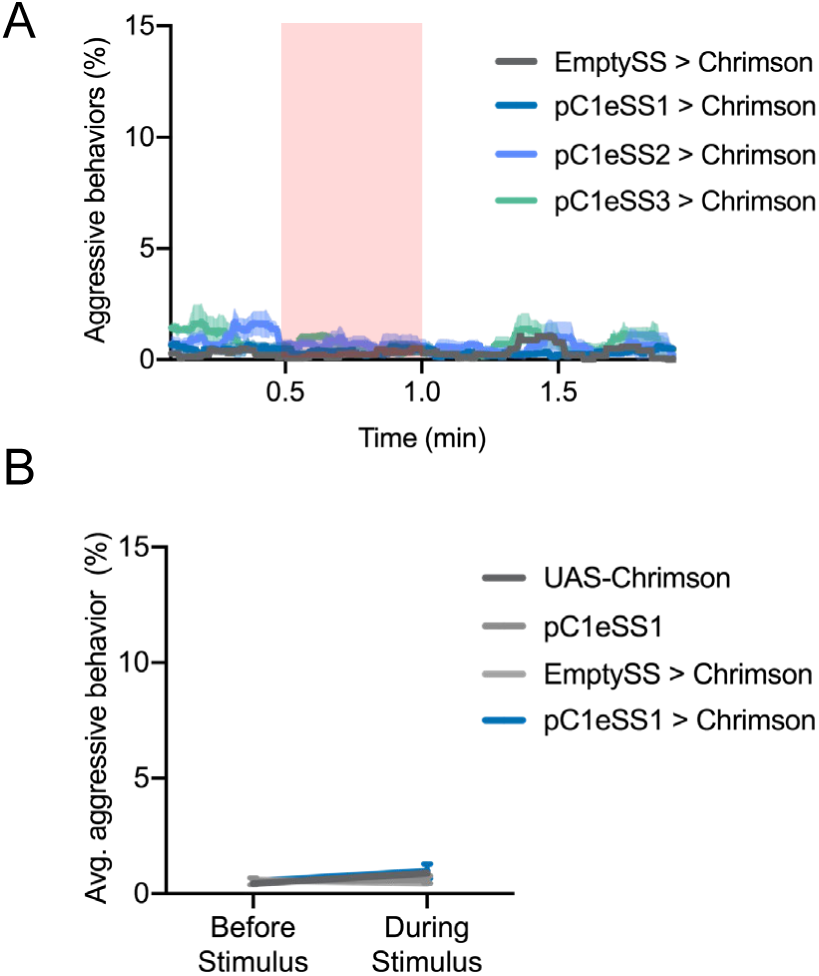
Optogenetic activation of additional lines labeling pC1e. (A) Percentage of flies engaging in aggressive behaviors over the course of a 2-minute trial during which a 30-second 0.1 mW/mm^2^ continuous light stimulus was delivered. Red shading indicates the stimulus period. The mean is represented as a solid line and shaded error bars represent variation between experiments. Each experiment included approximately 15 flies. EmptySS > Chrimson, n = 1 experiment; pC1eSS1 > Chrimson, n = 3 experiments; pC1eSS2 > Chrimson, n = 3 experiments; pC1eSS3 > Chrimson, n = 3 experiments. (B) Average percentage of flies engaging in aggressive behaviors over the 30 second period prior to or during the delivery of a 0.1 mW/mm^2^ stimulus. Each experiment included approximately 15 flies. 20xUAS-CsChrimson, n = 3 experiments; pC1eSS1, n = 2 experiments; EmptySS > 20xUAS-CsChrimson, n = 3 experiments; pC1eSS1 > 20xUAS-CsChrimson, n = 3 experiments. Bars are mean +/- S.E.M.

**Figure 6 – figure supplement 4.**
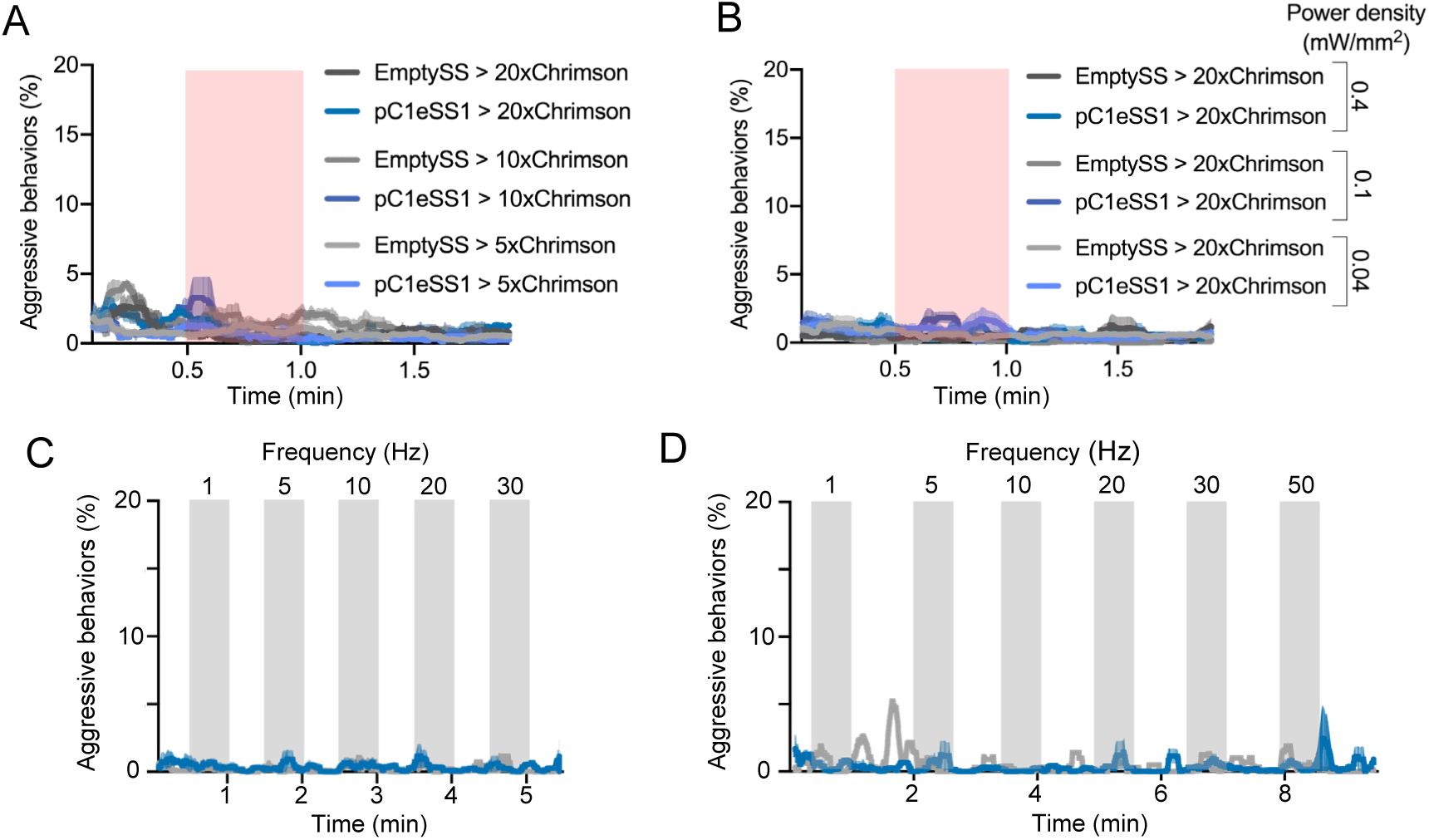
Behavioral effects of stimulus delivery and effector copy number. (A) Percentage of flies engaging in aggressive behaviors over the course of a 2-minute trial during which a 30-second 0.1 mW/mm^2^ continuous light stimulus was delivered to flies carrying either 5xUAS, 10XUAS or 20xUAS effectors. Each experiment included approximately 15 flies. EmptySS > 20xUAS-Chrimson, n = 2 experiments; pC1eSS1 > 20xUAS-Chrimson, n = 2 experiments; EmptySS > 10xUAS-Chrimson, n = 3 experiments; pC1eSS1 > 10xUAS-Chrimson, n = 2 experiments; EmptySS > 5xUAS-Chrimson, n = 6 experiments; pC1eSS1 > 5xUAS-Chrimson, n = 2 experiments. (B) Percentage of flies engaging in aggressive behaviors over the course of a 2-minute trial during which a 30-second 0.4, 0.1, or 0.04 mW/mm^2^ continuous light stimulus was delivered. Each experiment included approximately 15 flies. 0.04 mW/mm^2^: EmptySS > Chrimson, n = 5 experiments; pC1eSS1 > Chrimson, n = 5 experiments; 0.1 mW/mm^2^: EmptySS > Chrimson, n = 4 experiments; pC1eSS1 > Chrimson, n = 3 experiments; 0.4 mW/mm^2^: EmptySS > Chrimson, n = 3 experiments; pC1eSS1 > Chrimson, n = 3 experiments. (C – D) Blocks of 30-second photostimulation (grey bars) with increasing stimulation frequency separated by 30 (C) or 60 (D) second intervals were delivered sequentially to females. Light was delivered at 0.1mW/mm^2^ with a 10 ms pulse width and increasing frequencies resulting in higher numbers of pulses given over the 30-second stimulus on period. The pulse period and pulse number during each period was as follows: 1000ms, 30; 200ms, 150; 100ms, 300; 50ms, 600; 33ms, 909; 20ms, 1500. Each experiment included approximately 15 flies. A: EmptySS > 20xUAS-Chrimson, n = 3 experiments; pC1eSS1 > 20xUAS-Chrimson, n = 3 experiments; B: EmptySS1 > 20xUAS-Chrimson, n = 1 experiment; pC1eSS1 > 20xUAS-Chrimson, n = 2 experiments.

**Figure 6 – figure supplement 5.**
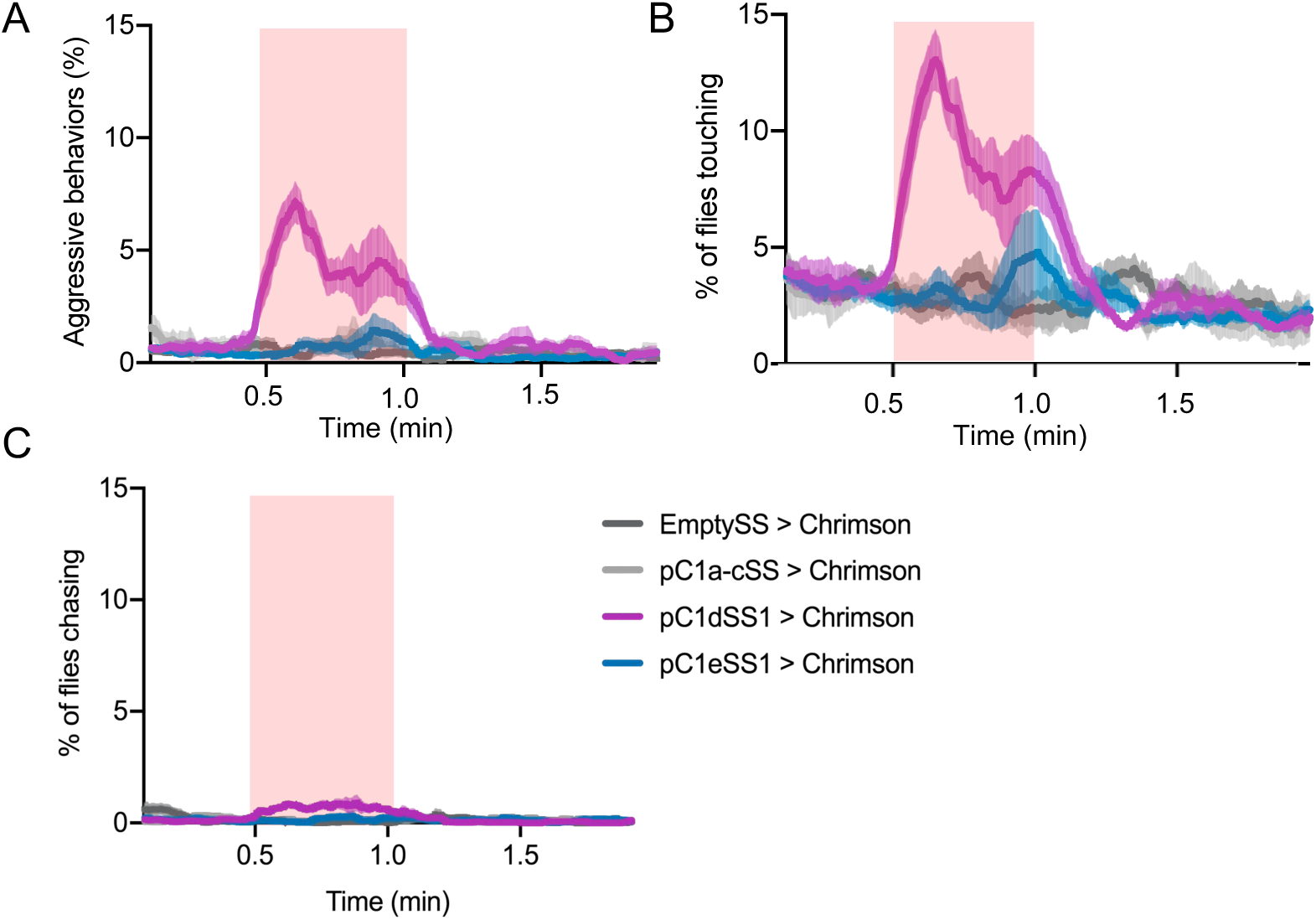
Comparison of activation phenotypes of pC1d, pC1e and pC1a-c. (A-C) Percentage of female flies engaged in aggressive behaviors (A) touching (B) or chasing (C) over the course of a two-minute trial during which a 30-second 0.1 mW/mm^2^ continuous light stimulus was delivered (red shading). Each experiment included approximately 15 flies. EmptySS > 20xUAS-Chrimson, n = 4 experiments; pC1a-cSS > 20xUAS-Chrimson, n = 4 experiments; pC1dSS1 > 20xUAS-Chrimson, n = 6 experiments; pC1eSS1 > 20xUAS-Chrimson, n = 6 experiments. The pC1a – c line was provided by K. Wang and Barry Dickson.

**Figure 7 – figure supplement 1.**
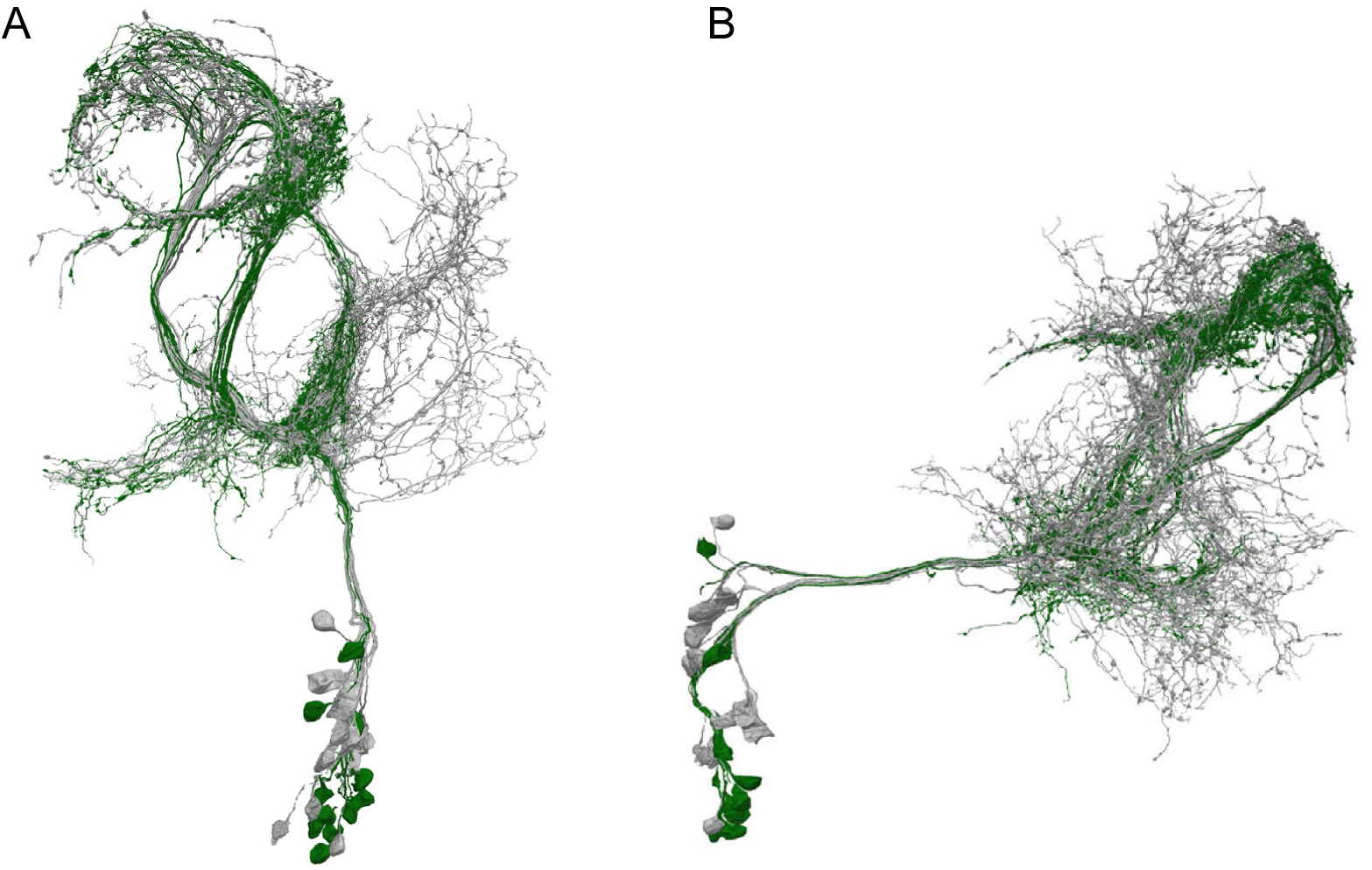
Identification of aIPg neurons in the hemibrain dataset. (A – B) Skeleton renderings of 26 putative aIP-g neurons described in Cachero et al. (2010) identified in the hemibrain dataset. The 11 aIPg type 1 – 3 neurons are displayed in green with rest of the putative aIPg neurons shown in grey. Note that the aIP-g cluster described in Cachero et al. (2010) included neurons with cell bodies in the anterior of the brain, none of which were traced for this project in the FAFB dataset or included in this grouping the hemibrain as the neurons in our split-GAL4 lines only contained posterior located cell bodies.

**Figure 8 – figure supplement 1.**
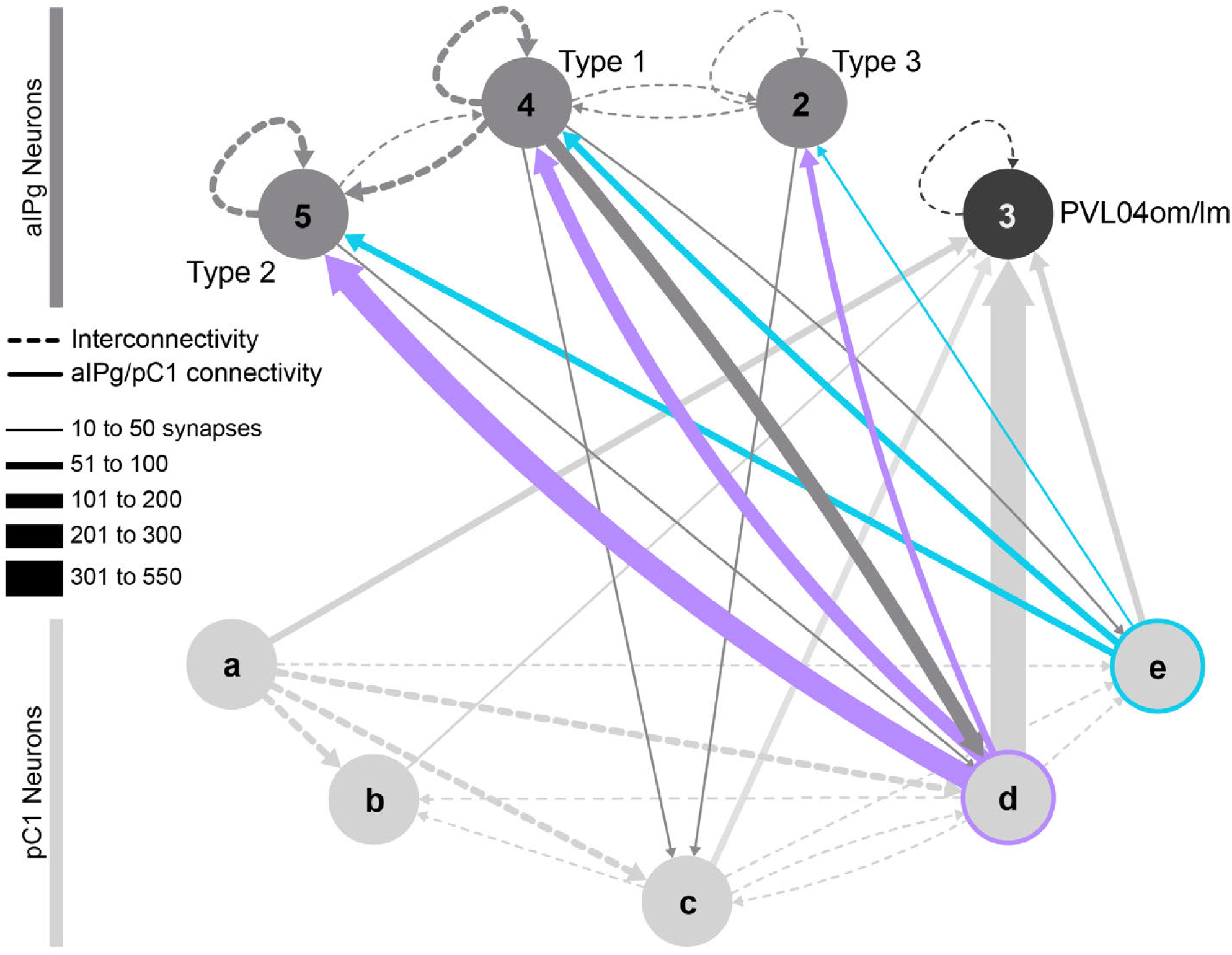
Connections between pC1a-c neurons and pC1d and pC1e. pC1a, pC1b and pC1c neurons have been added to the diagram shown in Figure 8. pC1a and pC1c provide weak input to pC1d, and all pC1 neurons provide input to the three PVL04om/lm neurons, which are among pC1d’s strongest downstream targets. Connections were thresholded at 10 synapses.

**Figure 11 – figure supplement 1.**
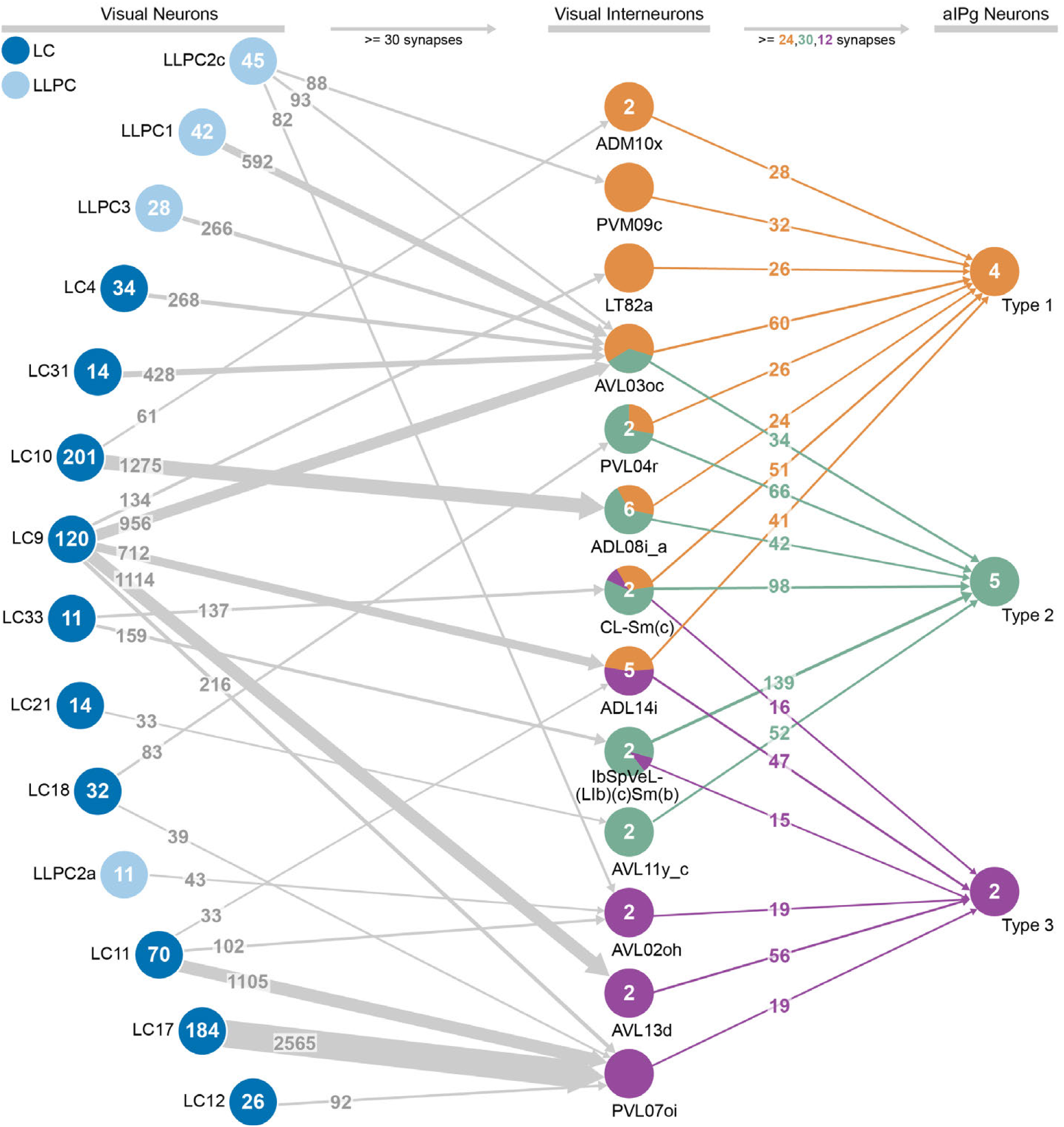
Connections between visual projection neurons, interneurons, and aIPg type 1, type 2 and type 3. Connections were thresholded at 30 combined synapses for each population of visual projection neurons to interneurons. Inputs to the aIPg neurons were thresholded at 6 synapses per cell, resulting in collective thresholds of 24, 30, and 12 synapses to type 1, type 2, and type 3, respectively.

---

**Supplementary Table 1. Framewise performance of automated classifiers.**

**Supplementary Table 2. List of the top pre-synaptic inputs and post-synaptic outputs of the 11 aIPg neurons identified.** Thresholded at 57 synapses for the inputs and 150 synapses for the outputs.

**Supplementary Table 3. List of the top pre-synaptic inputs and post-synaptic outputs of pC1d.** Thresholded at 90 synapses for the inputs and 95 synapses for the outputs.

**Supplementary File 4. Genotypes of the split-GAL4 lines used and cell counts for each of the aIPg split-GAL4 lines.**

**Supplementary File 5. Sample size and statistics for behavioral analysis.**

**Video 1. Movie of control (EmptySS > Chrimson) flies with 0.1 mW/mm*^2^* light density stimulation.**

**Video 2. Movie of stimulating flies from line labelling aIPg1-3 (aIPgSS1 > Chrimson) with 0.1 mW/mm*^2^* light density.**

**Video 3. High speed video of two aIPgSS1>Chrimson female flies.**

**Video 4. High speed video of an aIPgSS1>Chrimson female fly with a Wt (Canton-S) male.**

**Video 5. Movie of stimulating flies from line labelling pC1d (pC1dSS1 > Chrimson) with 0.1 mW/mm*^2^* light density.**

**Video 6. High speed video of two pC1dSS1>Chrimson female flies.**

**Video 7. Movie of the location of pre-and post-synaptic connections with aIPg1-3 neurons based on the EM dataset.** Skeleton renderings and synapse locations shown were taken from NeuPrint and NeuTu.

**Video 8. Movie of aIPg types 1 – 3 and their interconnectivity based on the EM dataset.** The number of neurons within the cell types depicted are in the circles and synapse number for each connection is displayed next to the arrow in the cartoon.

**Video 9. Movie of aIPg types 1 – 3 connectivity with pC1d and pC1e based on the EM dataset.**

**Video 10. Movie of the top pre-synaptic inputs to aIPg types 1 – 3 based on the EM dataset.**

**Video 11. Movie of the top post-synaptic outputs of aIPg types 1 – 3 based on the EM dataset.**

**Video 12. Movie of the top pre-synaptic inputs to pC1d based on the EM dataset.**

**Video 13. Movie of the top post-synaptic outputs of pC1d based on the EM dataset.**

